# Revealing grand-paternal programming of lipid metabolism using a novel computational tool

**DOI:** 10.1101/2020.06.23.166819

**Authors:** Samuel Furse, Adam J. Watkins, Davide Chiarugi, Nima Hojat, James Smith, Huw E. L. Williams, Albert Koulman

## Abstract

While the consequences of poor maternal diet on the offspring’s cardio-metabolic health have been studied in detail, the role of the father’s diet on the health of his offspring is poorly understood. We used a known mouse model to establish the impact of an isocaloric paternal low-protein high-carbohydrate diet on the offspring’s lipid metabolism. Detailed lipid profiles were acquired from F1 neonate (3 weeks), F1 adult (16 weeks) and F2 neonate male and female offspring, in serum, liver, brain, heart and abdominal adipose tissues by Mass Spectrometry and Nuclear Magnetic Resonance. Using a purpose-built computational tool for analysing lipid metabolism as a network, we characterised the number, type and abundance of lipid variables in and between tissues (Lipid Traffic Analysis), finding a variety of alterations associated with paternal diet. These elucidate a mechanism for the defective physiological behaviour of systems at risk of cardio-metabolic disease.

## Introduction

Beyond the serious risk to their metabolic health, obesity in both men and women has long-term consequences for their offspring through nutritional programming^1-4^. There is increasing evidence showing that the nutritional programming of offspring occurs through changes in lipid metabolism^5, 6^ and leads to increased risk of cardio-metabolic diseases (CMD)^7-13^. One contributor to obesity is excess carbohydrate intake. Specifically, high carbohydrate diets have been associated with the emergence of CMD^14-16^ and lower carbohydrate intake with improved recovery^17-19^. One possible explanation is that nutritional programming represents an adaptation to an unbalanced dietary intake in which there is an excess of non-essential nutrients and a deficiency of essential nutrients. However, the effects of a high carbohydrate diet on programming lipid metabolism are not understood. This led us to the hypothesis that a low-protein-high-carbohydrate (LP-HC) diet would alter programming of lipid metabolism in offspring.

This hypothesis was tested by feeding an isocaloric, non-obesogenic LP-HC diet was fed to the (grand)sires of the experimental groups (mouse model^3, 4, 20, 21^). This was designed to increase *de novo* lipogenesis; a high-fat diet would be less useful as it would alter lipid intake as well as biosynthesis. Although the programming effects on lipogenesis were expected to be focused on the offsprings’ liver, the products of lipid biosynthesis are typically distributed throughout the organism quickly, especially triglycerides (TG)^22^. Testing this hypothesis therefore also required a tool for analysing systemic lipid metabolism and distribution.

However, most computational tools developed to study metabolism are focused on one compartment, not to analyse networks and none for lipid traffic^23, 24^. Analysis of single tissues does not provide a complete picture of systemic programming effects. Furthermore, most of the current tools pivot on substrate-enzyme-product relationships to allow for direct linkage to genes and proteins, rather than the local function of metabolites, making it impossible to characterise a whole system. Equally, lipid metabolism is distinct from amino acid and nucleotide metabolism; lipids are not polymers, vary greatly in structure and comprise components from unconnected sources. A network analysis tool was therefore designed to characterise the number, type and abundance of lipids in and between tissues, referred to as lipid traffic.

The novel lipid computational tool reported here, Lipid Traffic Analysis, was used to analyse lipidomics data from liver, serum, brain, heart and adipose tissues (*Fig. 1A*). The connections between the tissues represented the major lipid ‘highways’ in the organism (*Fig. 1B*). Lipid Traffic Analysis identified altered lipid metabolism through a Switch Analysis (which lipids were present and where) and an Abundance Analysis (quantitative differences between phenotypes). Importantly, these analyses represent the state-of-the-art in characterisation of lipid metabolism across organs.

**Fig. 1.**
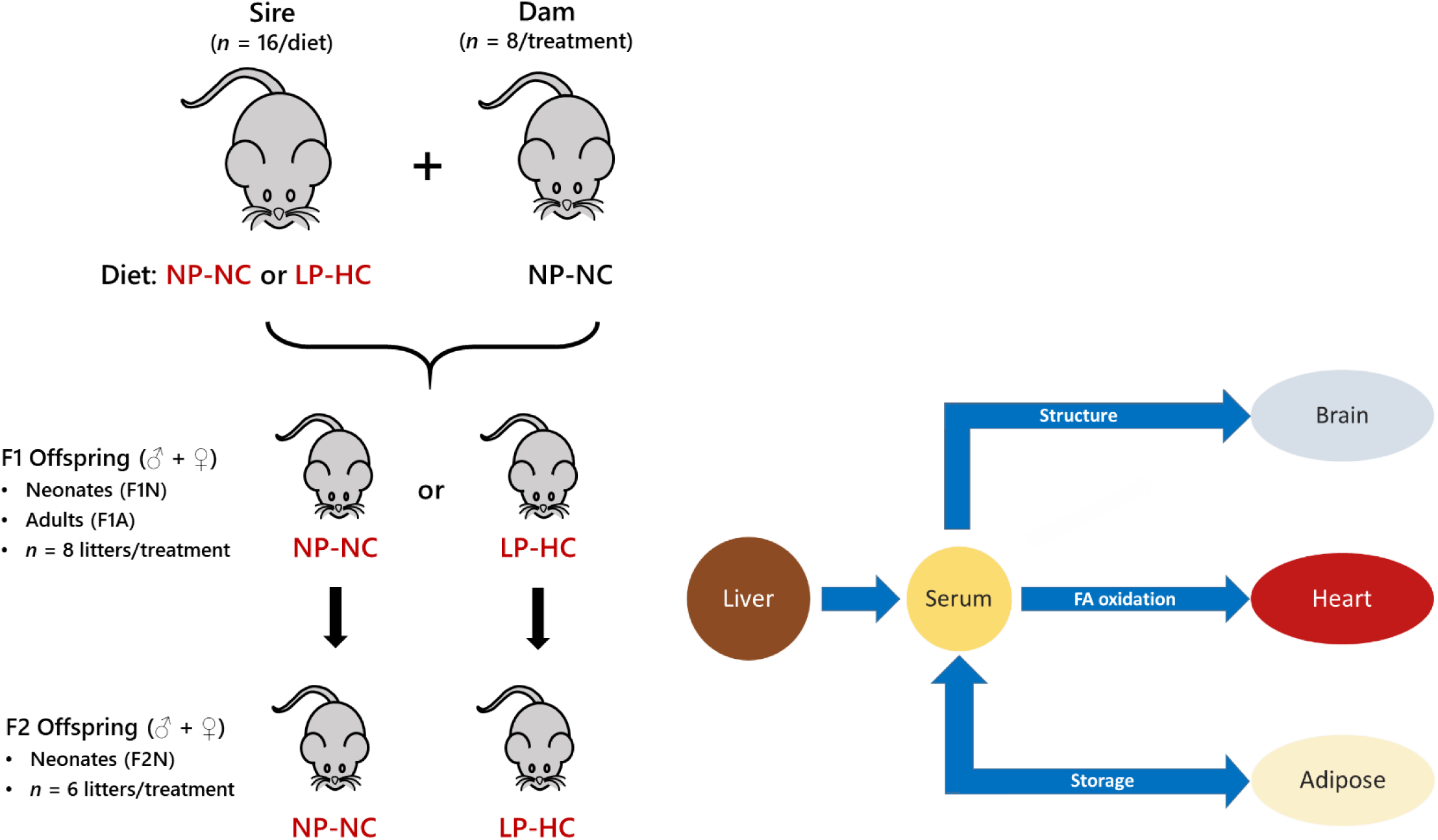
The mouse model and tissues used for lipid traffic analysis associated with de novo lipogenesis. Panel A, Schematic representation of the mouse model showing the generation of programmed offspring across two generations. Panel B, the network that describes the lipid traffic associated with de novo lipogenesis from the liver to termini (CNS, heart and adipose) via the serum. The termini represent traffic flow for structural purposes (CNS), fatty acid oxidation (heart) and storage (adipose). This metabolic relationship between tissues was used as the structure of the network for all analyses in the present study. NP-NC refers to a diet of normal protein-normal carbohydrate where LP-HC refers to a low protein-high carbohydrate diet. The NP-NC and LP-HC are the same as NN and LL groups used in earlier studies^3, 20^. Adipose was only available for F1A groups, whole brain samples used for F1N groups, with separate right brain and cerebellum for F1A and F2N.

We wanted to test the hypothesis that a higher carbohydrate intake in (grand)sires alters lipid metabolism in offspring as this contributes to our understanding of dietary intake and metabolic programming^25, 26^ and the effects of metabolic disease across generations^5^ in a model system. This gives us an insight into possible interventions to improve human metabolic health in familial circumstances.

## Results

A combination of Direct Infusion Mass Spectrometry (DI-MS^27-29^) and phosphorus nuclear magnetic resonance (^31^P NMR^30, 31^), known as dual spectroscopy^29^, was used to identify and verify the abundance of lipid classes between the two ionisation modes respectively (*Fig. S1*, NMR data for each compartment shown in *Supplementary Information*). This study identified up to 586 lipid variables in positive ionisation mode and up to 564 lipid variables in negative ionisation mode in liver, brain, heart and adipose homogenates and in serum.

### Lipid Traffic Analysis: Design of a novel computational tool for the network analysis of lipid metabolism

The first stage in developing this computational tool was to categorise lipid variables according to where they were found with respect to adjacent lipid compartments in the biological network (*Fig. 1*). Some lipid variables were found in *all* compartments, others in two adjacent compartments and others in one compartment only (*Fig. 2A*). These we refer to as ***A, B*** and ***U*** type lipids (or categories), respectively. Novel code written in R for identifying such lipids is described in Methods and can be found in the Supplementary Information. The basis of these categories was that they represented the intersections between lipid compartments, *i*.*e*. stations in the network. Distinct patterns of the presence of lipid species that appear in adjacent compartments or ubiquitously can represent systemic responses. Different axes between organs (*e*.*g*. Liver-Serum, Serum-Heart) can be considered and physiological metabolic functions compared, that correspond to the fate or *traffic* of shared lipids. This relationship was measured in the present study using unlabelled species as an average over longer periods, *e*.*g*. stage of development. These categorisations of the lipids are then used to address lipid traffic analysis from two different perspectives (*Fig. 2B*), namely a quantitative Abundance analysis and a binary Switch Analysis. Both of these represent novel analyses presented in this work.

**Fig. 2.**
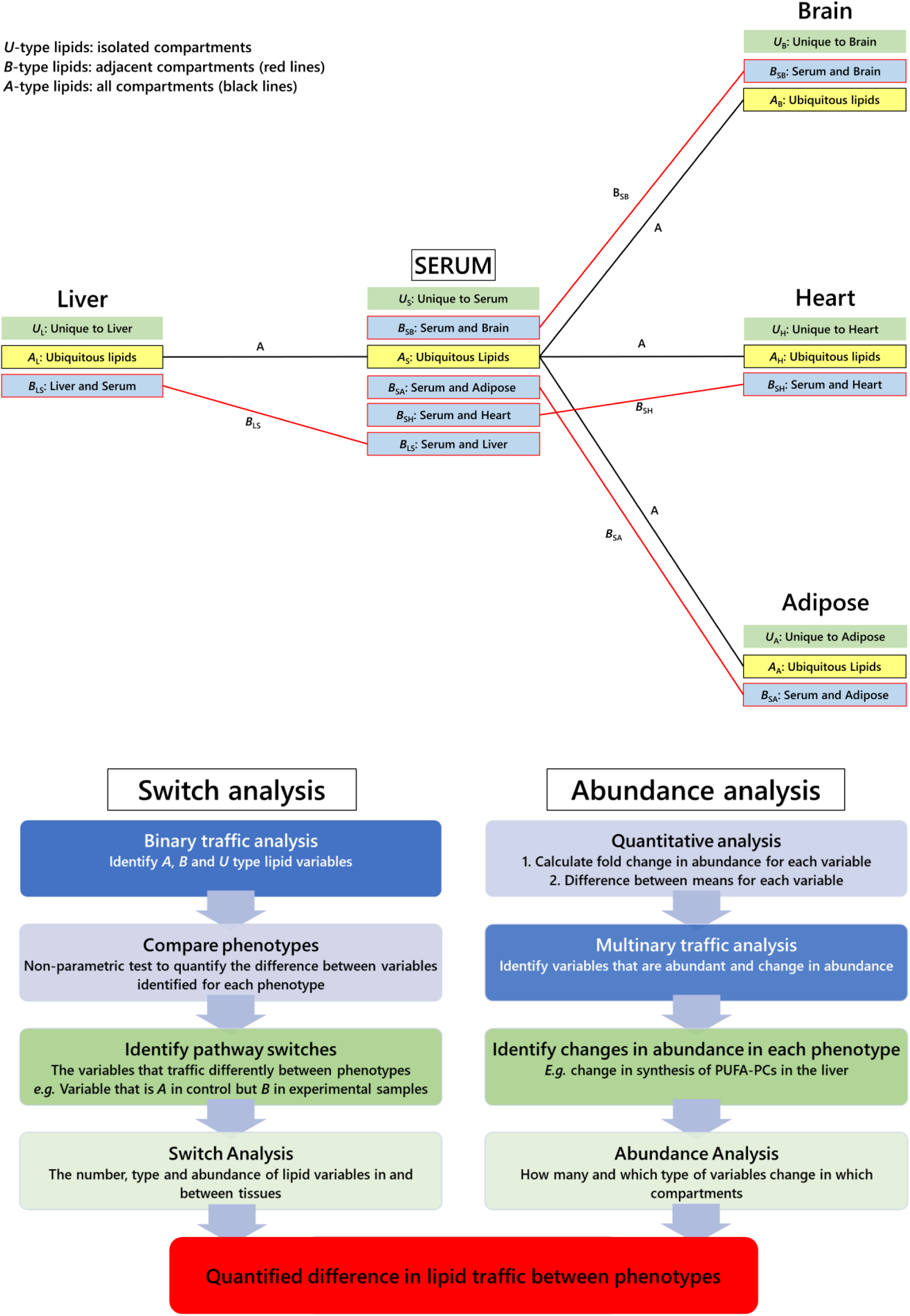
Structure of Traffic Analysis for quantifying changes in lipid metabolism. Panel A, categorisation of lipids according to where they are found; B, flow chart of Traffic Analysis showing the gross structure of the analysis. **A, B** and **U** are categories representing variables that appear in all compartments, in pairs of adjacent compartments and in only one compartment, respectively. Subscripts to these categories are pairs of one-letter codes indicating the direction of the traffic (reading left to right). Red connections show B-type lipid connections. Black connections show A-type lipid connections. The two strands of the flow chart represent separate analyses that use the same R code (see SI). Equations for the Quantitative Analysis are shown in Eq.1 and Eq. 2.

### Switch Analysis

The Switch Analysis developed in the present study identified the lipids that are above and below the limit of detection in a given compartment (***U***), between adjacent compartments (***B*** type lipids, *e*.*g*. Liver-Serum axis) or ubiquitously (***A*** type lipids). This therefore represented lipids that were switched on or off with respect to a measurement threshold. The Switch Analysis requires only straightforward lipidomics data rendered as relative abundance. The Switch Analysis of TGs in the F1N group is shown in *Fig. 3A*. Pie charts show the number of TG variables in each phenotype. Jaccard-Tanimoto coefficients (JTC, with accompanying *p*-values) were used to characterise the similarity between the compared groups. The JTC coefficient indicated the proportion of variables that appeared in the two groups^32, 33^ whereas the *p*-value indicated what differentiated them; a *p* < 0·5 indicated there were variables unique to both groups, whereas *p* > 0·5 meant that only one group could have any unique variables. Thus, a JTC of 0.67 with a *p*-value of 1 indicated that two thirds of variables appear in both groups and that the other third of the variables only appeared in one group.

**Fig. 3.**
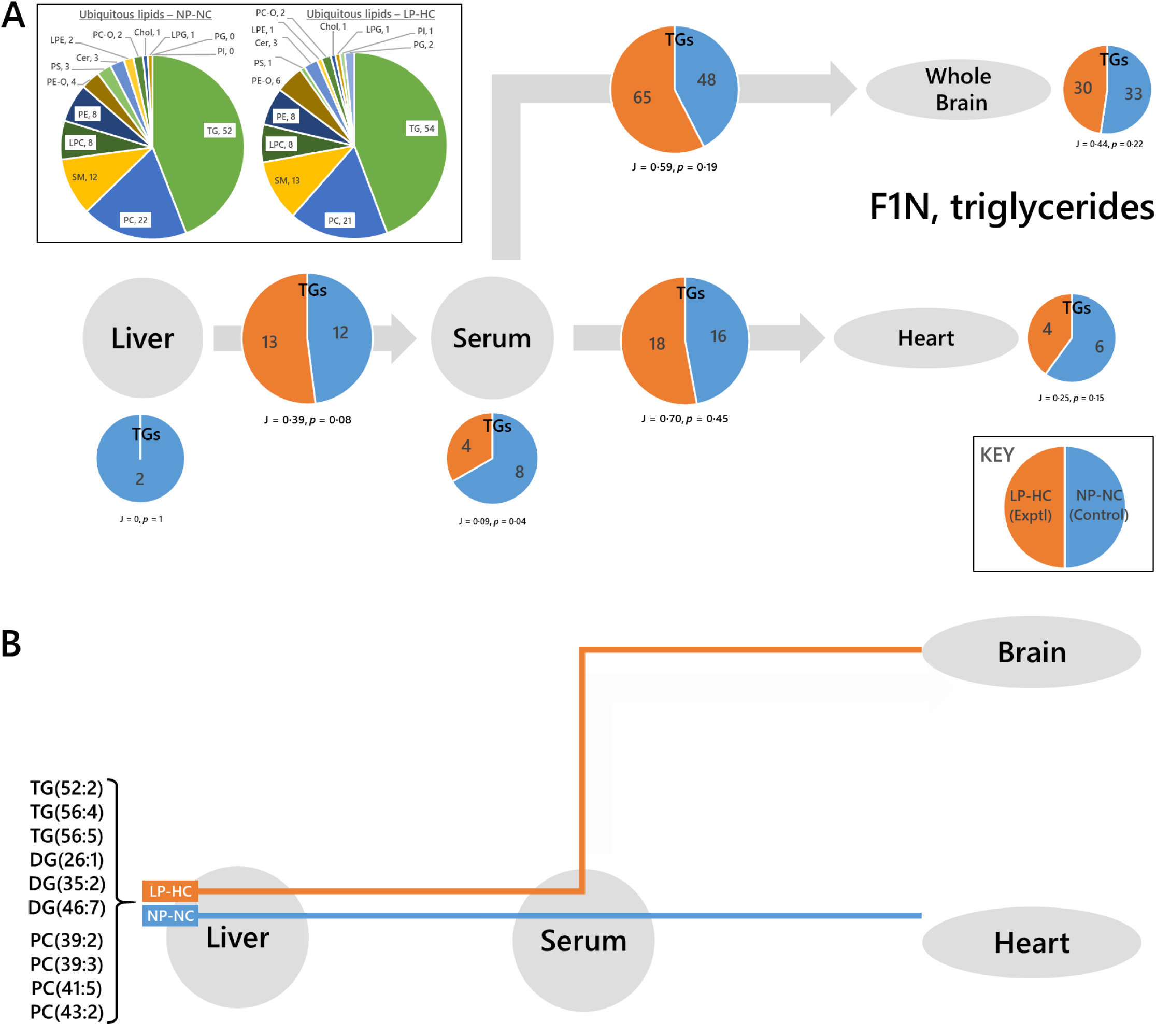
Switch analyses of triglyceride variables in the neonatal F1 offspring (F1N) of fathers fed a normal (NP-NC) or a low-protein, high carbohydrate (LP-HC) diet, measured by mass spectrometry in positive ionisation mode. Panel A, Traffic analysis of triglyceride variables; B, Switch analysis of triglyceride (measured in positive ionisation mode) and phosphatidylcholine (measured in negative ionisation mode) variables in F1 neonatal mice re-routed from the Serum-Heart in the control group to the Serum-Brain in the experimental group. TG and PC variables on the Serum-Heart axis in the control (NP-NC) group of F1Ns that are found on the Serum-Brain axis of the experimental (LP-HC) group of F1Ns but not their Serum-Heart axis. The pie charts in the insert show the number of ubiquitous lipid variables for that network, for each phenotype. Larger pie charts with J values represent lipids found in two adjacent compartments. Smaller pie charts with J values represent lipids found in isolated in compartments. J represents the Jaccard-Tanimoto coefficient for the comparison, with accompanying p-value, as a measure of the similarity between the variables identified in the two phenotypes for each comparison. The p-value shown represents the probability that the difference between the lists of variables for the two phenotypes occurred by random chance. Cer, ceramide; Chol, cholesterol; DG, diglyceride (water-loss adduct from fragmentation in source); LPC, lyso-phosphatidylcholine; LPE, lyso-phosphatidylethanolamine; LPG lyso-phosphatidylglycerol; PA, phosphatidic acid; PC, phosphatidylcholine; PC-O, phosphatidylcholine plasmalogen; PE, phosphatidylethanolamine; PE-O, phosphatidylethanolamine plasmalogen; PG, phosphatidylglycerol; PI, phosphatidylinositol; PS, phosphatidylserine; SM, sphingomyelin; TG, triglyceride.

We began by investigating TGs, as several of these are well established markers of *de novo* lipogenesis (DNL) and thus easily affected by changes in lipogenesis. Major TGs on the Liver-Serum axis were also found on the Serum-Brain and Serum-Heart axes in F1N NP-NC (control) mice. This led us to ask whether any of those variables were routed differently according to phenotype. Specifically, it was observed that 17 more TG variables appeared on the Serum-Brain axis in F1N of the LP-HC group than the NP-NC group (*Fig. 3A*). We therefore tested the hypothesis that variables found in the NP-NC (control) group Serum-Heart axis were also found in the Serum-Brain axis of the LP-HC group, without also appearing in the Serum-Brain axis of the NP-NC group. Six such variables were found to be on the Serum-Brain axis of the experimental group (LP-HC), implying that they were re-routed in this phenotype (*Fig. 3B*).

In the F1 Adults, for which we could include adipose tissue in the network, we found that all of the TG variables that appeared on the Liv-Ser axis in F1A NP-NC also appeared on one or more of the Serum-Cerebellum, Serum-Right Brain, Serum-Heart or Serum-Adipose axes for the NP-NC group (*Extended Data Table 1*).

The larger number of variables found on the Serum-Adipose axis in F1A adult NP-NC mice, and the larger number of TG variables found on the Serum-Cerebellum and Serum-Right Brain axes suggested to us that TGs were re-routed from the adipose to the CNS in F1 adults due to the low protein, high carbohydrate paternal diet(*Fig. 4A*). This was true for at least two variables, DG(33:1) and TG(52:5) (*Extended Data Table 2*, columns 3-5). However, this left several variables unaccounted for. Furthermore, there appeared to be a difference in the number of variables in the NP-NC and LP-HC groups on the Liv-Ser axis (*Fig. 4A*). We therefore tested the hypothesis that a portion of the variables in the network were associated with this difference. The seven variables that distinguished NP-NC from LP-HC on the Liver-Serum axis of F1A (shown in pale green cells in *Extended data table 2*) were all either found in the Serum-Cerebellum/Right Brain or Serum-Heart axes of the LP-HC phenotype, but not the NP-NC phenotype.

**Fig. 4.**
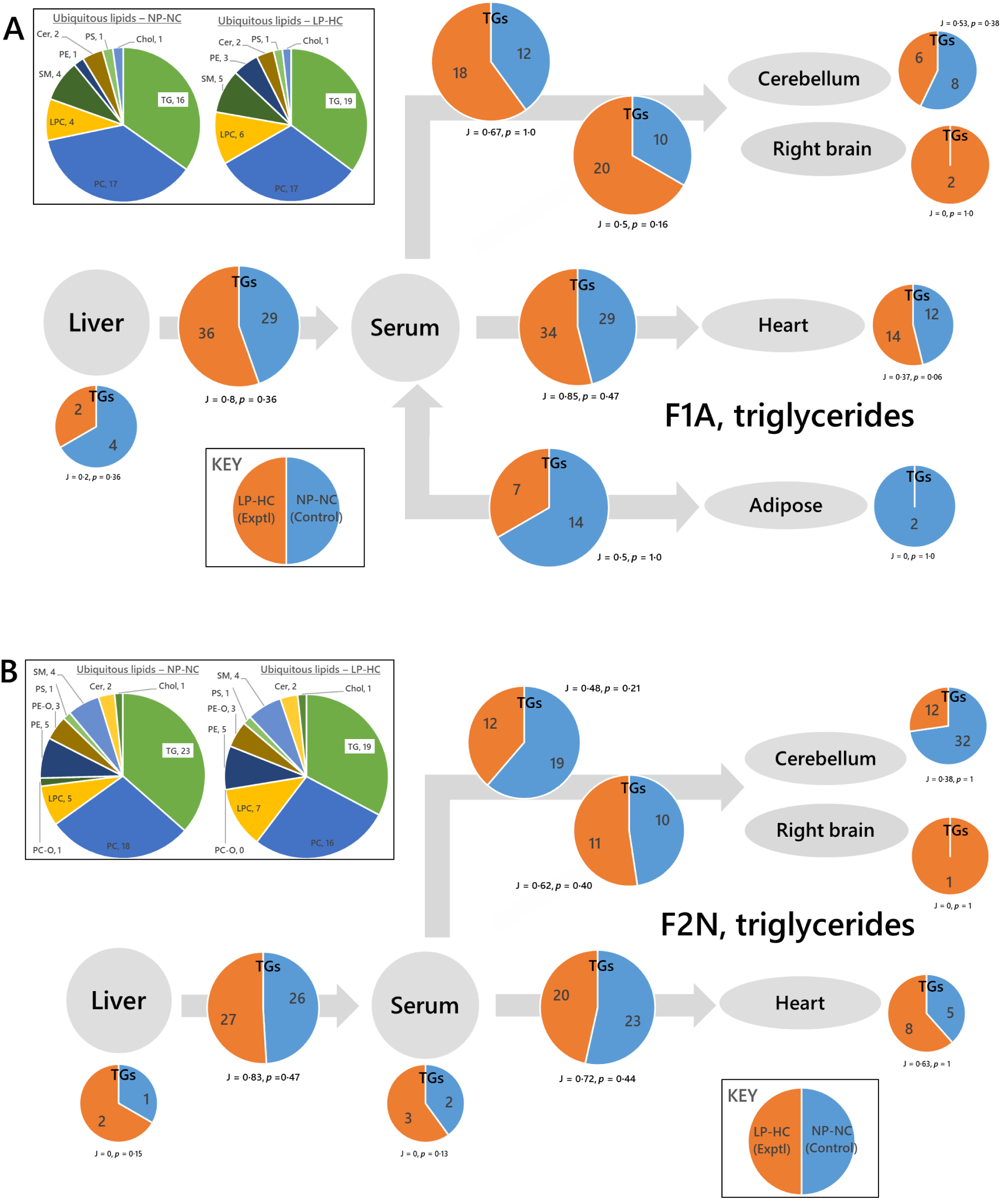
Switch analyses of triglyceride variables in the adult F1 (F1A) and neonatal F2 (F2N) offspring of fathers fed a normal (NP-NC) or a low-protein, high carbohydrate (LP-HC) diet, measured by mass spectrometry in positive ionisation mode. Panel A, F1 Adults (F1A); B, F2 Neonates (F2N). The pie charts in the insert show the number of ubiquitous lipid variables for that network, for each phenotype. Larger pie charts with J values represent triglyceride variables found in two adjacent compartments. Smaller pie charts with J values represent triglyceride variables found in isolated in compartments. J represents the Jaccard-Tanimoto coefficient for the comparison, with accompanying p-value, as a measure of the similarity between the variables identified in the two phenotypes for each comparison. The p-value shown represents the probability that the difference between the lists of variables for the two phenotypes occurred by random chance. Cer, ceramide; Chol, cholesterol; DG, diglyceride (water-loss adduct from fragmentation in source); LPC, lyso-phosphatidylcholine; LPE, lyso-phosphatidylethanolamine; LPG lyso-phosphatidylglycerol; PA, phosphatidic acid; PC, phosphatidylcholine; PC-O, phosphatidylcholine plasmalogen; PE, phosphatidylethanolamine; PE-O, phosphatidylethanolamine plasmalogen; PG, phosphatidylglycerol; PI, phosphatidylinositol; PS, phosphatidylserine; SM, sphingomyelin; TG, triglyceride.

Trafficking of TGs was also investigated in F2N individuals (*Fig. 4B*). This analysis showed that there were a considerable number of variables unique to the Cerebellum in the NP-NC group and not present in the LP-HC group. The network analysis used in this study showed that these variables were not found elsewhere in the system. This is remarkable because this tissue does not typically use FA for energy, raising the questions of why they are in the cerebellum and why they are not in the LP-HC phenotype. 14 of the 20 TGs comprise FAs with an odd number of carbons in the chain (*Extended Data Table 3*). This result forms part of a wider characterisation of the relationship between fats and the CNS, with re-routing of TGs to the CNS from the heart in the LP-HC phenotype (*Figs 3 and 4*). A close or complicated relationship of the CNS with energy supply by TGs is counter-intuitive because the principle carbon source of the CNS is glucose and not fat, and even under starvation conditions, only a small proportion of the ATP used in the CNS is made from energy released from primary metabolism of fats.

As the traffic of TGs differed between phenotypes, the hypothesis that phosphatidylcholine (PC) traffic was associated with a LP-HC diet in (grand)sires across F1N, F1A and F2N groups was also tested. The results of network analysis in F1N indicated the possibility of a re-routing of PC variables according to phenotype (*Extended Data Fig. 1A*). Specifically, more PC variables were found on the Serum-Brain axis in the LP-HC group, where the opposite was the case for the Serum-Heart axis. There were about four variables that distinguished the NP-NC Serum-Heart axis and the LP-HC axis, all of which also distinguished the Serum-Brain axis in the LP-HC group from the NP-NC (*Fig. 3B*).

In the F1A network, 8 PCs were found on the Serum-Heart axis in the NP-NC group that were not found on the same axis of the LP-HC group (*Fig. 5A*), of which two were re-routed to the CNS in the LP-HC (*e*.*g*. PC(43:2), *Fig. 5B*). We found that PC(33:4) and PE(41:4) were only found in the LP-HC phenotype, whereas PC(42:4) was found throughout the NP-NC phenotype but only in the LP-HC liver. This suggested there were several modulations to PC and PE metabolism associated with this phenotype, and is consistent with long-standing evidence that PC is used as a means for storing/transporting polyunsaturated FAs such as arachidonic acid^34^.

**Fig. 5.**
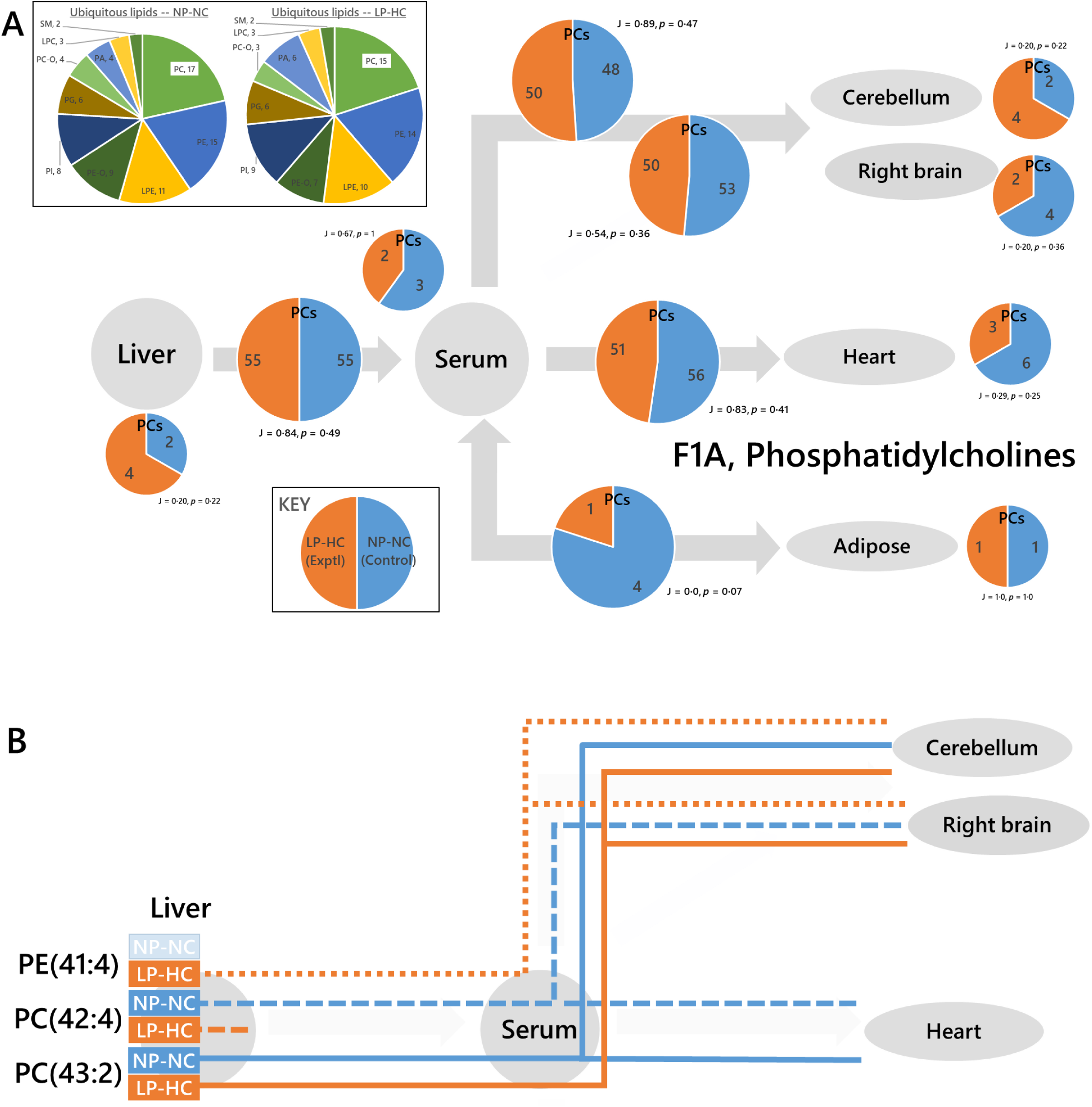
Switch analyses of phospholipid variables in the adult F1 (F1A) offspring of sires fed a normal (NP-NC) or a low-protein, high carbohydrate (LP-HC) diet, measured by mass spectrometry in negative ionisation mode. Panel A, all phosphatidylcholines; B, network diagram showing the distribution of PE(41:4) (dotted line), PC(42:4) (dashed line), PC(43:2) (solid line). The pie charts in the insert show the number of ubiquitous lipid variables for that network, for each phenotype. Larger pie charts with J values represent PC variables found in two adjacent compartments. Smaller pie charts with J values represent PC variables found in isolated in compartments. J represents the Jaccard-Tanimoto coefficient for the comparison, with accompanying p-value, as a measure of the similarity between the variables identified in the two phenotypes for each comparison. The p-value shown represents the probability that the difference between the lists of variables for the two phenotypes occurred by random chance. LPC, lyso-phosphatidylcholine; LPE lyso-phosphatidylethanolamine; PA, phosphatidic acid; PC, phosphatidylcholine; PC-O, phosphatidylcholine plasmalogen; PE, phosphatidylethanolamine; PE-O, phosphatidylethanolamine plasmalogen; PG, phosphatidylglycerol; PI, phosphatidylinositol; PS, phosphatidylserine; SM, sphingomyelin; TG, triglyceride.

PC traffic in the F2N network (*Extended data Fig. 1B*) was characterised by more PC variables being produced in the control (NP-NC) group (*Extended Data Table 4*). Seven additional variables were found on the Liver-Serum axis of the F2N NP-NC group. Several of these, PC(39:3, 40:4, 40:6, 41:4), were also found on the Serum-Cerebellum and Serum-Right Brain axes (*Extended Data Table 4*), suggesting that the grandsires’ dietary balance influenced the biosynthesis and distribution of phosphatidylcholine in F2 offspring.

### Abundance analysis

Perhaps the most striking result from the Switch analysis (*vide supra*) is the association of the grandsire’s diet to the re-routing of lipid variables from the Serum-heart axis to the serum-CNS axes. This was observed particularly clearly with phosphatidylcholines (*Figs 3* and *5*) for F1N and F1A, with a distinct difference in the number of PC variables trafficked to both the heart and CNS in F2N mice (*Extended Data Fig. 1*). This led us to the hypothesis that there would be a change in the abundance of the ubiquitous PCs associated with the phenotype, *i*.*e*. the PCs found in all tissues would have a difference in abundance between phenotypes. We designed a novel Abundance analysis to test this.

Two numerical dimensions were used in the Abundance Analysis. One of these describes the *margin* of the difference in abundance between the two phenotypes (Eq. 1) and the other the *magnitude* of the difference (error normalised fold change, Eq. 2). In these equations 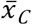 is the mean of values for that variable in the control group (NP-NC), 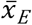 is the mean of values for the experimental group (LP-HC), ‘*a*’ is the standard deviation of the values of the NP-NC group and ‘*b*’ is the standard deviation of the values of the LP-HC group.

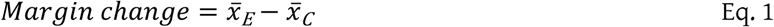

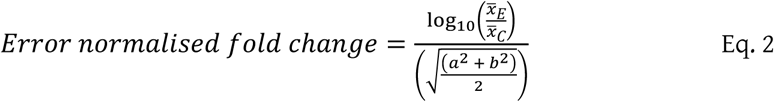

The margin difference is interpreted with a *p*-value calculated using a Student’s *t-*test (see *Methods*), whereas the fold change has a built-in confidence interval through a calculation of the propagated error. The margin changes and accompanying confidence intervals were used to identify the variables that describe the difference in lipid metabolism between phenotypes. The magnitude change was used to quantify this.

The Abundance Analysis found that PE(40:2) and (40:3) were more abundant in the livers of the LP-HC group of F1N mice (*p* = 0·0005 and 0·001, respectively). This was reflected in the abundance pattern in F2Ns, but not F1As (Error normalise fold change, ENFC, plotted in *Fig. 6A*). PE(34:1) was less abundant in the livers of LP-HC F2Ns (*p* = 0·0021) and PE(36:3) less abundant in the serum of LP-HC F2Ns (*p =* 0·0006), however they were in general more abundant in the CNS of LP-HC F2Ns (ENFC plotted in *Fig. 6B*). Two commonplace PC isoforms (38:1 and 38:4) were both less abundant in the CNS of LP-HC F2Ns (*p =* 0·0009 (Cerebellum) and 0·0015 (right brain) respectively), with mixed effects noted for PC(30:0 and 32:2) in the same tissues (*p =* 0·001 and 0·002 respectively, ENFC plotted in *Fig. 6B*). Importantly, 38:1 and 38:4 are commonplace and typical isoforms of PC found in the CNS, as are the PEs. When taken with the Switch Analysis results (*Figs 3* and *5*), this suggests that commonplace isoforms of PC are rerouted away from the CNS and replaced by more recondite ones, *e*.*g*. PC(39:2, 43:2), and PE (*e*.*g*. 36:3), in the offspring of fathers fed a high carbohydrate diet. The higher abundance of PE(34:1) in the circulation of F2N (ENFC = 27·2, *Fig. 6*) and PEs(40:2, 40:3) in the liver of F2N (ENFC = 3·1, 15·8) without a consummate increase in the CNS suggests that despite the difference between phenotypes, PEs(34:1, 40:2, 40:3) are handled differently to PE(36:3) in LP-HC offspring.

**Fig. 6.**
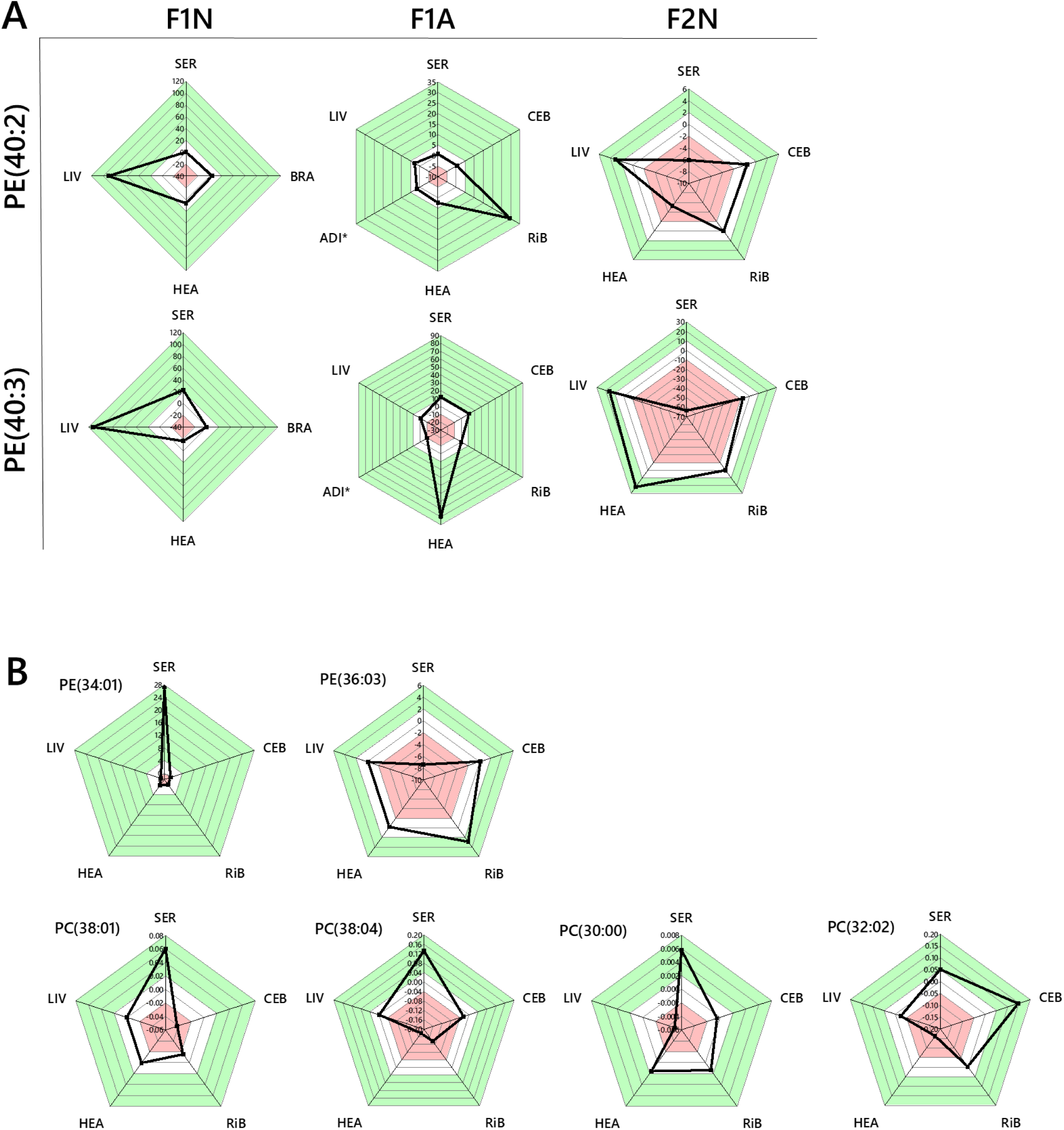
Radar plots of the error normalised fold change in abundance of phosphatidylcholine and phosphatidylethanolamine lipid variables associated with a high carbohydrate dietary intake of (grand)sires. Panel A, PE(40:2, 40:3) were identified as more abundant in F1N livers using statistical approaches, this was followed through all generational groups;. Panel B, PC and PE variables whose abundance in the CNS changes in a manner associated with the dietary phenotype. The white areas represent the 0 point and one division above and below this. The red areas represent values more negative, and the green areas values more positive than this. ADI, adipose; BRA, brain; CEB, cerebellum; HEA, heart; LIV, liver; RiB, right brain; SER, serum. PC, phosphatidylcholine; PE, phosphatidylethanolamine. *Sample treated with petroleum ether to concentrate phospholipid fraction (See Methods).

It is also clear from the Switch analysis that triglycerides are trafficked differently in these two systems (*Figs 3* and *4*), including evidence for TG variables being rerouted (*Fig. 3*). This led us to test the hypothesis that a higher carbohydrate diet consumed by fathers altered *de novo* lipogenesis (DNL) in offspring. We elected to use a targeted approach for testing this, using known markers of *de novo* lipogenesis (DNL)^22^ and reference variables not associated with DNL. The abundance of all DNL TGs was typically much higher in CNS tissue in the LP-HC group (*Extended data Fig. 2A-C*). This was especially clear in F1A individuals (*Extended data Fig. 2B*), where all of the DNL variables were much more abundant in the Right Brain of LP-HC mice. The abundance of a dietary TG and a species made endogenously (TG(54:4) and cholesterol, respectively) were also higher in F1A the CNS. However, reference species not associated with DNL such as phospholipids were not more abundant in the LP-HC phenotype (*Extended data Fig. 2B*), suggesting that the change in lipid traffic is not restricted to DNL species only. These data are consistent with the switch analysis (*Figs 3 and 4*).

## Discussion

This study was motivated by the hypothesis that the ratio of protein and carbohydrate in the paternal grandsire diet influences the lipid metabolism of their offspring. Detailed molecular lipid surveys of several tissues associated with these two phenotypes (NP-NC, control; LP-HC, experimental) were analysed using a novel bioinformatics tool for lipid metabolism. This showed that the number, type and abundance of lipid variables in and between tissues (known collectively as lipid traffic) differed between phenotypes and generations. The intervention was focussed on dietary intake expected to alter *de novo* lipogenesis and thus metabolic activity in the liver. A focused characterisation of the lipid metabolism across two succeeding generations from sires fed in this way revealed that both triglyceride (TG) and phosphatidylcholine (PC) metabolism were altered throughout the network by this dietary intake, and over two generations. However, the changes to lipid traffic and biosynthesis do not appear to be restricted to species associated with DNL.

In particular, the evidence for both TG and phospholipid variables being re-routed to the CNS from the heart and adipose is striking. This shows that both the structural molecules and molecules for the supply of energy (TGs) are associated with LP-HC programming. These results are important because they suggest a molecular mechanism that contributes to the emergence of cardio-metabolic disease. The change in supply of phospholipids is consistent with changes in the physical behaviour of the cardiovascular system that are associated with cardio-metabolic disease. A change in the supply of TGs may also be consistent with pronounced molecular changes in the shift from fatty acid oxidation to glucose metabolism associated with cardiac hypertrophy. Also relevant mechanistically is a larger role for the developing CNS in TG metabolism than is generally understood. Evidence for changes to metabolic control as a result of TGs crossing the blood-brain barrier is already known, through central leptin and insulin receptor resistance^35^. Combined with evidence from the present study, a high carbohydrate diet in fathers may programme their infants for insulin and leptin resistance. Furthermore, there is evidence to link changes to lipid processing in the brain with metabolic disease (review^36^). Thus, the present study shows that the nutritional programming associated with a non-obese phenotype tends towards metabolic disease. This suggests that unbalanced as well as excess nutrition can result in altered metabolism over two generations. This is revealing because it offers a possible mechanism for metabolic disorders in individuals with a healthy adipose volume.

The evidence of shifts in lipid metabolism in the Abundance Analysis (*Fig. 6*) from the present study suggest that the chow diet fed to F1 offspring softens some of the effects of the low protein diet for the F2 generation. Specifically, the pronounced increase in the abundance of PE(40:2, 40:3) in the livers of F1N LP-HC mice is lower in the F2N LP-HCs. Similar patterns are observed for the ENFC of SM(36:1) and TG(48:0) between F1N and F2Ns (*Extended Data Fig. 2*). It is not clear from the present study precisely what causes this, however with appropriate experimental design, markers of programming and re-programming could be identified. Anabolic hormones such as insulin regulate the release and reuptake of lipids from one organ to another, making Traffic Analysis a powerful tool in characterising the change in lipid metabolism and accumulation ^5, 37-42^. Currently, these analyses are often limited to comparisons of blood plasma or serum samples. The new method described here is therefore capable of uncovering new biological meaning in lipid metabolism in timely topics, and relate lipids in different compartments in a way not possible for simple comparisons.

In conclusion, this study has shown that the hypothesis that lipid metabolism is altered in offspring as a result of unbalanced dietary intake by grandsires is correct. The biosynthesis of both TGs and PCs is altered in the liver, with a particular increase in TG traffic reaching the CNS. Furthermore, it is associated with all TGs and not exclusively those associated with DNL. This work shows that a non-obesogenic high carbohydrate, low-protein diet consumed by fathers influences lipid metabolism in offspring over at least two generations. Specifically, the distribution of both triglycerides and phosphatidylcholines is altered in F1 and F2 generations. The network approach to the analysis of lipid metabolism reported here was essential for identifying changes in lipid metabolism that occur across pathways (TG/PL) and with components from different sources (endogenous/dietary), however further work is required to understand how the changes identified can be reversed.

## Methods

### Materials, animals, consumables and chemicals

Purified lipids were purchased from Avanti Polar lipids Inc. (Alabaster, Alabama, US). Solvents and fine chemicals were purchased from SigmaAldrich (Gillingham, Dorset, UK) and not purified further. Mice were purchased from Harlan Laboratories Ltd (Alconbury, Cambridgeshire, UK). Hormones were purchased from Intervet (Milton Keynes, UK).

### Animal model

All procedures were conducted in accordance with the UK Home Office Animal (Scientific Procedures) Act 1986 and local ethics committees at Aston University. Animals were maintained at Aston University’s biomedical research facility as described previously^3^ and is shown in *Fig. 1A* in the context of the present study. Briefly, entire and vasectomised 8 week old C57BL6 males were fed either control normal protein, normal carbohydrate diet (NP-NC; **18% casein**, 21% sucrose, 42% corn starch, 10% corn oil; *n* = 16 entire and 8 vasectomised males) or isocaloric low protein, high carbohydrate diet (LP-HC; **9% casein**, 24% sucrose, 49% corn starch, 10% corn oil; *n* = 16 entire and 8 vasectomised males) for a period of 8-12 weeks. Diets were manufactured commercially (Special Dietary Services Ltd; UK) and their composition described previously^43^.

### F1 offspring generation

Virgin 8-week-old female C57BL/6 mice (*n* = 8 litters per treatment) were super-ovulated by intraperitoneal injections of pregnant mare serum gonadotrophin (1 IU) and human chorionic gonadotrophin (1 IU) 46-48 hours later. Intact NP-NC and LP-HC fed males were culled by cervical dislocation after a minimum of 8 weeks on respective diets. Sperm were isolated from caudal epididymi of NP-NC and LP-HC sires as described^3, 20^ and allowed to capacitate *in vitro* (37 °C, 135 mM NaCl, 5 mM KCl, 1 mM MgSO_4_, 2 mM CaCl_2_, 30 mM HEPES; supplemented immediately before use with 10mM lactic acid, 1 mM sodium pyruvate, 20 mg/mL^-1^ BSA, 25 mM NaHCO^3^). Females were artificially inseminated 12 h post human chorionic gonadotrophin injection with ∼10^7^ sperm and subsequently housed overnight with a vasectomized C57BL/6 male fed either NP-NC or LP-HC diet. Females were weighed regularly (every 4–5 days) for the detection of weight gain associated with a developing pregnancy. Four groups of offspring were generated, termed NN (NP-NC sperm and NP-NC seminal plasma), LL (LP-HC sperm and LP-HC seminal plasma), NL (NP-NC sperm and LP-HC seminal plasma) and LN (LP-HC sperm and NP-NC seminal plasma). The number of females inseminated, pregnancy rates, gestation lengths and litter parameters have been reported^3^. In the current study, we focused on tissues collected from F1 and F2 NN (NL-NC) and LL (LP-HC) groups as these provide a model for normal- and high carbohydrate intake in humans, and in order to reduce complicating factors.

### F2 offspring generation

16-week-old F1 males (*n* = 6 males per treatment group; each from a different litter) were mated naturally to virgin, 8-week-old female C57BL/6 mice acquired separately for mating with F1 males. Females were allowed to develop to term and all dams and F2 offspring received standard chow and water *ad libitum*.

### Tissue collection

F1 offspring were culled by cervical dislocation at either 3 (juvenile) or 16 (adult) weeks of age, whereas all F2 offspring were culled by cervical dislocation at 3 weeks of age. Blood samples were taken via heart puncture, centrifuged at 8k × *g* (4°C, 10 min) and the serum aliquoted and stored at –80°C. Liver, brain, heart and adipose were dissected, weighed, snap frozen and stored at –80°C.

### Stock solutions

1. GCTU. Guanidine (6 M guanidinium chloride) and thiourea (1·5 M) were dissolved in deionised H_2_O together and stored at room temperature out of direct sunlight.
2. DMT. Dichloromethane (3 parts), methanol (1 part) and triethylammonium chloride (0.0005 parts, *i*.*e*. 500 mg/L) were mixed and stored at room temperature out of direct sunlight.
3. MS-mix. Propan-2-ol (2 parts) was mixed with methanol (1 part) and used to produce a solution of CH_3_COO.NH_4_ (7·5 mM).

### Tissue sample preparation and extraction of the lipid fraction

Whole tissue/organ samples were prepared and extracted as described recently^29, 44^. Solutions of homogenized organ preparations were injected into a well (96 well plate, Esslab Plate+™, 2·4 mL/well, glass-coated) followed by methanol spiked with internal standards (150 µL, internal standards shown in *Supplementary Table 1*), water (500 µL) and DMT (500 µL) using a 96 channel pipette. The mixture was agitated (96 channel pipette) before being centrifuged (3·2k × *g*, 2 min). A portion of the organic solution (20 µL) was transferred to a high throughput plate (384 well, glass-coated, Esslab Plate+™) before being dried (N_2 (g)_). When 4 × 96 well plates had been placed in the 384 well and the instrument was available, the dried films were re-dissolved (*tert*-butylmethyl ether, 20 µL/well, and MS-mix, 80 µL/well) and the plate was heat-sealed and queued immediately, with the first injection within 10 min.

Samples with a high concentration of triglycerides (TGs; *e*.*g*. adipose tissue) were also treated to concentrate the phospholipid fraction so it too could be profiled^28, 29, 44^.. A second portion of the organic phase from the extraction (100 µL) of was transferred to a shallow plate (96 well, glass-coated) before being dried (N_2 (g)_), washed (hexane, 3 × 100 µL/well) and re-dissolved (DMT, 30 µL). The samples were transferred immediately to the high throughput analytical plate as above and dried (N_2 (g)_).

### Direct Infusion Mass Spectrometry (DI-MS)

All samples were infused into an Exactive Orbitrap (Thermo, Hemel Hampstead, UK), using a TriVersa NanoMate (Advion, Ithaca US), for direct infusion mass spectrometry (DI-MS^27^). Samples (15 μL ea.) were sprayed at 1·2 kV in the positive ion mode. The Exactive started acquiring data 20 s after sample aspiration began. The Exactive acquired data with a scan rate of 1 Hz (resulting in a mass resolution of 100,000 full width at half-maximum [fwhm] at 400 *m/z*). The Automatic Gain Control was set to 3,000,000 and the maximum ion injection time to 50 ms. After 72 s of acquisition in positive mode the NanoMate and the Exactive switched over to negative ionization mode, decreasing the voltage to −1·5 kV and the maximum ion injection time to 50 ms. The spray was maintained for another 66 s, after which the NanoMate and Exactive switched over to negative mode with collision-induced dissociation (CID, 70 eV) for a further 66 s. After this time, the spray was stopped and the tip discarded, before the analysis of the next sample began. The sample plate was kept at 15 °C throughout the acquisition. Samples were run in row order. The instrument was operated in full scan mode from *m/z* 150–1200 Da.

### DI-MS Data processing

The lipid signals obtained were relative abundance (‘semi-quantitative’), with the signal intensity of each lipid expressed relative to the total lipid signal intensity, for each individual, per mille (‰). Raw high-resolution mass-spectrometry data were processed using XCMS (www.bioconductor.org) and Peakpicker v 2.0 (an in-house R script^27^). Lists of known species (by *m/z*) were used for both positive ion and negative ionisation mode (∼8k species). Signals that deviated by more than 9 ppm were ignored, as were those with a signal/noise ratio of <3 and those pertaining to fewer than 50% of samples. The correlation of signal intensity to concentration of plasma in QCs (0.25, 0.5, 1.0×) was used to identify which lipid signals were linearly proportional to abundance in the sample type and volume used (threshold for acceptance was a correlation of >0.75). Signals were then signal corrected (divided by the sum of signals for that sample not including internal standards), in order to be able to compare samples unconfounded by total lipid mass. All statistical calculations were done on these finalised values. Final signals files are in *Supplementary data*, in the form of 32× csv files. ‘(PW)’ refers to adipose that was washed with petrol; the data from petrol-washed samples were used for negative ionisation mode (in which phospholipids are measured) where untreated samples were used for positive ionisation mode (in which triglycerides and their fragmentation products were measured).

### Lipid extraction and sample preparations for ^31^P NMR

The extraction of larger sample volumes for NMR was based on a method described previously^29, 31^. Tissue homogenates were combined to give 5-10 mg of phospholipid per NMR sample. The samples of serum and prepared brain tissues from all groups were pooled and GCTU (250 µL) added to serum mixtures. Pooled solutions (5-8 mL) were diluted (DMT, 15 mL, Falcon tube) and made uniphasic (methanol, 15 mL). The mixture was agitated and diluted and made biphasic (dichloromethane, 10 mL) before centrifugation (3·2k × *g*, 2 min). The aqueous portion and any mesophasic solid was removed and discarded, and the organic solution dried under a flow of nitrogen. Samples were stored at −80 °C. Samples were dissolved in a modified^29, 30^ form of the ‘CUBO’ solvent system^45-48^ (the amount of dueteriated dimethylformamide *d*_7_ -DMF was minimised). Stock solutions of the solvent consisted of dimethylformamide (3·5 mL), *d*_7_ -DMF (1·5 mL), triethylamine (1·5 mL) and guanidinium chloride (500 mg). Wilmad^®^ 507PP tubes were used. Sample concentration was 5-10 mg lipids per sample (600 µL).

### NMR spectrometer and probe

Lipid samples were run on a Bruker Avance Neo 800 MHz spectrometer, equipped with a QCI cryoprobe probe. 1D Phosphorous experiments were acquired using inverse gated proton decoupling. Spectra were averaged over 1312 transients with 3882 complex points with a spectral width of 14·98 ppm. An overall recovery delay of 8·4 s was used. Data were processed using an exponential line broadening window function of 1·5 Hz prior to zero filling to 32768 points and Fourier transform. Data were processed and deconvoluted using TopSpin 4.07. Subsequent integrations above a noise threshold of 0·01% of the total ^31^P were used to establish the relative molar quantity of a given phosphorus environment. A survey of ^31^P traces is in *Supplementary Data*.

### Interpretation of profiling data and preparation of final lipidomics sheets

Dual spectroscopy^29^ was used to interpret lipidomics data. Specifically: ^31^P NMR data of hearts and livers from all generations and both phenotypes were collected and assigned (according to refs^29, 31, 45-48^) and compared and found to be much more similar to one another than other sample types (tissues/compartments). Only a small number of representative, pooled samples from the CNS and serum were therefore run. One liver sample (F2N, NP-NC) was run twice, 48h apart, to assess degradation within the sample. It was found that a small change in the abundance of *lyso-*PC was just measurable in this time, suggesting that sample preparation and running (<72h) was sound. One large scale petrol-wash^28^ was done on an adipose sample (F1A, LP-HC). One large scale petrol-wash^28^ was done on an adipose sample (F1A, LP-HC). These spectra were used to check for sample degradation in handling (*e*.*g*. appearance of PA) and inform assignments of signals measured using DI-MS. For example, serum has around 100× PC than PE, with very little or no PS, indicating that the balance of probabilities for assignments falls on the PC rather than the isobaric PE (positive ionisation mode) or PS (negative ionisation mode) isoform. These spectra were also used to interpret the difference in ionisation efficiency between species. These data show that the ionisation efficiency of *lyso-*PC and *lyso-*PE are both very high in negative ionisation mode, where that of sphingomyelin is under-represented in both ionisation modes.

### Statistical methods

Univariate and bivariate statistical calculations were made using Microsoft Excel 2016, as were calculations of Eq. 1 and 2. Graphs were prepared in OriginLab 2018 or Excel 2016 from mean (including Eq. 1) and standard deviation or error-normalised fold change (Eq. 2) as appropriate. Eq. 1 and 2 were generated *de novo* in the present study. Jaccard-Tanimoto Coefficients (JTCs) were used as a non-parametric measure of the distinctions between lipid variables associated with phenotype(s)^32, 33^. The associated *p*-values were calculated following Rahman *et al*.^49^. The *p*-value associated with each *J* represents the probability that the difference between the lists of variables for the two phenotypes occurred by random chance and should not be confused with *p*-values from Student’s *t-*test. The *p*- values that are associated with the Student’s *t*-tests (Abundance Analyses) were interpreted using a corrected *p*-value of 0·0021 based on 586 dependent variables^37^. Only lipid variables with a *p*-value below this and that were relevant to the hypothesis were used.

### Lipid Traffic Analysis

The tissues used were mapped to the known biological/metabolic network (*Fig. 1*). Lipid variables in each compartment (lipid station) were categorised according to whether they are unique to it (***U*** type lipids), shared with one adjacent to it (***B*** type lipids, uni- and bidirectional) or found in all compartments (***A*** type lipids), as shown in *Fig. 2*. Dimensions for the Abundance Analysis were calculated using Equations 1 and 2 (*vide supra*). Variables were regarded as present if they had a signal strength >0 in ≥50% of samples of either phenotype group.

Novel code for the Binary Traffic analysis (for the Switch analysis) and Multinary Traffic Analysis (Abundance analysis) was written in R(v3.6.x)^50^ and processed in RStudio(v1.2.5x). The full code can be found in the *Supplementary Information*. Briefly, MS signals data in Excel-readable *.csv format was uploaded with removal of the metadata (organ, extraction, plate location, enumeration of mass/charge ratios [*m/z*]), giving n (rows of observations) vs p (columns of lipids) of signal data. Layered functions were used to identify which variables were present in all (***A***), adjacent (***B***) or single (***U***) compartments. For each observation, the detection of the signal data commenced initially with FALSE representing no lipid signal (NA) and TRUE representing abundance of a lipid (above a signal threshold). For a particular compartment (tissue/pool/station), all observations were sampled into a single binary vector of presence and absences. The detection was performed using non-redundant lipid names. The function Reduce(intersect, list (…)) represented the common lipids for a given axis. Matched lipids were obtained across each pool to identify the common intersection, SetA(All). Positive and negative ionisation mode mass spectrometry data were run in series. The lists of lipids for the NP-NC and LP-HC groups were processed for the common intersection giving SetA (***A***, ubiquitous lipid variables), SetB (***B***, lipid variables found in two adjacent compartments) and SetU (***U***, unique, for lipid variables found in one compartment but not its neighbours).

## Acknowledgements

The authors wish to thank Drs S. G. Snowden and A Sleigh for helpful discussions. The authors also gratefully acknowledge funding from the BBSRC (BB/M027252/1 and BB/M027252/2 for SF). AJW was supported by an Aston Research Centre for a Healthy Ageing fellowship. SF would like to dedicate this paper to Cryptographer W. Gordon Welchman (1906-1985) whose work on Traffic Analysis for wartime communications inspired the method for characterising lipid traffic described here.

## Author Contributions

AJW developed the mouse model, did all animal work and produced all tissue samples. SF conceived and supervised the research, did all lipidomics, analysed data and wrote the manuscript. JS and DC designed the code and developed the lipid categories. NH, DC and JS wrote the code. HELW collected and analysed NMR data with SF. SF and AK designed the experiments. AK, JS and AJW wrote the original grant proposals. SF and AK analysed and interpreted data and revised the manuscript with comments from all authors. All authors commented on the manuscript and approved the final version.

## Electronic Supplementary Material

1. MS signals files (32× csv files)
2. R code for Lipid Traffic Analysis
3. Full Switch analysis results
4. ^31^P NMR spectra (1× pptx file)
5. Supplementary table

## Extended data

**Extended Data Table 1.** Triglyceride variables on the Liver-Serum (Liv-Ser) axis in the control (NP-NC) group that also appear on the Serum-Cerebellum (Ser-Ceb), Serum-Right Brain (Ser-RiB), Serum-Heart (Ser-Hea) or Serum-Adipose (Ser-Adi) axes of the control group. ^ø^Ammoniated adduct, ^§^Protonated, water-loss ion; *Sodiated adduct. DG, diglyceride (water-loss adduct from fragmentation in source); TG, triglyceride.

| F1A Liv-Ser<br>NP-NC | F1A Ser-Ceb<br>NP-NC | F1A Ser-RiB<br>NP-NC | F1A Ser-Hea<br>NP-NC | F1A Ser-Adi<br>NP-NC |
| --- | --- | --- | --- | --- |
| TG(33:00)<sup>a</sup> | TG(33:00)<sup>a</sup> | TG(33:00)<sup>a</sup> | TG(33:00)<sup>a</sup> |  |
|  | TG(35:01)<sup>a</sup> | TG(35:01)<sup>a</sup> |  |  |
| TG(48:01)<sup>a</sup> |  |  | TG(48:01)<sup>a</sup> |  |
| TG(50:02)<sup>a</sup> |  |  | TG(50:02)<sup>a</sup> | TG(50:02)<sup>a</sup> |
| TG(50:03)<sup>a</sup> |  |  | TG(50:03)<sup>a</sup> | TG(50:03)<sup>a</sup> |
| TG(52:01)<sup>a</sup> |  |  | TG(52:01)<sup>a</sup> | TG(52:01)<sup>a</sup> |
| TG(52:02)<sup>a</sup> | TG(52:02)<sup>a</sup> |  | TG(52:02)<sup>a</sup> | TG(52:02)<sup>a</sup> |
| TG(52:03)<sup>a</sup> | TG(52:03)<sup>a</sup> |  | TG(52:03)<sup>a</sup> | TG(52:03)<sup>a</sup> |
| TG(52:05)<sup>a</sup> | TG(52:05)<sup>a</sup> |  | TG(52:05)<sup>a</sup> | TG(52:05)<sup>a</sup> |
| TG(54:02)<sup>a</sup> |  |  | TG(54:02)<sup>a</sup> |  |
| TG(54:03)<sup>a</sup> |  |  | TG(54:03)<sup>a</sup> | TG(54:03)<sup>a</sup> |
| TG(54:04)<sup>a</sup> |  |  | TG(54:04)<sup>a</sup> | TG(54:04)<sup>a</sup> |
| TG(54:05)<sup>a</sup> |  |  | TG(54:05)<sup>a</sup> | TG(54:05)<sup>a</sup> |
| TG(54:06)<sup>a</sup> |  |  | TG(54:06)<sup>a</sup> | TG(54:06)<sup>a</sup> |
| TG(54:07)<sup>a</sup> |  |  | TG(54:07)<sup>a</sup> |  |
| TG(56:05)<sup>a</sup> |  |  | TG(56:05)<sup>a</sup> | TG(56:05)<sup>a</sup> |
| TG(56:06)<sup>a</sup> |  |  | TG(56:06)<sup>a</sup> |  |
| TG(56:07)<sup>a</sup> |  |  | TG(56:07)<sup>a</sup> |  |
| TG(56:08)<sup>a</sup> |  |  | TG(56:08)<sup>a</sup> |  |
|  | DG(20:00)<sup>§</sup> |  |  |  |
| DG(26:00)<sup>§</sup> |  |  | DG(26:00)<sup>§</sup> |  |
|  |  | DG(31:02)<sup>§</sup> |  |  |
| DG(32:03)<sup>§</sup> |  |  | DG(32:03)<sup>§</sup> |  |
| DG(33:01)<sup>§</sup> |  |  | DG(33:01)<sup>§</sup> | DG(33:01)<sup>§</sup> |
| DG(34:02)<sup>a</sup> | DG(34:02)<sup>a</sup> | DG(34:02)<sup>a</sup> | DG(34:02)<sup>a</sup> |  |
| DG(34:03)<sup>§</sup> |  |  | DG(34:03)<sup>§</sup> | DG(34:03)<sup>§</sup> |
|  |  | DG(36:01)<sup>§</sup> |  |  |
| DG(36:05)<sup>§</sup> |  |  | DG(36:05)<sup>§</sup> |  |
| DG(37:04)<sup>§</sup> |  | DG(37:04)<sup>§</sup> |  |  |
| DG(40:04)<sup>§</sup> | DG(40:04)<sup>§</sup> | DG(40:04)<sup>§</sup> | DG(40:04)<sup>§</sup> |  |
| DG(40:07)<sup>§</sup> | DG(40:07)<sup>§</sup> | DG(40:07)<sup>§</sup> | DG(40:07)<sup>§</sup> |  |
| DG(42:04)<sup>§</sup> | DG(42:04)<sup>§</sup> |  | DG(42:04)<sup>§</sup> |  |
|  | DG(44:07)<sup>§</sup> | DG(44:07)<sup>§</sup> | DG(44:07)<sup>§</sup> | DG(44:07)<sup>§</sup> |
| MG(20:03)<sup>a</sup> | MG(20:03)<sup>a</sup> | MG(20:03)<sup>a</sup> | MG(20:03)<sup>a</sup> |  |

**Extended Data Table 2.**
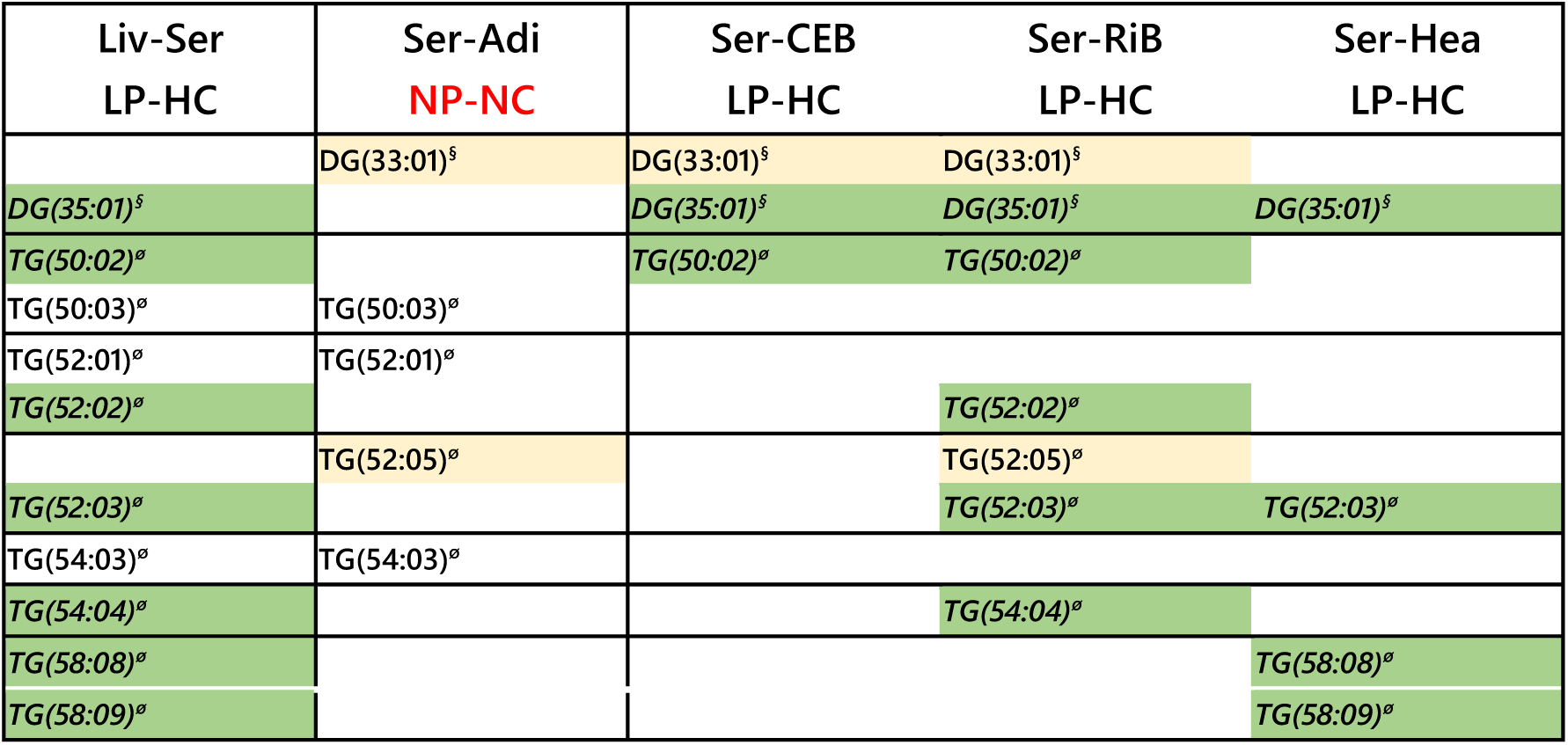
Triglyceride variables on the Liver-Serum (Liv-Ser) axis in the control (NP-NC) group of F1As that also appear on the Serum-Cerebellum (Ser-Ceb), Serum-Right Brain (Ser-RiB), Serum-Heart (Ser-Hea) or Serum-Adipose (Ser-Adi) axes of the control group. ^§^Protonated, water-loss ion; *Sodiated adduct; ^ø^Ammoniated adduct. DG, diglyceride (water-loss adduct from fragmentation in source); TG, triglyceride.

| Liv-Ser<br>LP-HC | Ser-Adi<br>**NP-NC** | Ser-CEB<br>LP-HC | Ser-RiB<br>LP-HC | Ser-Hea<br>LP-HC |
| --- | --- | --- | --- | --- |
|  | DG(33:01)<sup>§</sup> | DG(33:01)<sup>§</sup> | DG(33:01)<sup>§</sup> |  |
| DG(35:01)<sup>§</sup> |  | DG(35:01)<sup>§</sup> | DG(35:01)<sup>§</sup> | DG(35:01)<sup>§</sup> |
| TG(50:02)<sup>°</sup> |  | TG(50:02)<sup>°</sup> | TG(50:02)<sup>°</sup> |  |
| TG(50:03)<sup>°</sup> | TG(50:03)<sup>°</sup> |  |  |  |
| TG(52:01)<sup>°</sup> | TG(52:01)<sup>°</sup> |  |  |  |
| TG(52:02)<sup>°</sup> |  |  | TG(52:02)<sup>°</sup> |  |
|  | TG(52:05)<sup>°</sup> |  | TG(52:05)<sup>°</sup> |  |
| TG(52:03)<sup>°</sup> |  |  | TG(52:03)<sup>°</sup> | TG(52:03)<sup>°</sup> |
| TG(54:03)<sup>°</sup> | TG(54:03)<sup>°</sup> |  |  |  |
| TG(54:04)<sup>°</sup> |  |  | TG(54:04)<sup>°</sup> |  |
| TG(58:08)<sup>°</sup> |  |  |  | TG(58:08)<sup>°</sup> |
| TG(58:09)<sup>°</sup> |  |  |  | TG(58:09)<sup>°</sup> |

**Extended Data Table 3.**
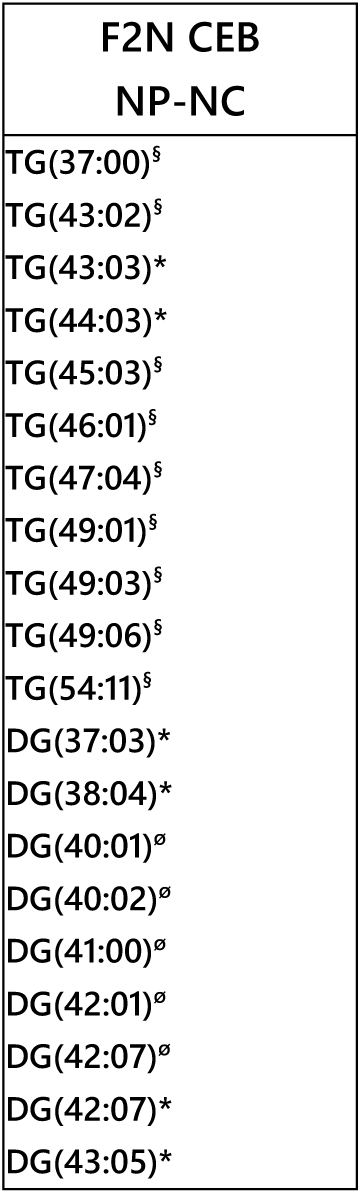
Triglyceride variables unique to the cerebella of normal protein-normal carbohydrate (NP-NC) F2N individuals. ^§^Sodiated adduct; *Ammoniated adduct; ^ø^Protonated, water-loss ion. CEB, cerebellum; DG, diglyceride (water-loss adduct from fragmentation in source); TG, triglyceride.

**Extended Data Table 4.** Phosphatidylcholine variables on the Liver-Serum (Liv-Ser) axis in the control (NP-NC) group of F2Ns that are also found on the Serum-Cerebellum (Ser-Ceb) and/or Serum-Heart (Ser-Hea) axis of the same (NP-NC) group. ^§^Chloride adduct; *acetate adduct. PC, phosphatidylcholine.

| F2N Liv-Ser<br>NP-NC | F2N Ser-Ceb<br>NP-NC | F2N Ser-Hea<br>NP-NC |
| --- | --- | --- |
|  |  | PC(30:3)*<br>PC(36:3)*<br>PC(38:5)* |
|  | PC(39:2)* |  |
| PC(39:3)<sup>§</sup> | PC(39:3)<sup>§</sup> |  |
|  | PC(39:4)<sup>§</sup> |  |
|  |  | PC(39:7)*<br>PC(39:8)<sup>§</sup> |
| PC(40:4)<sup>§</sup> | PC(40:4)<sup>§</sup> |  |
| PC(40:6)<sup>§</sup> | PC(40:6)<sup>§</sup> | PC(40:6)<sup>§</sup> |
|  |  | PC(40:7)* |
| PC(41:4)<sup>§</sup> | PC(41:4)<sup>§</sup> |  |
| PC(42:4)<sup>§</sup> |  |  |
| PC(44:2)* |  |  |
| PC(46:2)* |  |  |

**Extended Data Fig. 1.**
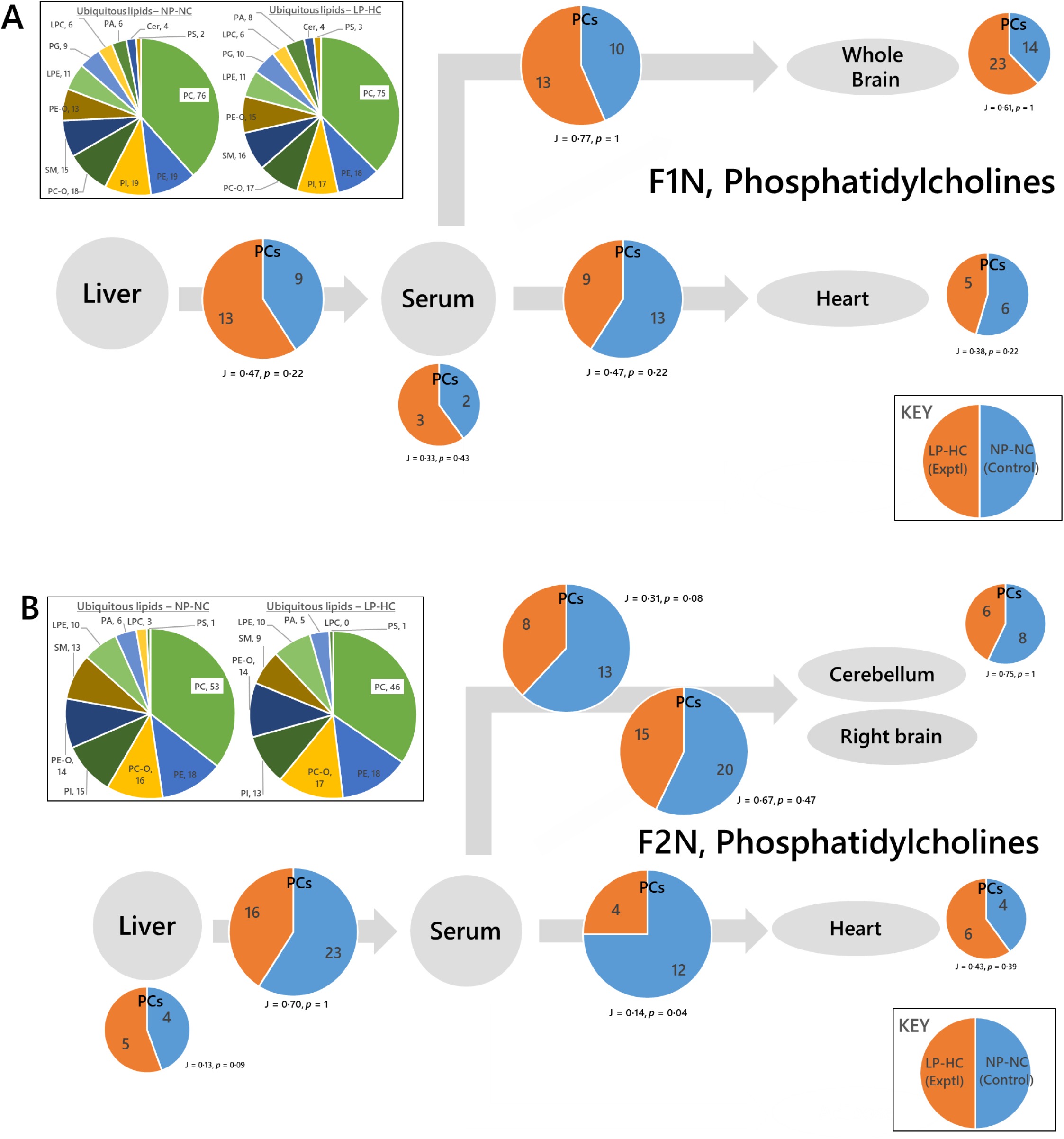
Switch analyses of phosphatidylcholines in control (NP-NC) mice and a phenotype whose (grand)sires were fed a low-protein, high carbohydrate (LP-HC) diet, measured by mass spectrometry in negative ionisation mode. Panel A, F1 Neonates (F1N); B, F2 Neonates (F2N). The pie charts in the insert show the number of ubiquitous lipid variables for that network, for each phenotype. Larger pie charts with J values represent PC variables found in two adjacent compartments. Smaller pie charts with J values represent PC variables found in isolated in compartments. J represents the Jaccard-Tanimoto coefficient for the comparison, with accompanying p-value, as a measure of the similarity between the variables identified in the two phenotypes for each comparison. The p-value shown represents the probability that the difference between the lists of variables for the two phenotypes occurred by random chance. LPC, lyso-phosphatidylcholine; LPE lyso-phosphatidylethanolamine; PA, phosphatidic acid; PC, phosphatidylcholine; PC-O, phosphatidylcholine plasmalogen; PE, phosphatidylethanolamine; PE-O, phosphatidylethanolamine plasmalogen; PG, phosphatidylglycerol; PI, phosphatidylinositol; PS, phosphatidylserine; SM, sphingomyelin; TG, triglyceride.

**Extended Data Fig. 2.**
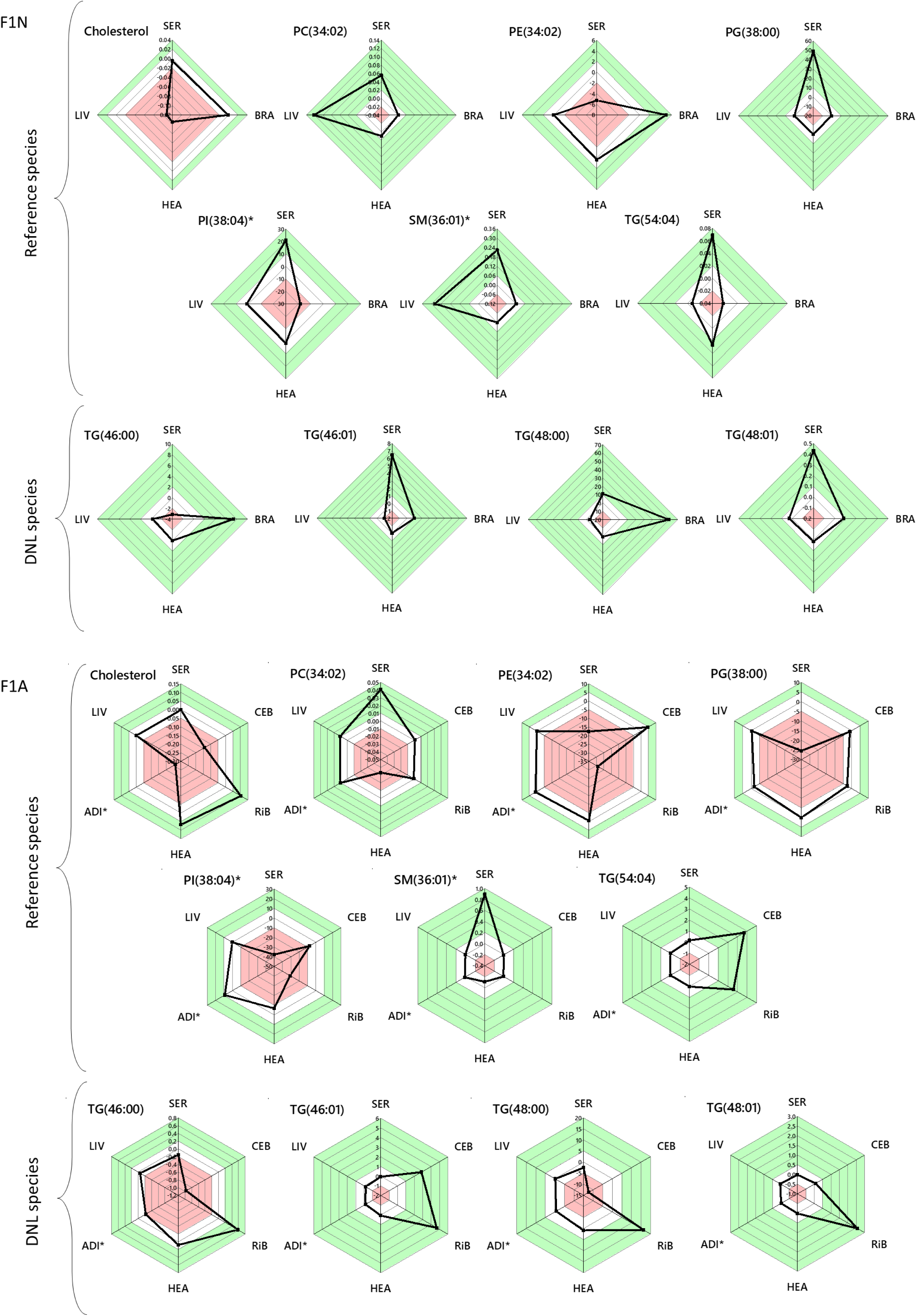
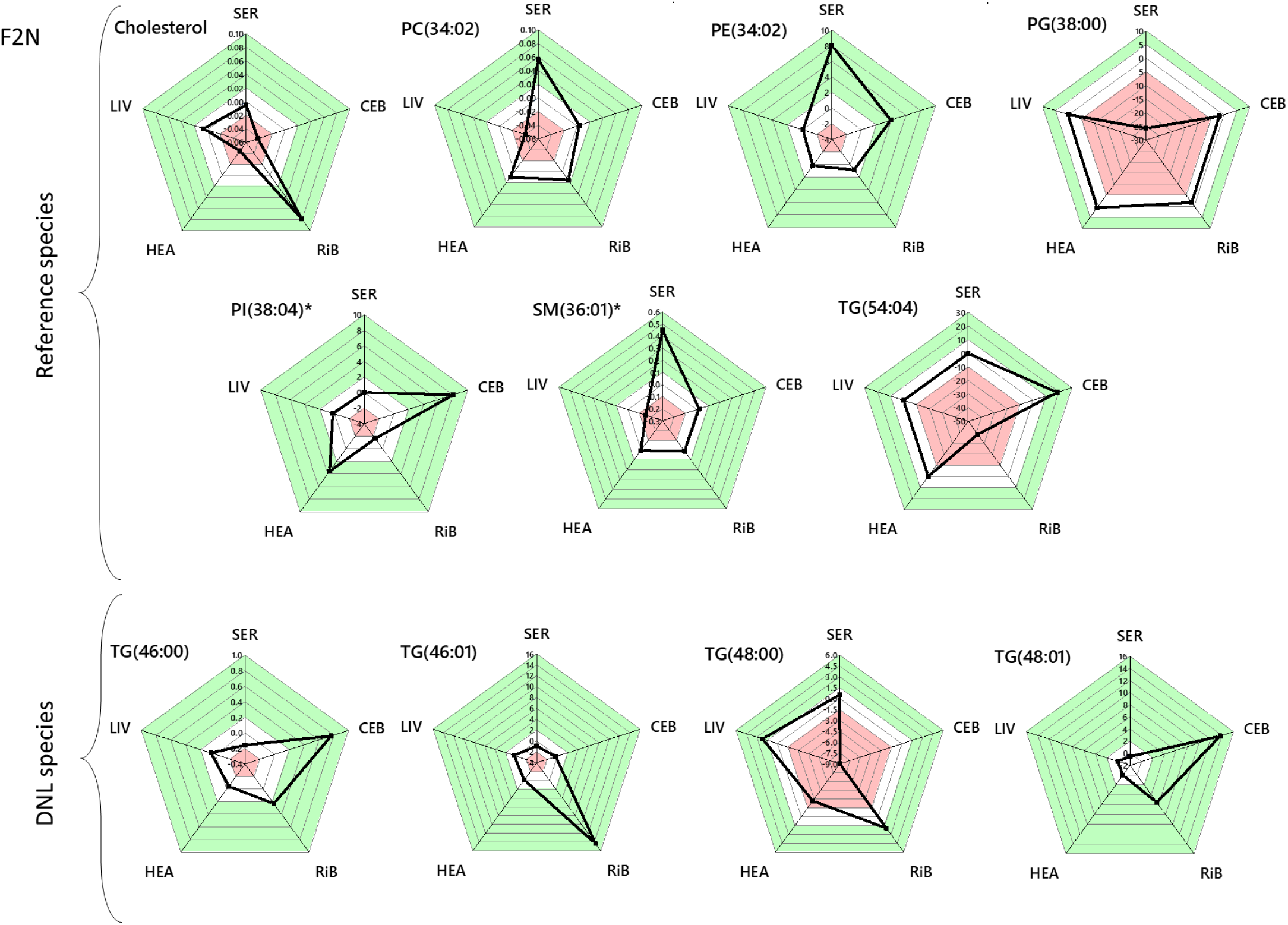
Radar plots of the fold change in abundance of lipid variables (error-normalised) unrelated (rows 1 and 2) and related (row 3) to de novo lipogenesis in mice. Panel A, F1 Neonate compartments; B, F1 adult compartments; C, F2 neonate compartments. The value given is the log of the mean of experimental abundance values divided by the mean of control values, divided by the propagated error for that variable (Eq. 1). Values of 0 show no change between phenotypes, negative values show lower in the experimental group (LP-HC offspring), positive values show increased abundance in the LP-HC offspring. The white areas represent the 0 point and one division above and below this. The red areas represent values more negative, and the green areas values more positive than this. ADI*, adipose (petrol wash); BRA, brain; CEB, cerebellum; HEA, heart; LIV, liver; RiB, right brain; SER, serum. PC, phosphatidylcholine; PE, phosphatidylethanolamine; PG, phosphatidylglycerol; PI, phosphatidylinositol; SM, sphingomyelin; TG, triglyceride.

## Supplementary Tables

**Supplementary Table 1.**
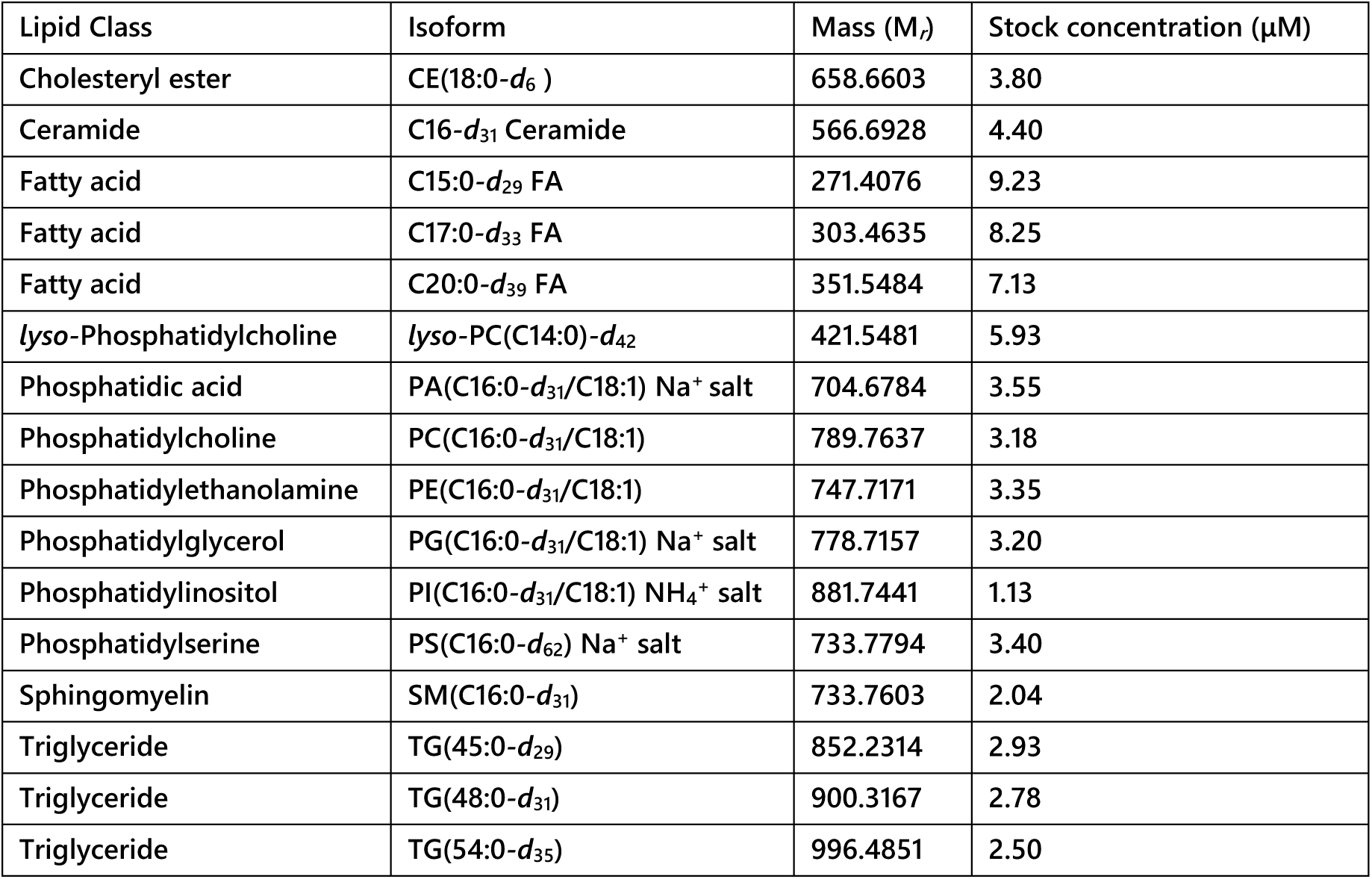
List of internal standards used for lipid profiling in the present study.

| Lipid Class | Isoform | Mass (M<sub>r</sub>) | Stock concentration (μM) |
| --- | --- | --- | --- |
| Cholesteryl ester | CE(18:0-*d*<sub>6</sub>) | 658.6603 | 3.80 |
| Ceramide | C16-*d*<sub>31</sub> Ceramide | 566.6928 | 4.40 |
| Fatty acid | C15:0-*d*<sub>29</sub> FA | 271.4076 | 9.23 |
| Fatty acid | C17:0-*d*<sub>33</sub> FA | 303.4635 | 8.25 |
| Fatty acid | C20:0-*d*<sub>39</sub> FA | 351.5484 | 7.13 |
| *lyso*-Phosphatidylcholine | *lyso*-PC(C14:0)-*d*<sub>42</sub> | 421.5481 | 5.93 |
| Phosphatidic acid | PA(C16:0-*d*<sub>31</sub>/C18:1) Na<sup>+</sup> salt | 704.6784 | 3.55 |
| Phosphatidylcholine | PC(C16:0-*d*<sub>31</sub>/C18:1) | 789.7637 | 3.18 |
| Phosphatidylethanolamine | PE(C16:0-*d*<sub>31</sub>/C18:1) | 747.7171 | 3.35 |
| Phosphatidylglycerol | PG(C16:0-*d*<sub>31</sub>/C18:1) Na<sup>+</sup> salt | 778.7157 | 3.20 |
| Phosphatidylinositol | PI(C16:0-*d*<sub>31</sub>/C18:1) NH<sub>4</sub><sup>+</sup> salt | 881.7441 | 1.13 |
| Phosphatidylserine | PS(C16:0-*d*<sub>62</sub>) Na<sup>+</sup> salt | 733.7794 | 3.40 |
| Sphingomyelin | SM(C16:0-*d*<sub>31</sub>) | 733.7603 | 2.04 |
| Triglyceride | TG(45:0-*d*<sub>29</sub>) | 852.2314 | 2.93 |
| Triglyceride | TG(48:0-*d*<sub>31</sub>) | 900.3167 | 2.78 |
| Triglyceride | TG(54:0-*d*<sub>35</sub>) | 996.4851 | 2.50 |

## References

1. Jazwiec, P.A. & Sloboda, D.M. Nutritional adversity, sex and reproduction: 30 years of DOHaD and what have we learned? 242, T51 (2019).

2. Tarry-Adkins, J.L. & Ozanne, S.E. Nutrition in early life and age-associated diseases. Ageing Research Reviews 39, 96–105 (2017).

3. Watkins, A.J. et al. Paternal diet programs offspring health through sperm- and seminal plasma-specific pathways in mice. Proceedings of the National Academy of Sciences 115, 10064 (2018).

4. Watkins, A.J. & Sinclair, K.D. Paternal low protein diet affects adult offspring cardiovascular and metabolic function in mice. American Journal of Physiology-Heart and Circulatory Physiology 306, H1444–H1452 (2014).

5. Furse, S. et al. Evidence that feeding post partum and exposures in utero shape lipid metabolism in infancy. Scientific Reports 9, 14321 (2019).

6. Furse, S. et al. Relationship between the lipid composition of maternal plasma and infant plasma through breast milk. Metabolomics 15, 129 (2019).

7. Duque-Guimarães, D.E. & Ozanne, S.E. Nutritional programming of insulin resistance: causes and consequences. Trends in Endocrinology & Metabolism 24, 525–535 (2013).

8. Yajnik, C.S., Godbole, K., Otiv, S.R. & Lubree, H.G. Fetal Programming of Type 2 Diabetes. Is sex important? 30, 2754–2755 (2007).

9. Fernandez-Twinn, D.S., Hjort, L., Novakovic, B., Ozanne, S.E. & Saffery, R. Intrauterine programming of obesity and type 2 diabetes. Diabetologia 62, 1789–1801 (2019).

10. Perng, W., Oken, E. & Dabelea, D. Developmental overnutrition and obesity and type 2 diabetes in offspring. Diabetologia 62, 1779–1788 (2019).

11. Ng, S.-F. et al. Chronic high-fat diet in fathers programs β-cell dysfunction in female rat offspring. Nature 467, 963–966 (2010).

12. Wei, Y. et al. Paternally induced transgenerational inheritance of susceptibility to diabetes in mammals. Proceedings of the National Academy of Sciences 111, 1873–1878 (2014).

13. Cropley, J.E. et al. Male-lineage transmission of an acquired metabolic phenotype induced by grand-paternal obesity. Molecular Metabolism 5, 699–708 (2016).

14. Schulze, M.B. & Hu, F.B. Dietary Approaches to Prevent the Metabolic Syndrome. Quality versus quantity of carbohydrates 27, 613–614 (2004).

15. Lee, Y.J., Song, S. & Song, Y. High-Carbohydrate Diets and Food Patterns and Their Associations with Metabolic Disease in the Korean Population. Yonsei Med J 59, 834–842 (2018).

16. Jeppesen, J. et al. Effects of low-fat, high-carbohydrate diets on risk factors for ischemic heart disease in postmenopausal women. Am J Clin Nutr 65, 1027–1033 (1997).

17. Hyde, P.N. et al. Dietary carbohydrate restriction improves metabolic syndrome independent of weight loss. JCI Insight 4 (2019).

18. Volek, J.S. & Feinman, R.D. Carbohydrate restriction improves the features of Metabolic Syndrome. Metabolic Syndrome may be defined by the response to carbohydrate restriction. Nutrition & metabolism 2, 31 (2005).

19. Dong, T., Guo, M., Zhang, P., Sun, G. & Chen, B. The effects of low-carbohydrate diets on cardiovascular risk factors: A meta-analysis. PLOS ONE 15, e0225348 (2020).

20. Morgan, H.L. et al. Paternal diet impairs F1 and F2 offspring vascular function through sperm and seminal plasma specific mechanisms in mice. The Journal of Physiology 598, 699 (2020).

21. Watkins, A.J. et al. Paternal low protein diet programs preimplantation embryo gene expression, fetal growth and skeletal development in mice. Biochimica et Biophysica Acta (BBA) - Molecular Basis of Disease 1863, 1371–1381 (2017).

22. Sanders, F. et al. Hepatic steatosis risk is partly driven by increased de novo lipogenesis following carbohydrate consumption. Genome Biology 19 (2018).

23. Copeland, W.B. et al. Computational tools for metabolic engineering. Metabolic Engineering 14, 270–280 (2012).

24. Wang, L., Dash, S., Ng, C.Y. & Maranas, C.D. A review of computational tools for design and reconstruction of metabolic pathways. Synth Syst Biotechnol 2, 243–252 (2017).

25. Guida, M.C. et al. Intergenerational inheritance of high fat diet-induced cardiac lipotoxicity in Drosophila. Nature Communications 10, 193 (2019).

26. Kilpeläinen, T.O. et al. Multi-ancestry study of blood lipid levels identifies four loci interacting with physical activity. Nature Communications 10, 376 (2019).

27. Harshfield, E.L. et al. An Unbiased Lipid Phenotyping Approach To Study the Genetic Determinants of Lipids and Their Association with Coronary Heart Disease Risk Factors. Journal of Proteome Research 18, 2397–2410 (2019).

28. Furse, S. & Koulman, A. The Lipid and Glyceride Profiles of Infant Formula Differ by Manufacturer, Region and Date Sold. Nutrients 11, 1122 (2019).

29. Furse, S. et al. A platform for detailed, high throughput lipidomics surveys of a range of mouse and human tissue types at a molecular level. Anal Bioanal Chem 412, 2851–2862 (2020).

30. Furse, S. et al. Evidence that Listeria innocua modulates its membrane’s stored curvature elastic stress, but not fluidity, through the cell cycle. Scientific Reports 7, 8012 (2017).

31. Furse, S. et al. The lipidome and proteome of oil bodies from Helianthus annuus (common sunflower). Journal of chemical biology 6, 63–76 (2013).

32. Jaccard, P. THE DISTRIBUTION OF THE FLORA IN THE ALPINE ZONE. New Phytologist 11, 37–50 (1912).

33. Tanimoto, T.T. (IBM, 1958).

34. Furse, S. & de Kroon, A.I.P.M. Phosphatidylcholine’s functions beyond that of a membrane brick. Molecular Membrane Biology 32, 117–119 (2015).

35. Banks, W.A. et al. Triglycerides cross the blood-brain barrier and induce central leptin and insulin receptor resistance. Int J Obes (Lond) 42, 391–397 (2018).

36. Bruce, K.D., Zsombok, A. & Eckel, R.H. Lipid Processing in the Brain: A Key Regulator of Systemic Metabolism. Front Endocrinol (Lausanne) 8 (2017).

37. Furse, S. et al. Altered triglyceride and phospholipid metabolism predates the diagnosis of gestational diabetes in obese pregnancy. Molecular Omics 15, 420–430 (2019).

38. Meek, C.L. et al. Metabolic insights into materno-fetal pregnancy complications of type 1 diabetes. A pre-specified analysis of the CONCEPTT trial. In prep (2020).

39. Mamtani, M. et al. Lipidomic risk score independently and cost-effectively predicts risk of future type 2 diabetes: results from diverse cohorts. Lipids Health Dis 15, 67 (2016).

40. Meikle, P.J. et al. Plasma Lipid Profiling Shows Similar Associations with Prediabetes and Type 2 Diabetes. PLOS ONE 8, e74341 (2013).

41. Aziz, R. & Mahboob, T. Pre-eclampsia and lipid profile. Pak J Med Sci 23, 751–754 (2007).

42. Gratacós, E. Lipid-mediated endothelial dysfunction: a common factor to preeclampsia and chronic vascular disease. European Journal of Obstetrics and Gynecology and Reproductive Biology 92, 63–66 (2000).

43. Watkins, A.J. et al. Paternal diet programs offspring health through sperm- and seminal plasma-specific pathways in mice. Proc Natl Acad Sci U S A 115, 10064–10069 (2018).

44. Furse, S., Torres, A.G. & Koulman, A. Fermentation of milk into yoghurt and cheese leads to contrasting lipid and glyceride profiles. Nutrients 11, 2178 (2019).

45. Bosco, M., Culeddu, N., Toffanin, R. & Pollesello, P. Organic solvent systems for P-31 nuclear magnetic resonance analysis of lecithin phospholipids: Applications to two-dimensional gradient-enhanced H-1-detected heteronuclear multiple quantum coherence experiments. Anal. Biochem. 245, 38–47 (1997).

46. Cremonini, M.A., Laghi, L. & Placucci, G. Investigation of commercial lecithin by P-31 NMR in a ternary CUBO solvent. Journal of the Science of Food and Agriculture 84, 786–790 (2004).

47. Culeddu, N., Bosco, M., Toffanin, R. & Pollesello, P. P-31 NMR analysis of phospholipids in crude extracts from different sources: improved efficiency of the solvent system. Magnetic Resonance in Chemistry 36, 907–912 (1998).

48. Murgia, S., Mele, S. & Monduzzi, M. Quantitative characterization of phospholipids in milk fat via P-31 NMR using a monophasic solvent mixture. Lipids 38, 585–591 (2003).

49. Rahman, S.A., Cuesta, S.M., Furnham, N., Holliday, G.L. & Thornton, J.M. EC-BLAST: a tool to automatically search and compare enzyme reactions. Nature Methods 11, 171–174 (2014).

50. Team, R.C. (R Foundation for Statistical Computing, Vienna, Austria, 2020).

