## Supplementary material for "Revealing grand-paternal programming of lipid metabolism using a novel computational tool": Switch analysis results for SI final.pdf

Supplementary data

Furse *et al.*

### Switch Analyses (all variables)

**Caption: Network diagrams showing which and how many lipid variables were detected across all compartments (pie charts, A type/ubiquitous lipids), between two tissues (blue tables, B type/adjacent lipids) and in only one compartment (orange tables, U type/unique lipids) in the two phenotypes studied. NP-NC, normal protein-normal carbohydrate; LP-HC, low protein, high carbohydrate. Cer, ceramide; Chol, cholesterol; LPC, *lyso*-phosphatidylcholine; LPE, *lyso*-phosphatidylethanolamine; LPG, *lyso*-phosphatidylglycerol; PA, phosphatidic acid; PC, phosphatidylcholine; PC-O, phosphatidylcholine plasmalogen; PE, phosphatidylethanolamine; PE-O, phosphatidylethanolamine plasmalogen; PG, phosphatidylglycerol; PI, phosphatidylinositol; PS, phosphatidylserine; S & SE, Sterols and Steryl Esters; SM, sphingomyelin; TG, triglyceride (comprises diglyceride water-loss adducts from fragmentation in source).**

### Sections

- F1 neonates, positive ionisation mode
- F1 adults, positive ionisation mode
- F2 neonates, positive ionisation mode
- F1 neonates, negative ionisation mode
- F1 adults, negative ionisation mode
- F2 neonates, negative ionisation mode

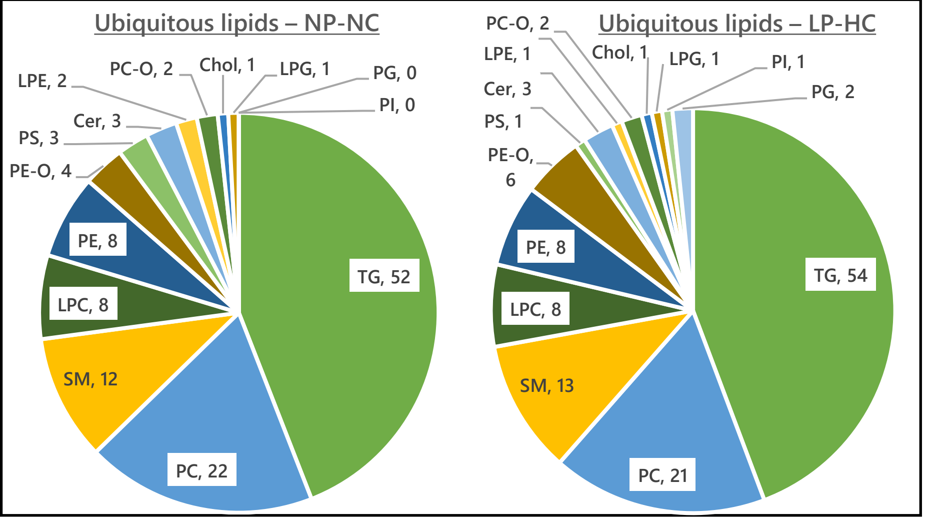


| | J | p |
|---|---|---|
| TG | 0·96 | 0·011 |
| PC | 0·95 | 1 |
| SM | 0·92 | 1 |
| LPC | 1 | 1 |
| PE | 1 | 1 |
| PE-O | 0·67 | 1 |
| PS | 0·33 | 1 |
| LPE | 0·50 | 1 |
| LPG | 1 | 1 |
| PC-O | 1 | 1 |
| PG | 0 | 1 |
| PI | 0 | 1 |
| Cer | 1 | 1 |
| Chol | 1 | 1 |

# F1N, +ve

Liver

$U_{LIV}$

| | NP-NC | LP-HC | J | p |
|---|---|---|---|---|
| TG | 2 | 0 | 0 | 1 |
| PI | 1 | 1 | 0 | 0·25 |

$B_{LIV-SER}$

| | NP-NC | LP-HC | J | p |
|---|---|---|---|---|
| TG | 12 | 13 | 0·39 | 0·08 |
| LPE | 3 | 2 | 0·33 | 1 |
| PG | 3 | 2 | 0 | 0·17 |
| PC | 2 | 2 | 0·33 | 0·44 |
| PI | 2 | 2 | 0·33 | 0·44 |
| PS | 2 | 2 | 0·33 | 0·44 |
| Cer | 1 | 2 | 0 | 0·15 |
| LPC | 0 | 1 | 0 | 1 |
| PC-O | 0 | 1 | 0 | 1 |
| PE | 0 | 3 | 0 | 1 |
| PE-O | 0 | 4 | 0 | 1 |
| SM | 0 | 3 | 0 | 1 |

Serum

$U_{SER}$

| | NP-NC | LP-HC | J | p |
|---|---|---|---|---|
| TG | 8 | 4 | 0·09 | 0·04 |
| PI | 6 | 0 | 0 | 1 |
| PS | 6 | 3 | 0 | 0·03 |
| S & SE | 3 | 4 | 0·4 | 0·36 |
| LPC | 2 | 2 | 1 | 1 |
| Cer | 1 | 2 | 0 | 0·15 |
| PA | 1 | 0 | 0 | 1 |
| PG | 1 | 0 | 0 | 1 |
| SM | 1 | 1 | 0 | 0·25 |

$B_{SER-BRA}$

| | NP-NC | LP-HC | J | p |
|---|---|---|---|---|
| TG | 48 | 65 | 0·59 | 0·19 |
| PS | 15 | 13 | 0·60 | 0·22 |
| PI | 14 | 20 | 0·45 | 0·13 |
| PG | 13 | 15 | 0·87 | 1 |
| Cer | 12 | 17 | 0·53 | 0·26 |
| S & SE | 7 | 9 | 0·75 | 0·39 |
| PC-O | 7 | 8 | 0·63 | 0·48 |
| LPG | 3 | 3 | 1 | 1 |
| LPI | 3 | 4 | 0·75 | 1 |
| LPC | 5 | 5 | 1 | 1 |
| PC | 5 | 7 | 0·50 | 0·40 |
| PE | 5 | 7 | 0·67 | 1 |
| LPE | 5 | 5 | 1 | 1 |
| SM | 4 | 8 | 0·43 | 1 |
| PE-O | 2 | 9 | 0·22 | 1 |
| PA | 1 | 2 | 0 | 1 |

Whole Brian

$U_{BRA}$

| | NP-NC | LP-HC | J | p |
|---|---|---|---|---|
| TG | 33 | 30 | 0·44 | 0·22 |
| PI | 15 | 13 | 0·55 | 0·23 |
| PS | 13 | 16 | 0·53 | 0·20 |
| PC-O | 11 | 14 | 0·56 | 0·31 |
| Cer | 11 | 8 | 0·11 | 0·36 |
| PC | 10 | 10 | 1 | 1 |
| PG | 9 | 10 | 0·46 | 0·19 |
| PI-O | 3 | 8 | 0·10 | 0·05 |
| SM | 6 | 5 | 0·37 | 0·22 |
| PE-O | 5 | 4 | 0·80 | 1 |
| PE | 4 | 4 | 0·33 | 0·24 |
| PA | 1 | 1 | 1 | 1 |
| LPS | 1 | 1 | 1 | 1 |
| S & SE | 0 | 3 | 0 | 1 |

$B_{SER-HEA}$

| | NP-NC | LP-HC | J | p |
|---|---|---|---|---|
| TG | 16 | 18 | 0·70 | 0·45 |
| PS | 4 | 1 | 0·25 | 1 |
| PC-O | 3 | 3 | 0·50 | 0·43 |
| SM | 3 | 3 | 0·50 | 0·43 |
| PI | 2 | 1 | 0 | 0·15 |
| LPO | 2 | 1 | 0·50 | 1 |
| PC | 2 | 0 | 0 | 1 |
| PE | 2 | 3 | 0·25 | 0·43 |
| LPE | 2 | 0 | 0 | 1 |
| PG | 1 | 3 | 0·33 | 1 |
| LPC | 1 | 0 | 0 | 1 |
| PE-O | 1 | 5 | 0·20 | 1 |
| CE | 0 | 2 | 0 | 1 |

Heart

$U_{HEA}$

| | NP-NC | LP-HC | J | p |
|---|---|---|---|---|
| TG | 6 | 4 | 0·25 | 0·15 |
| PC-O | 2 | 0 | 0 | 1 |

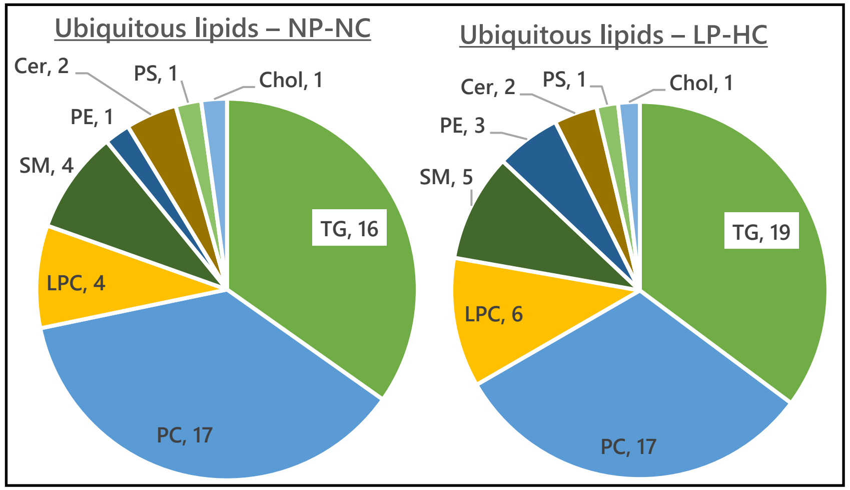


| | J | p |
|---|---|---|
| TG | 0·84 | 1 |
| PC | 0·89 | 0·41 |
| LPC | 0·67 | 1 |
| SM | 0·80 | 1 |
| PE | 1 | 3 |
| Cer | 2 | 2 |
| PS | 1 | 1 |
| Chol | 1 | 1 |
| PA | 0·33 | 1 |

# F1A, +ve

Liver → Serum → Cerebellum, Right brain, Heart, Adipose

$B_{SER-CEB}$

| | NP-NC | LP-HC | J | p |
|---|---|---|---|---|
| TG | 12 | 18 | 0·67 | 1 |
| PE-O | 5 | 4 | 0·80 | 1 |
| SM | 1 | 3 | 0·33 | 1 |
| PE | 5 | 6 | 0·83 | 1 |
| LPC | 4 | 5 | 0·80 | 1 |
| PC | 3 | 2 | 0·67 | 1 |
| LPG | 3 | 1 | 0·33 | 1 |
| Cer | 2 | 2 | 1 | 1 |
| LPE | 2 | 3 | 0·67 | 1 |
| SE | 1 | 1 | 1 | 1 |
| LPI | 1 | 0 | 0 | 1 |
| PS | 1 | 0 | 0 | 1 |

$B_{SER-RiB}$

| | NP-NC | LP-HC | J | p |
|---|---|---|---|---|
| TG | 10 | 20 | 0·50 | 0·16 |
| LPC | 4 | 5 | 0·80 | 1 |
| PE-O | 5 | 4 | 0·80 | 1 |
| PE | 3 | 4 | 0·75 | 1 |
| LPE | 2 | 3 | 0·67 | 1 |
| PC | 3 | 2 | 0·67 | 1 |
| LPG | 3 | 2 | 0·67 | 1 |
| Cer | 2 | 2 | 1 | 1 |
| SM | 1 | 2 | 0·50 | 1 |
| SE | 1 | 1 | 1 | 1 |
| PS | 1 | 0 | 0 | 1 |

Cerebellum

$U_{CEB}$

| | NP-NC | LP-HC | J | p |
|---|---|---|---|---|
| TG | 8 | 6 | 0·53 | 0·38 |
| PI | 2 | 4 | 0·50 | 1 |
| PC | 1 | 4 | 0·25 | 1 |
| PE | 2 | 2 | 0·33 | 0·44 |
| PS | 1 | 2 | 0·50 | 1 |
| SM | 2 | 1 | 0·50 | 1 |
| Cer | 1 | 1 | 0 | 0·33 |
| PC-O | 1 | 1 | 1 | 1 |

Right brain

$U_{RiB}$

| | NP-NC | LP-HC | J | p |
|---|---|---|---|---|
| TG | 0 | 2 | 0 | 1 |
| PI | 1 | 3 | 0·33 | 1 |
| PS | 1 | 2 | 0·50 | 1 |
| SM | 0 | 1 | 0 | 1 |
| PC | 1 | 0 | 0 | 1 |
| PE | 1 | 0 | 0 | 1 |
| SE | 1 | 0 | 0 | 1 |
| PG | 1 | 0 | 0 | 1 |

Liver

$B_{LIV-SER}$

| | NP-NC | LP-HC | J | p |
|---|---|---|---|---|
| TG | 29 | 36 | 0·80 | 0·36 |
| PE-O | 4 | 4 | 1 | 1 |
| SM | 4 | 6 | 0·67 | 1 |
| PE | 3 | 5 | 0·60 | 1 |
| LPC | 3 | 4 | 0·75 | 1 |
| PC | 2 | 2 | 1 | 1 |
| LPG | 2 | 1 | 0·50 | 1 |
| Cer | 1 | 1 | 1 | 1 |
| LPE | 0 | 2 | 0 | 1 |

Serum

$B_{SER-HEA}$

| | NP-NC | LP-HC | J | p |
|---|---|---|---|---|
| TG | 29 | 34 | 0·85 | 0·47 |
| PE-O | 5 | 4 | 0·80 | 1 |
| LPC | 4 | 5 | 0·80 | 1 |
| PE | 4 | 4 | 0·60 | 0·43 |
| SM | 3 | 5 | 0·60 | 1 |
| PC | 3 | 2 | 0·67 | 1 |
| LPE | 2 | 3 | 0·67 | 1 |
| LPG | 1 | 2 | 0·50 | 1 |
| PS | 1 | 0 | 0 | 1 |
| Cer | 0 | 1 | 0 | 1 |

Heart

$U_{HEA}$

| | NP-NC | LP-HC | J | p |
|---|---|---|---|---|
| TG | 12 | 14 | 0·37 | 0·06 |
| PI | 1 | 0 | 0 | 1 |
| PS | 1 | 1 | 1 | 1 |
| SM | 2 | 0 | 0 | 1 |
| PC | 0 | 1 | 0 | 1 |
| PG | 0 | 1 | 0 | 1 |
| PC-O | 1 | 0 | 0 | 1 |

$U_{LIV}$

| | NP-NC | LP-HC | J | p |
|---|---|---|---|---|
| TG | 4 | 2 | 0·20 | 0·22 |
| PI | 1 | 3 | 0·33 | 1 |
| SE | 0 | 1 | 0 | 1 |
| SM | 1 | 0 | 0 | 1 |
| PS | 1 | 1 | 1 | 1 |

$U_{SER}$

| | NP-NC | LP-HC | J | p |
|---|---|---|---|---|
| S & SE | 6 | 6 | 1 | 1 |
| LPC | 4 | 3 | 0·75 | 1 |
| LPE | 1 | 1 | 1 | 1 |
| PC | 1 | 0 | 0 | 1 |
| LPI | 0 | 1 | 0 | 1 |
| PG | 1 | 0 | 0 | 1 |
| SM | 1 | 1 | 1 | 1 |

$B_{SER-ADI}$

| | NP-NC | LP-HC | J | p |
|---|---|---|---|---|
| TG | 14 | 7 | 0·50 | 1 |
| LPE | 2 | 1 | 0·50 | 1 |
| SM | 2 | 1 | 0 | 0·29 |
| PE | 1 | 3 | 0·33 | 1 |
| LPC | 0 | 2 | 0 | 1 |
| PC | 1 | 1 | 1 | 0·13 |

Adipose

$U_{ADI}$

| | NP-NC | LP-HC | J | p |
|---|---|---|---|---|
| TG | 2 | 0 | 0 | 1 |
| PI | 2 | 1 | 0·50 | 1 |
| PS | 2 | 0 | 0 | 1 |
| SM | 1 | 1 | 1 | 1 |

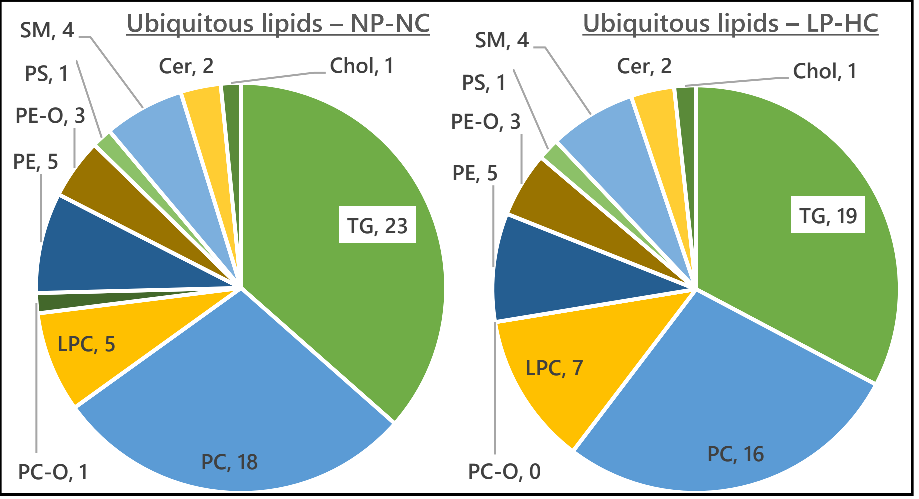


| | J | p |
|---|---|---|
| TG | 0·83 | 1 |
| PC | 0·95 | 0·03 |
| LPC | 0·71 | 1 |
| PC-O | 0 | 1 |
| PE | 0·67 | 0·43 |
| PE-O | 1 | 1 |
| PS | 1 | 1 |
| SM | 1 | 1 |
| Cer | 1 | 1 |
| Chol | 1 | 1 |

# F2N +ve

Liver

Serum

Cerebellum

Right brain

Heart

$B_{SER-CEB}$

| | NP-NC | LP-HC | J | p |
|---|---|---|---|---|
| TG | 19 | 12 | 0·48 | 0·21 |
| PC | 2 | 1 | 0 | 0·14 |
| LPC | 4 | 3 | 0·75 | 1 |
| PC-O | 1 | 0 | 0 | 1 |
| PE | 2 | 1 | 0 | 0·14 |
| LPE | 3 | 3 | 1 | 1 |
| PE-O | 4 | 4 | 1 | 1 |
| LPG | 3 | 2 | 0·67 | 1 |
| SM | 2 | 0 | 0 | 1 |
| PG | 1 | 1 | 1 | 1 |
| PI | 2 | 1 | 0·5 | 1 |
| PS | 1 | 1 | 1 | 1 |
| S & SE | 1 | 1 | 1 | 1 |
| Cer | 2 | 1 | 0·5 | 1 |
| LPI | 1 | 1 | 1 | 1 |

$B_{SER-RiB}$

| | NP-NC | LP-HC | J | p |
|---|---|---|---|---|
| TG | 10 | 11 | 0·62 | 0·40 |
| PC | 3 | 2 | 0·25 | 0·43 |
| LPC | 1 | 3 | 0·33 | 1 |
| PC-O | 1 | 1 | 0 | 1 |
| PE | 1 | 1 | 0 | 1 |
| LPE | 3 | 3 | 1 | 1 |
| PE-O | 1 | 1 | 1 | 1 |
| PG | 1 | 1 | 1 | 1 |
| LPG | 3 | 3 | 1 | 1 |
| Cer | 2 | 2 | 1 | 1 |
| LPI | 1 | 1 | 1 | 1 |
| PI | 2 | 1 | 0·5 | 1 |

$U_{CEB}$

| | NP-NC | LP-HC | J | p |
|---|---|---|---|---|
| TG | 32 | 12 | 0·38 | 1 |
| PG | 8 | 0 | 0 | 1 |
| PC | 7 | 2 | 0·29 | 1 |
| PC-O | 6 | 2 | 0·33 | 1 |
| SM | 6 | 2 | 0·33 | 1 |
| PI | 5 | 2 | 0·5 | 1 |
| PE | 4 | 3 | 0·4 | 0·41 |
| Cer | 4 | 2 | 0·5 | 1 |
| PS | 3 | 1 | 0 | 0·17 |
| PE-O | 1 | 1 | 1 | 1 |

$U_{RiB}$

| | NP-NC | LP-HC | J | p |
|---|---|---|---|---|
| TG | 0 | 1 | 0 | 1 |
| PC | 0 | 1 | 0 | 1 |
| PS | 1 | 4 | 0 | 0·07 |
| PI | 2 | 3 | 0·67 | 1 |
| SM | 0 | 1 | 0 | 1 |

$B_{LIV-SER}$

| | NP-NC | LP-HC | J | p |
|---|---|---|---|---|
| TG | 26 | 27 | 0·83 | 0·47 |
| PC | 2 | 1 | 0·5 | 1 |
| LPC | 2 | 3 | 0·67 | 1 |
| PC-O | 1 | 0 | 0 | 1 |
| PE | 1 | 1 | 0 | 0·25 |
| LPE | 2 | 1 | 0·5 | 1 |
| PE-O | 1 | 1 | 1 | 1 |
| LPG | 3 | 2 | 0·67 | 1 |
| SM | 3 | 2 | 0·67 | 1 |

$B_{SER-HEA}$

| | NP-NC | LP-HC | J | p |
|---|---|---|---|---|
| TG | 23 | 20 | 0·72 | 0·44 |
| PC | 2 | 0 | 0 | 1 |
| LPC | 1 | 3 | 0·33 | 1 |
| PC-O | 1 | 0 | 0 | 1 |
| PE | 1 | 1 | 0 | 0·25 |
| LPE | 1 | 1 | 1 | 1 |
| SM | 3 | 2 | 0·67 | 1 |
| Cer | 1 | 1 | 1 | 1 |

$U_{HEA}$

| | NP-NC | LP-HC | J | p |
|---|---|---|---|---|
| TG | 5 | 8 | 0·63 | 1 |
| PI | 0 | 3 | 0 | 1 |
| PC-O | 0 | 1 | 0 | 1 |
| PG | 1 | 0 | 0 | 1 |
| CE | 0 | 1 | 0 | 1 |

$U_{LIV}$

| | NP-NC | LP-HC | J | p |
|---|---|---|---|---|
| TG | 1 | 2 | 0 | 0·15 |
| PI | 1 | 0 | 0 | 1 |
| SM | 1 | 0 | 0 | 1 |
| CE | 1 | 0 | 0 | 1 |

$U_{SER}$

| | NP-NC | LP-HC | J | p |
|---|---|---|---|---|
| TG | 2 | 3 | 0 | 0·13 |
| S & SE | 3 | 3 | 1 | 1 |
| LPC | 5 | 3 | 0·6 | 1 |
| SM | 0 | 1 | 0 | 1 |

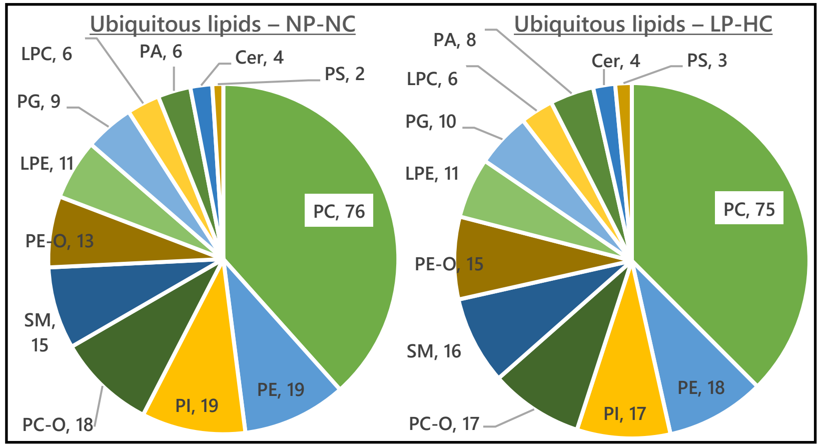


| | J | p |
|---|---|---|
| PC | 0·98 | 0·02 |
| PE | 0·95 | 1 |
| PI | 0·89 | 0·47 |
| PC-O | 0·94 | 1 |
| SM | 0·82 | 0·46 |
| PE-O | 0·87 | 1 |
| LPE | 1 | 1 |
| PG | 0·73 | 0·43 |
| LPC | 1 | 1 |
| PA | 0·75 | 1 |
| Cer | 1 | 1 |
| PS | 0·67 | 1 |

# F1N, -ve

Liver

$U_{LIV}$

| | NP-NC | LP-HC | J | p |
|---|---|---|---|---|
| Cer | 3 | 5 | 0·6 | 1 |
| PC | 0 | 1 | 0 | 1 |
| PG | 0 | 1 | 0 | 1 |
| PI | 0 | 1 | 0 | 1 |
| PS | 1 | 0 | 0 | 1 |

$B_{LIV\text{-}SER}$

| | NP-NC | LP-HC | J | p |
|---|---|---|---|---|
| PC | 9 | 13 | 0·47 | 0·22 |
| LPC | 12 | 10 | 0·83 | 1 |
| PC-O | 4 | 4 | 1 | 1 |
| SM | 4 | 3 | 0·75 | 1 |
| PI | 3 | 3 | 0·5 | 0·43 |
| PG | 3 | 3 | 0·5 | 0·43 |
| Cer | 3 | 2 | 0·67 | 1 |
| PE | 2 | 1 | 0·5 | 1 |
| LPE | 1 | 1 | 1 | 1 |
| PA | 1 | 2 | 0·5 | 1 |
| PE-O | 0 | 1 | 0 | 1 |
| PS | 0 | 1 | 0 | 1 |

Serum

$U_{SER}$

| | NP-NC | LP-HC | J | p |
|---|---|---|---|---|
| PC | 2 | 3 | 0·33 | 0·43 |
| LPC | 2 | 4 | 0·50 | 1 |
| PC-O | 4 | 4 | 0·33 | 0·24 |
| PE | 2 | 0 | 0 | 1 |
| PG | 0 | 1 | 0 | 1 |
| PI | 1 | 0 | 0 | 1 |
| PS | 7 | 5 | 0·71 | 1 |
| SM | 1 | 0 | 0 | 1 |

$B_{SER\text{-}BRA}$

| | NP-NC | LP-HC | J | p |
|---|---|---|---|---|
| PC | 10 | 13 | 0·77 | 1 |
| PC-O | 9 | 11 | 0·82 | 1 |
| PE | 7 | 10 | 0·55 | 0·42 |
| SM | 7 | 6 | 0·63 | 0·48 |
| PI | 5 | 7 | 0·33 | 0·17 |
| PG | 2 | 5 | 0·25 | 0·42 |
| Cer | 1 | 2 | 0·50 | 1 |
| PA | 1 | 2 | 0·50 | 1 |
| LPI | 1 | 1 | 1 | 1 |
| PE-O | 0 | 6 | 0 | 1 |
| PS | 0 | 1 | 0 | 1 |

Whole Brian

$U_{BRA}$

| | NP-NC | LP-HC | J | p |
|---|---|---|---|---|
| PC | 14 | 23 | 0·61 | 1 |
| PC-O | 8 | 13 | 0·62 | 1 |
| CL | 40 | 43 | 0·93 | 1 |
| PA | 1 | 1 | 1 | 1 |
| Cer | 2 | 2 | 1 | 1 |
| PE | 9 | 7 | 0·60 | 0·44 |
| PE-O | 14 | 15 | 0·71 | 0·45 |
| PG | 1 | 1 | 0 | 0·25 |
| PI | 4 | 3 | 0·75 | 1 |
| PS | 1 | 2 | 0 | 0·15 |
| SM | 0 | 2 | 0 | 1 |

$B_{SER\text{-}HEA}$

| | NP-NC | LP-HC | J | p |
|---|---|---|---|---|
| PC | 13 | 9 | 0·47 | 0·22 |
| LPC | 15 | 13 | 0·75 | 0·47 |
| PI | 4 | 1 | 0 | 0·07 |
| PC-O | 5 | 3 | 0·33 | 0·32 |
| PE | 4 | 2 | 0·22 | 0·20 |
| SM | 4 | 3 | 0·17 | 0·12 |
| LPE | 1 | 1 | 1 | 1 |
| PE-O | 0 | 3 | 0 | 1 |
| PG | 4 | 4 | 0·60 | 0·43 |
| PA | 0 | 2 | 0 | 1 |
| PS | 0 | 2 | 0 | 1 |

Heart

$U_{HEA}$

| | NP-NC | LP-HC | J | p |
|---|---|---|---|---|
| PI | 10 | 8 | 0·64 | 0·46 |
| PC | 6 | 5 | 0·38 | 0·22 |
| PC-O | 2 | 0 | 0 | 1 |
| PE | 2 | 0 | 0 | 1 |
| PG | 3 | 2 | 0·67 | 1 |
| PS | 1 | 0 | 0 | 1 |
| SM | 1 | 1 | 1 | 1 |
| Cer | 1 | 0 | 0 | 1 |
| PA | 0 | 1 | 0 | 1 |

# F1A, -ve

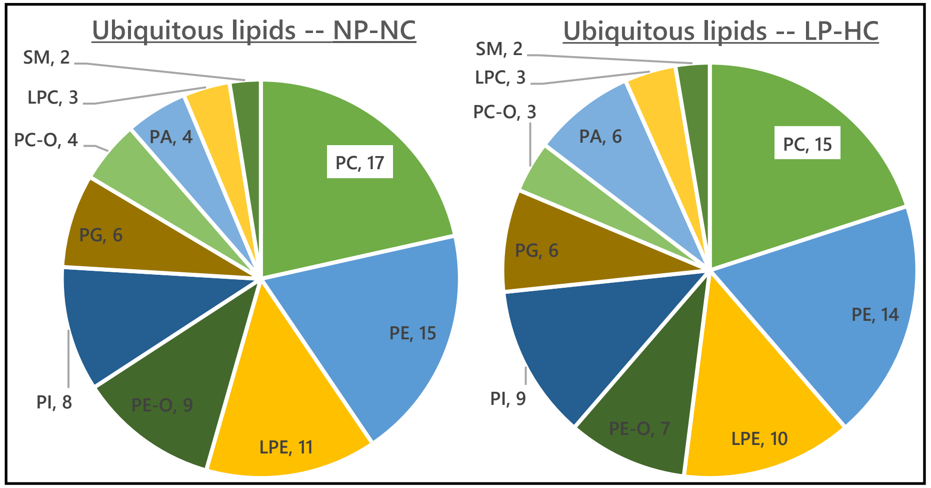


| | J | p |
|---|---|---|
| PC | 0·78 | 0·46 |
| PE | 0·93 | 1 |
| LPE | 0·91 | 1 |
| PE-O | 0·78 | 1 |
| PI | 0·7 | 1 |
| PG | 1 | 1 |
| PC-O | 0·75 | 1 |
| PA | 0·83 | 1 |
| LPC | 1 | 1 |
| SM | 1 | 1 |

Liver

Serum

Cerebellum

Right brain

Heart

Adipose

$B_{LIV-SER}$

| | NP-NC | LP-HC | J | p |
|---|---|---|---|---|
| PC | 55 | 55 | 0·84 | 0·49 |
| PI | 14 | 15 | 0·93 | 1 |
| LPC | 13 | 14 | 0·58 | 0·29 |
| PC-O | 12 | 15 | 0·69 | 0·47 |
| SM | 12 | 13 | 0·92 | 1 |
| PE | 8 | 8 | 0·78 | 0·43 |
| PE-O | 8 | 8 | 1 | 1 |
| PG | 4 | 5 | 0·80 | 1 |
| PS | 3 | 3 | 0·50 | 0·43 |
| LPE | 2 | 2 | 1 | 1 |
| Cer | 1 | 2 | 0·50 | 1 |
| PA | 1 | 3 | 0·33 | 1 |

$U_{LIV}$

| | NP-NC | LP-HC | J | p |
|---|---|---|---|---|
| PC | 2 | 4 | 0·20 | 0·22 |
| Cer | 3 | 2 | 0·67 | 1 |
| PA | 1 | 0 | 0 | 1 |
| PC-O | 0 | 1 | 0 | 1 |
| PI | 1 | 1 | 1 | 0·33 |
| SM | 1 | 1 | 1 | 1 |

$U_{SER}$

| | NP-NC | LP-HC | J | p |
|---|---|---|---|---|
| PC | 3 | 2 | 0·67 | 1 |
| LPC | 4 | 4 | 1 | 1 |
| PC-O | 2 | 4 | 0·2 | 0·22 |
| PI | 1 | 0 | 0 | 1 |
| PS | 7 | 7 | 1 | 1 |
| SM | 0 | 1 | 0 | 1 |

$B_{SER-CEB}$

| | NP-NC | LP-HC | J | p |
|---|---|---|---|---|
| PC | 48 | 50 | 0·89 | 0·47 |
| PC-O | 19 | 22 | 0·83 | 0·48 |
| PE-O | 14 | 17 | 0·82 | 1 |
| PI | 13 | 14 | 0·70 | 0·47 |
| SM | 13 | 15 | 0·87 | 1 |
| PE | 8 | 10 | 0·64 | 0·46 |
| LPC | 3 | 3 | 0·50 | 0·43 |
| PG | 3 | 3 | 0·75 | 1 |
| LPE | 1 | 1 | 0 | 1 |
| PA | 1 | 3 | 0·33 | 1 |
| PS | 1 | 2 | 0·50 | 1 |
| Cer | 0 | 1 | 0 | 1 |

$B_{SER-RiB}$

| | NP-NC | LP-HC | J | p |
|---|---|---|---|---|
| PC | 53 | 50 | 0·54 | 0·36 |
| PC-O | 20 | 20 | 0·67 | 0·48 |
| PI | 15 | 13 | 0·75 | 0·47 |
| SM | 14 | 15 | 0·81 | 0·47 |
| PE | 9 | 9 | 0·64 | 0·40 |
| PE-O | 8 | 11 | 0·72 | 1 |
| LPC | 5 | 1 | 0·20 | 1 |
| PS | 3 | 1 | 0·33 | 1 |
| PG | 5 | 2 | 0·40 | 1 |
| LPE | 1 | 0 | 0 | 1 |
| LPI | 1 | 0 | 0 | 1 |
| PA | 1 | 3 | 0·33 | 1 |
| Cer | 0 | 1 | 0 | 1 |

$U_{Ceb}$

| | NP-NC | LP-HC | J | p |
|---|---|---|---|---|
| PC | 2 | 4 | 0·20 | 0·22 |
| PC-O | 3 | 2 | 0·25 | 0·43 |
| PE | 1 | 1 | 0 | 1 |
| PE-O | 1 | 1 | 0 | 1 |
| PI | 0 | 1 | 1 | 1 |
| SM | 1 | 1 | 0 | 1 |
| Cer | 0 | 1 | 1 | 1 |

$U_{RiB}$

| | NP-NC | LP-HC | J | p |
|---|---|---|---|---|
| PC | 4 | 2 | 0·20 | 0·22 |
| PE | 1 | 1 | 1 | 1 |
| PI | 1 | 0 | 0 | 1 |

$B_{SER-HEA}$

| | NP-NC | LP-HC | J | p |
|---|---|---|---|---|
| PC | 56 | 51 | 0·83 | 0·48 |
| PC-O | 17 | 14 | 0·72 | 0·47 |
| LPC | 15 | 15 | 1 | 1 |
| PI | 13 | 14 | 0·87 | 1 |
| SM | 13 | 13 | 1 | 1 |
| PE | 8 | 5 | 0·63 | 1 |
| PE-O | 7 | 8 | 0·88 | 1 |
| PG | 4 | 4 | 1 | 1 |
| LPE | 2 | 2 | 1 | 1 |
| PS | 2 | 2 | 0·86 | 0·42 |
| Cer | 1 | 2 | 0·50 | 1 |
| PA | 1 | 3 | 0·33 | 1 |

$U_{HEA}$

| | NP-NC | LP-HC | J | p |
|---|---|---|---|---|
| PC | 6 | 3 | 0·29 | 0·25 |
| PI | 4 | 4 | 0·60 | 0·43 |
| PI-O | 3 | 1 | 0·33 | 1 |
| PE | 2 | 0 | 0 | 1 |
| PE-O | 1 | 1 | 1 | 1 |
| PG | 2 | 1 | 0·50 | 1 |
| SM | 1 | 1 | 1 | 1 |
| PA | 0 | 1 | 0 | 1 |
| PC-O | 0 | 1 | 0 | 1 |

$B_{SER-ADI}$

| | NP-NC | LP-HC | J | p |
|---|---|---|---|---|
| LPC | 5 | 3 | 0·60 | 0·32 |
| PC | 4 | 1 | 0 | 0·07 |
| PE-O | 3 | 1 | 0·33 | 1 |
| LPE | 2 | 1 | 0 | 1 |
| PC-O | 2 | 1 | 0·50 | 1 |
| PE | 2 | 1 | 0·50 | 1 |
| PI | 2 | 3 | 0·67 | 1 |
| PS | 2 | 1 | 0·50 | 1 |
| SM | 1 | 1 | 1 | 1 |
| PA | 0 | 2 | 0 | 1 |

$U_{ADI}$

| | NP-NC | LP-HC | J | p |
|---|---|---|---|---|
| PI | 8 | 6 | 0·75 | 1 |
| PC-O | 2 | 0 | 0 | 1 |
| PC | 1 | 1 | 1 | 1 |
| LPC | 1 | 1 | 1 | 1 |
| PG | 1 | 1 | 1 | 1 |
| PS | 1 | 0 | 0 | 1 |

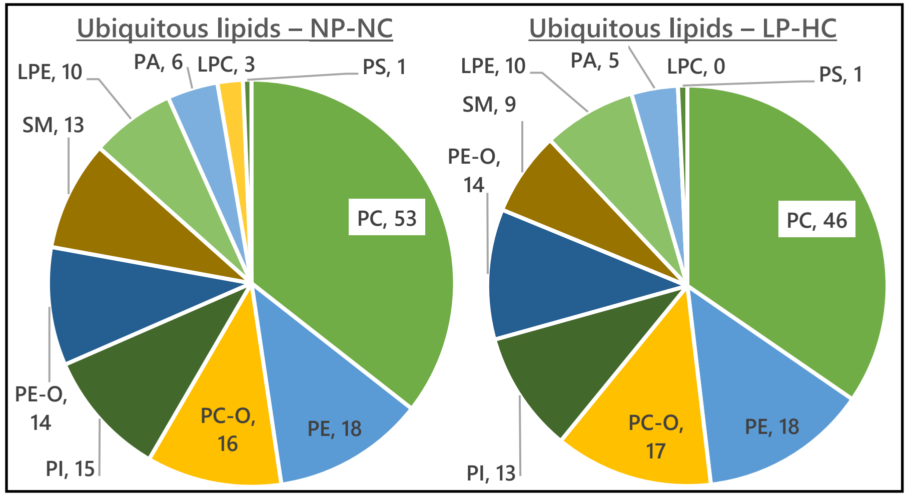


| | J | p |
|---|---|---|
| PC | 0·87 | 0·04 |
| PE | 1 | 1 |
| PC-O | 0·94 | 0·03 |
| PI | 0·87 | 1 |
| PE-O | 1 | 1 |
| SM | 0·69 | 1 |
| LPE | 0·82 | 0·42 |
| PA | 0·83 | 0·09 |
| PG | 1 | 1 |
| LPC | 0 | 1 |
| PS | 1 | 1 |

# F2N -ve

#### Liver

$U_{LIV}$

| | NP-NC | LP-HC | J | p |
|---|---|---|---|---|
| Cer | 6 | 5 | 0·83 | 1 |
| PC | 4 | 5 | 0·13 | 0·09 |
| PA | 2 | 2 | 1 | 1 |
| PG | 0 | 1 | 0 | 1 |
| PS | 2 | 1 | 0·50 | 1 |
| PI | 2 | 1 | 0 | 0·14 |

$B_{LIV-SER}$

| | NP-NC | LP-HC | J | p |
|---|---|---|---|---|
| PC | 23 | 16 | 0·70 | 1 |
| LPC | 19 | 18 | 0·85 | 0·47 |
| PI | 7 | 4 | 0·57 | 1 |
| SM | 5 | 4 | 0·80 | 1 |
| LPE | 3 | 3 | 0·75 | 0·16 |
| PC-O | 3 | 2 | 0·67 | 1 |
| PE | 3 | 3 | 1 | 1 |
| PG | 3 | 3 | 1 | 1 |
| PA | 2 | 3 | 0·67 | 1 |
| PS | 2 | 2 | 1 | 1 |
| Cer | 1 | 1 | 1 | 1 |
| PE-O | 1 | 0 | 0 | 1 |
| LPI | 0 | 1 | 0 | 1 |

#### Serum

$U_{SER}$

| | NP-NC | LP-HC | J | p |
|---|---|---|---|---|
| PC | 1 | 1 | 0 | 0·25 |
| PC-O | 3 | 3 | 0·50 | 0·43 |
| PS | 3 | 4 | 0·75 | 1 |
| LPC | 2 | 3 | 0·67 | 1 |
| SM | 1 | 1 | 1 | 1 |
| PI | 0 | 1 | 0 | 1 |

$B_{SER-CEB}$

| | NP-NC | LP-HC | J | p |
|---|---|---|---|---|
| PC | 13 | 8 | 0·31 | 0·08 |
| PC-O | 13 | 9 | 0·68 | 1 |
| PI | 7 | 5 | 0·71 | 1 |
| LPC | 6 | 8 | 0·75 | 1 |
| SM | 6 | 5 | 0·83 | 1 |
| PE | 4 | 4 | 1 | 1 |
| PE-O | 3 | 1 | 0·33 | 1 |
| PS | 2 | 2 | 1 | 1 |
| LPE | 1 | 2 | 0·5 | 1 |
| LPI | 1 | 1 | 1 | 1 |
| PA | 0 | 1 | 0 | 1 |
| Cer | 1 | 1 | 0 | 1 |

$B_{SER-RiB}$

| | NP-NC | LP-HC | J | p |
|---|---|---|---|---|
| PC | 20 | 15 | 0·67 | 0·47 |
| LPC | 11 | 7 | 0·64 | 1 |
| PE | 10 | 10 | 1 | 1 |
| PC-O | 9 | 6 | 0·67 | 1 |
| PI | 8 | 5 | 0·63 | 1 |
| SM | 5 | 6 | 0·57 | 0·44 |
| PS | 3 | 3 | 1 | 1 |
| PE-O | 3 | 1 | 0·33 | 1 |
| LPE | 2 | 2 | 1 | 1 |
| PA | 2 | 3 | 0·67 | 1 |
| LPI | 1 | 1 | 1 | 1 |
| Cer | 1 | 1 | 0 | 1 |

$B_{SER-HEA}$

| | NP-NC | LP-HC | J | p |
|---|---|---|---|---|
| PC | 12 | 4 | 0·14 | 0·04 |
| LPE | 11 | 10 | 0·91 | 1 |
| LPC | 10 | 6 | 0·45 | 0·38 |
| PI | 4 | 0 | 0 | 0 |
| SM | 4 | 2 | 0 | 0·06 |
| PA | 2 | 1 | 0 | 0·15 |
| PC-O | 1 | 3 | 0·33 | 1 |
| PG | 1 | 1 | 0·25 | 0 |
| PS | 1 | 1 | 1 | 1 |
| Cer | 1 | 0 | 0 | 1 |

#### Cerebellum

$U_{CEB}$

| | NP-NC | LP-HC | J | p |
|---|---|---|---|---|
| PC | 8 | 6 | 0·75 | 1 |
| PC-O | 5 | 7 | 0·71 | 1 |
| PE | 3 | 3 | 1 | 1 |
| PE-O | 1 | 2 | 0·50 | 1 |
| SM | 1 | 2 | 0·50 | 1 |
| PG | 1 | 0 | 0 | 1 |
| PI-O | 1 | 1 | 1 | 1 |
| Cer | 0 | 1 | 0 | 1 |

#### Right brain

$U_{RiB}$

| | NP-NC | LP-HC |
|---|---|---|
| - | - | - |

#### Heart

$U_{HEA}$

| | NP-NC | LP-HC | J | p |
|---|---|---|---|---|
| PC | 4 | 6 | 0·43 | 0·39 |
| PC-O | 1 | 2 | 0·50 | 1 |
| PE | 0 | 2 | 0 | 1 |
| PI | 8 | 8 | 1 | 1 |
| PS | 1 | 0 | 0 | 1 |
