## Supplementary material for "Revealing grand-paternal programming of lipid metabolism using a novel computational tool": Watkins proj_31P data_SI_final.pdf

Supplementary data

Furse *et al.*

### Sections

- Assignments
- Samples
- Degradation test

### Assignments

| Shift (ppm) | Assignment |
|---|---|
| 0·00 | Phosphatidylcholine* |
| 0·05 | plasmalogen-Phosphatidylcholine |
| 0·18-0·22 | *Unknown* |
| 0·45 | *lyso*-Phosphatidylcholine |
| 0·52 | Phosphatidylserine |
| 0·55 | Phosphatidylethanolamine** |
| 0·58 | plasmalogen-Phosphatidylethanolamine |
| 0·78 | Cardiolipin |
| 0·83 | Sphingomyelin |
| 0·91 | *lyso*-Phosphatidylethanolamine |
| 1·08-1·12 | Phosphatidylinositol |
| 1·23 | Phosphatidylglycerol |
| 1·53 | *lyso*-Phosphatidylinositol |
| 1·72 | *lyso*-Phosphatidylglycerol |
| 4·80-5·50 | Phosphatidic acid** |
| 6·00-6·50 | *lyso*-Phosphatidic acid |

*PC is 0·00 ppm by definition, *i.e.* data were referenced to give the PC signal at 0·00 ppm.

** PE and PA have several resonances, due to interactions with the solvent system and several possible adducts

References

- Bosco *et al.* 1997, Anal. Biochem. DOI: 10.1006/abio.1996.9907
- Culeddu *et al.* 1998, Magn. Res. Chem. DOI: 10.1002/(sici)1097-458x(199812)36:12<907::aid-omr394>3.0.co;2-5
- Murgia, *et al.* 2003, Lipids, DOI: 10.1007/s11745-003-1500-3
- Cremonini *et al.* 2004, J. Sci. Food Agri. DOI: 10.1002/jsfa.1683
- Furse *et al.* 2013, *J. Chem Biol*, DOI: 10.1007/s12154-012-0090-1

### Samples

- Liver
  - F1N
    - NP-NC
    - LP-HC
  - F1A
    - NP-NC
    - LP-HC
  - F2N
    - NP-NC
    - LP-HC

- Heart
  - F1N
    - NP-NC
    - LP-HC
  - F1A
    - NP-NC
    - LP-HC
  - F2N
    - NP-NC
    - LP-HC

- Adipose
  - F1A
    - NP-NC
    - LP-HC
    - LP-HC (petrol washed)

- Serum
  - F2N (pooled)

- Right Brain
  - F1A (pooled)

- Cerebellum
  - F1A male
  - F2N female

### Liver
# F1N
# NP-NC

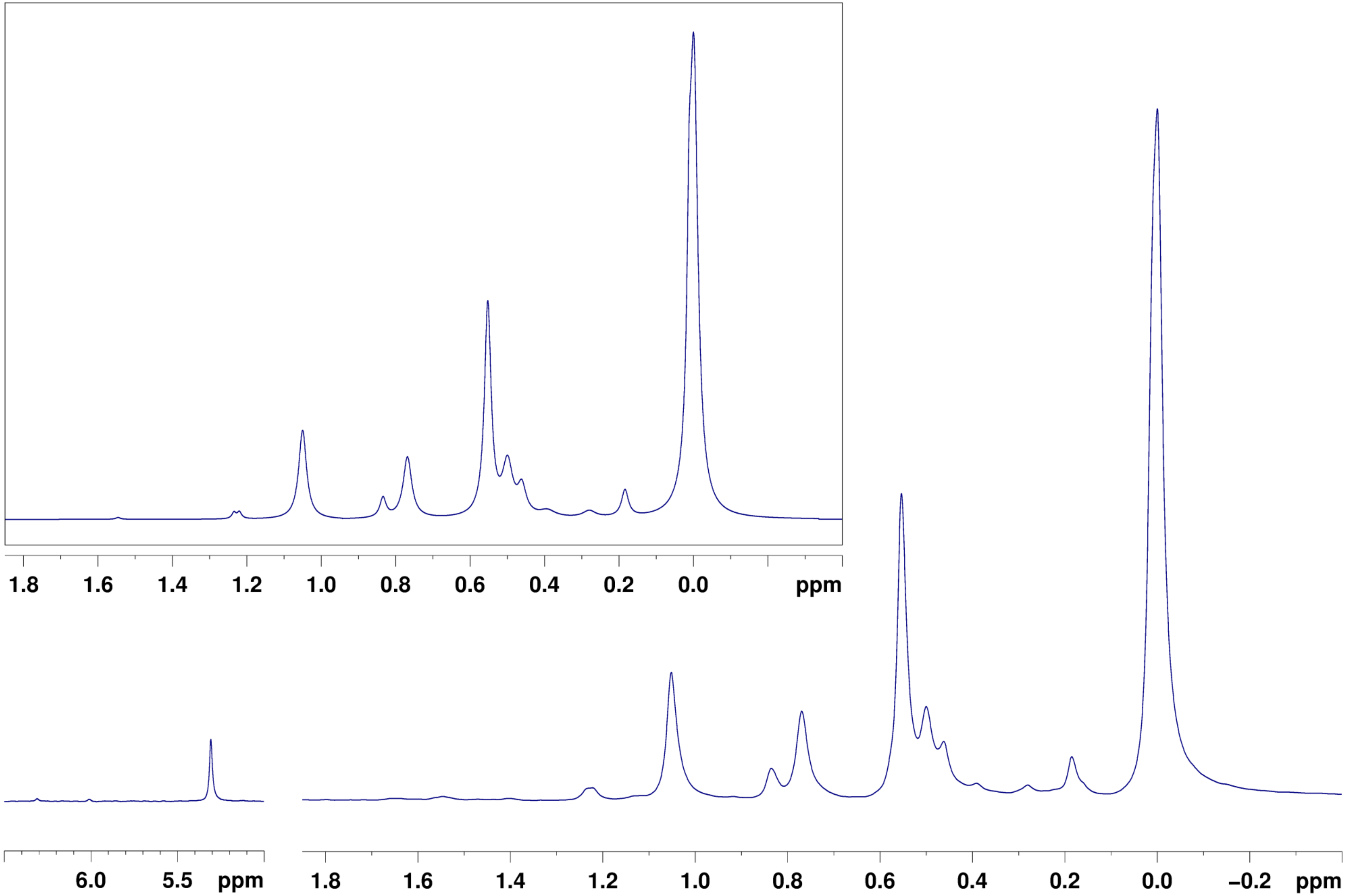

### Liver
# F1N
# LP-HC

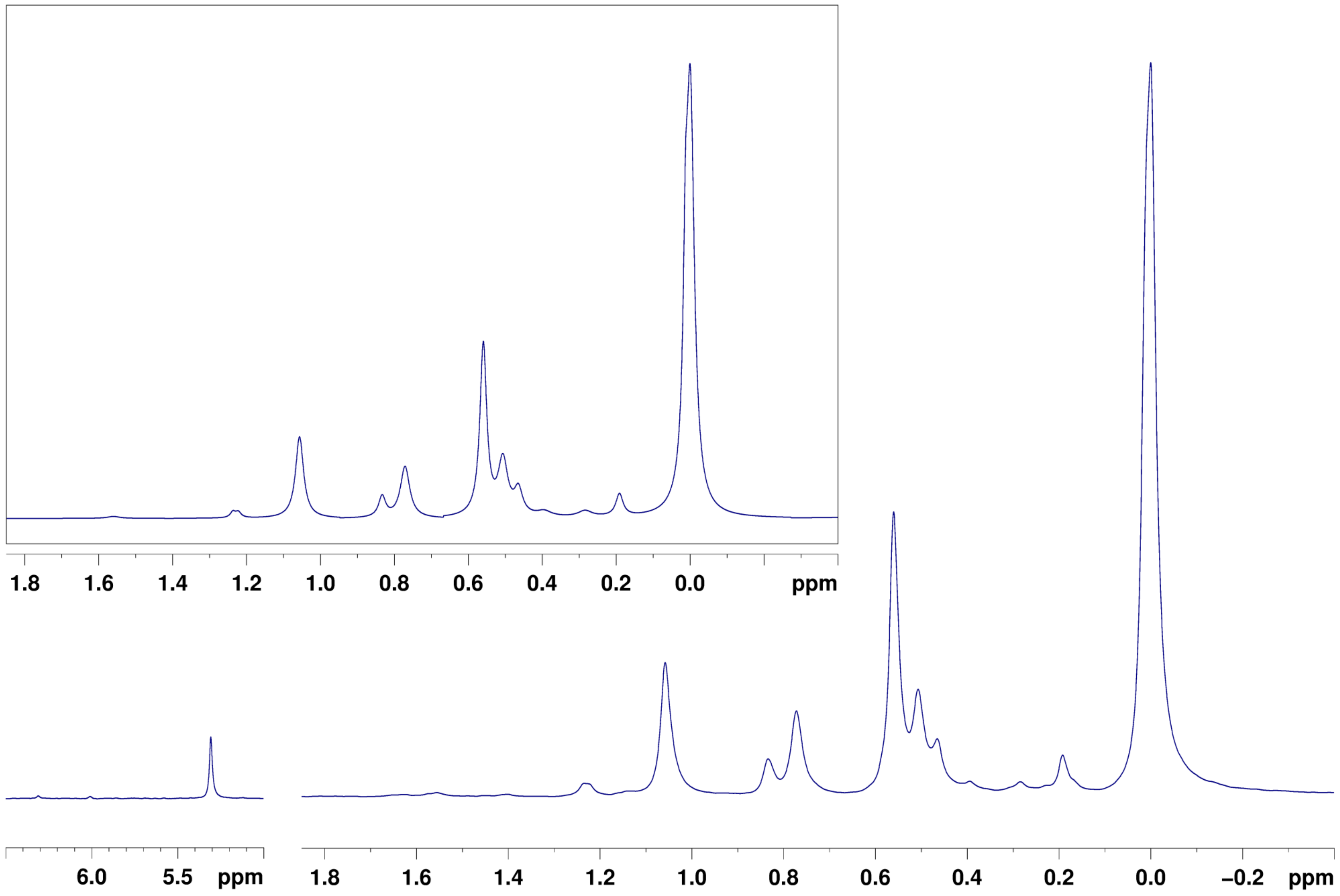

### Liver
# F1A
# NP-NC

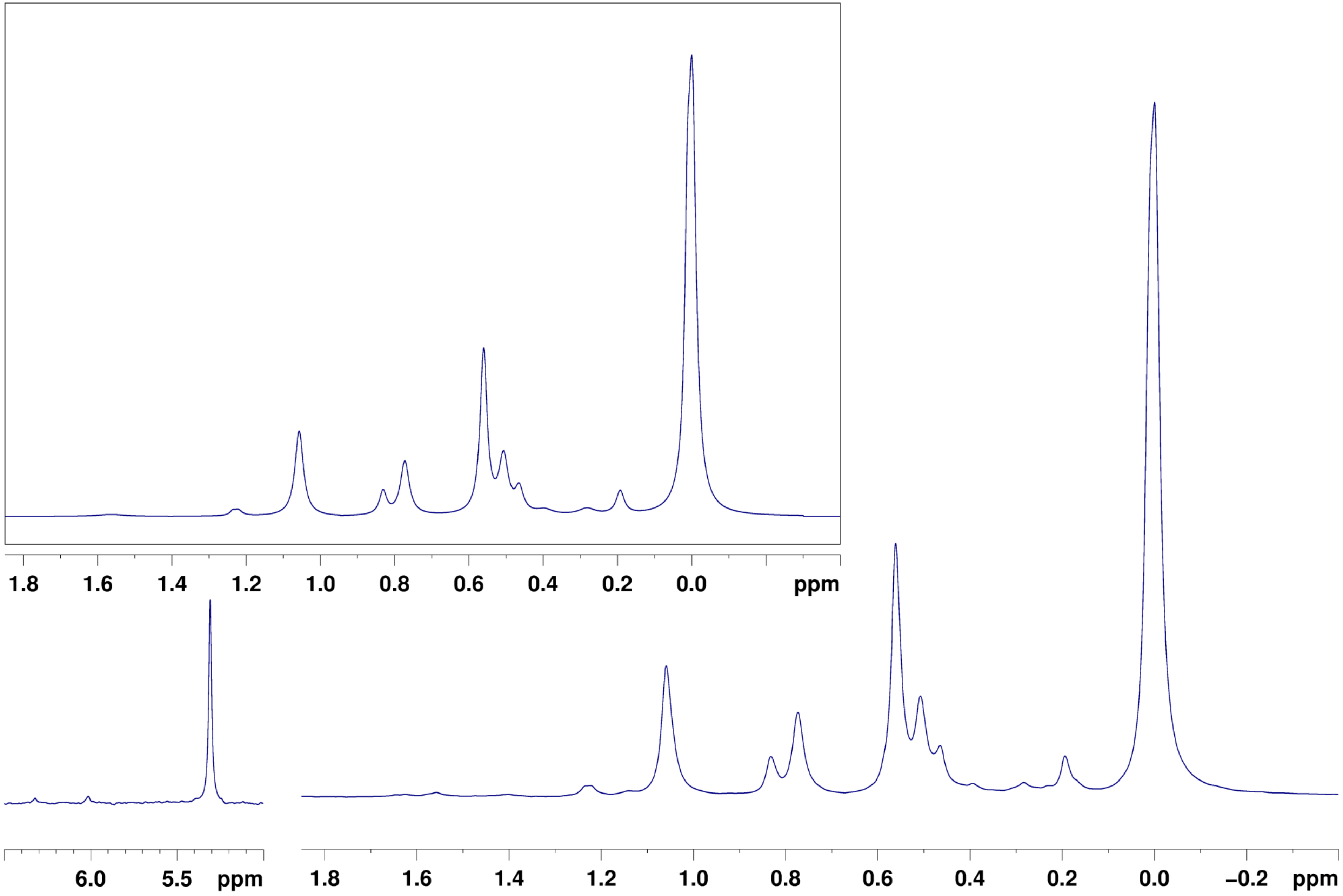

### Liver
# F1A
# LP-HC

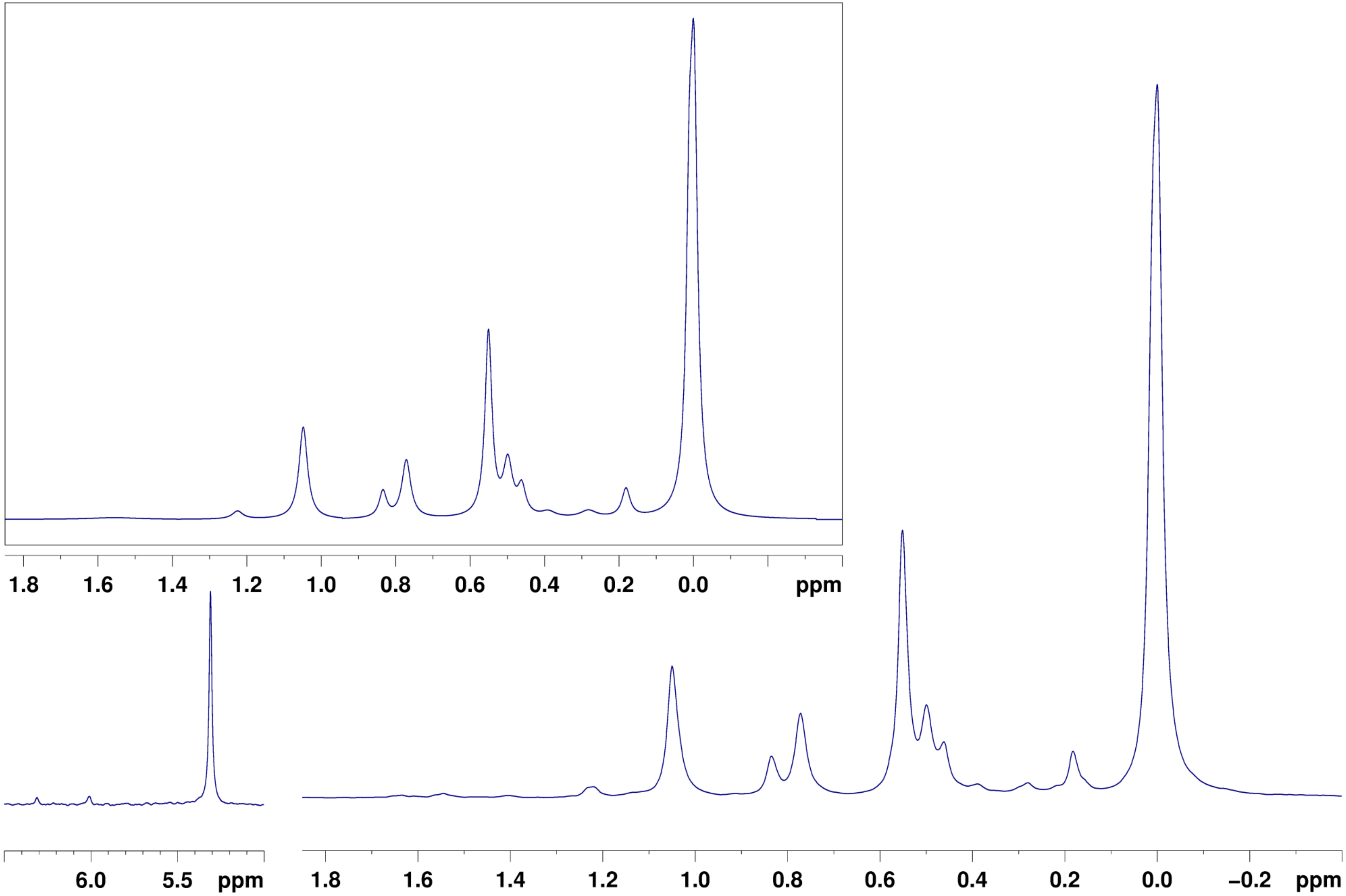

### Liver
# F2N
# NP-NC

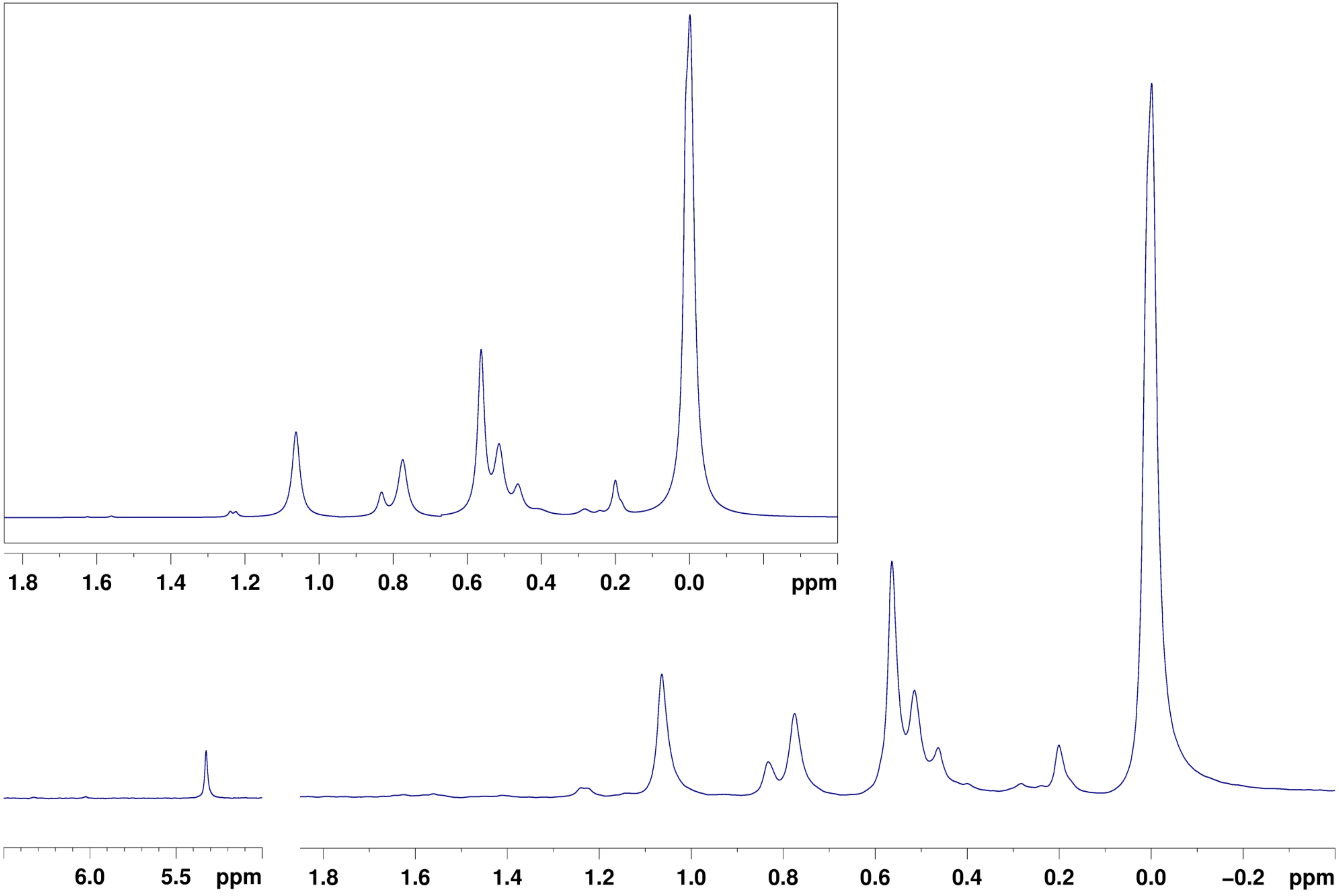

### Liver
# F2N
# LP-HC

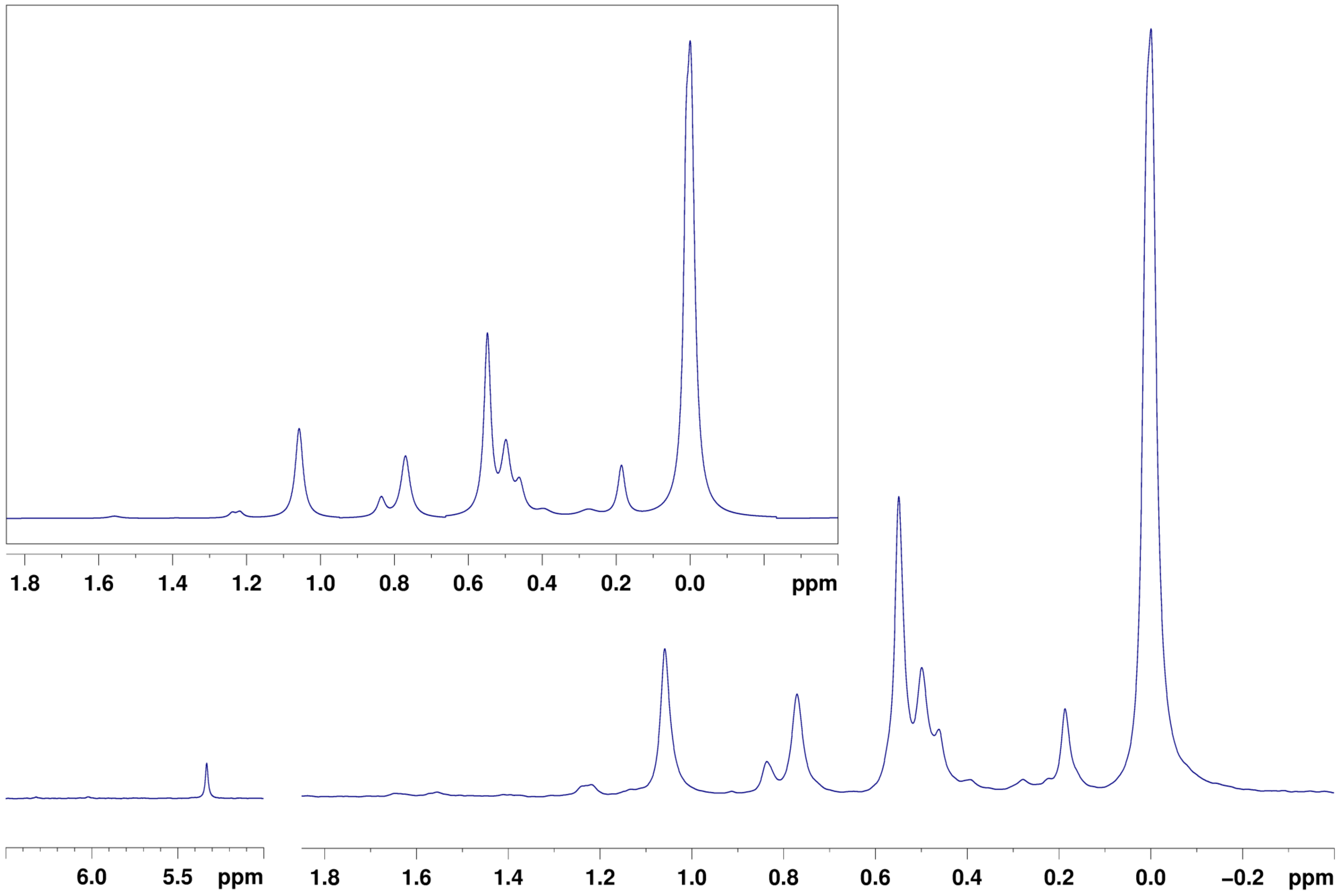

Heart
F1N
NP-NC

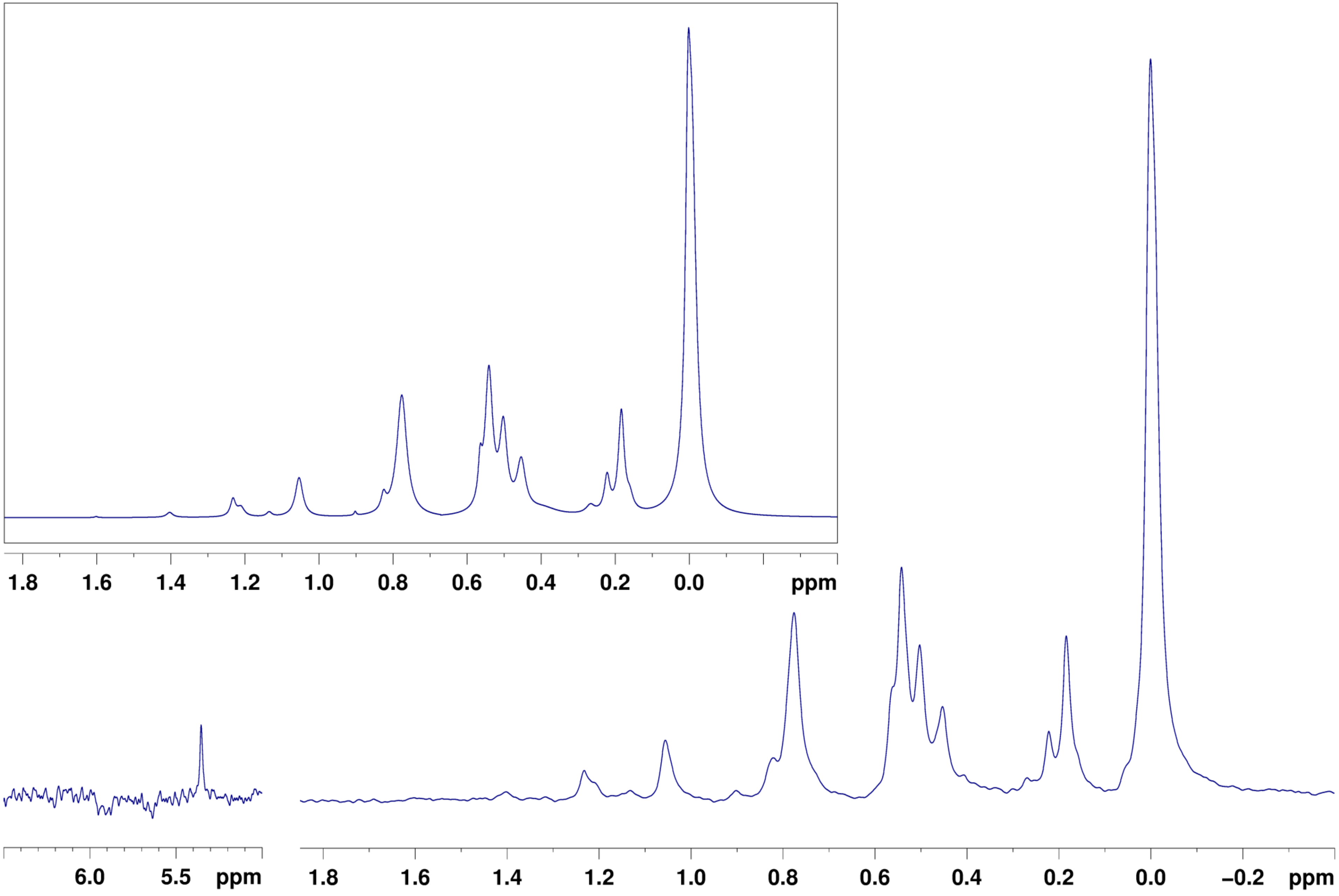

### Heart
# F1N
# LP-HC

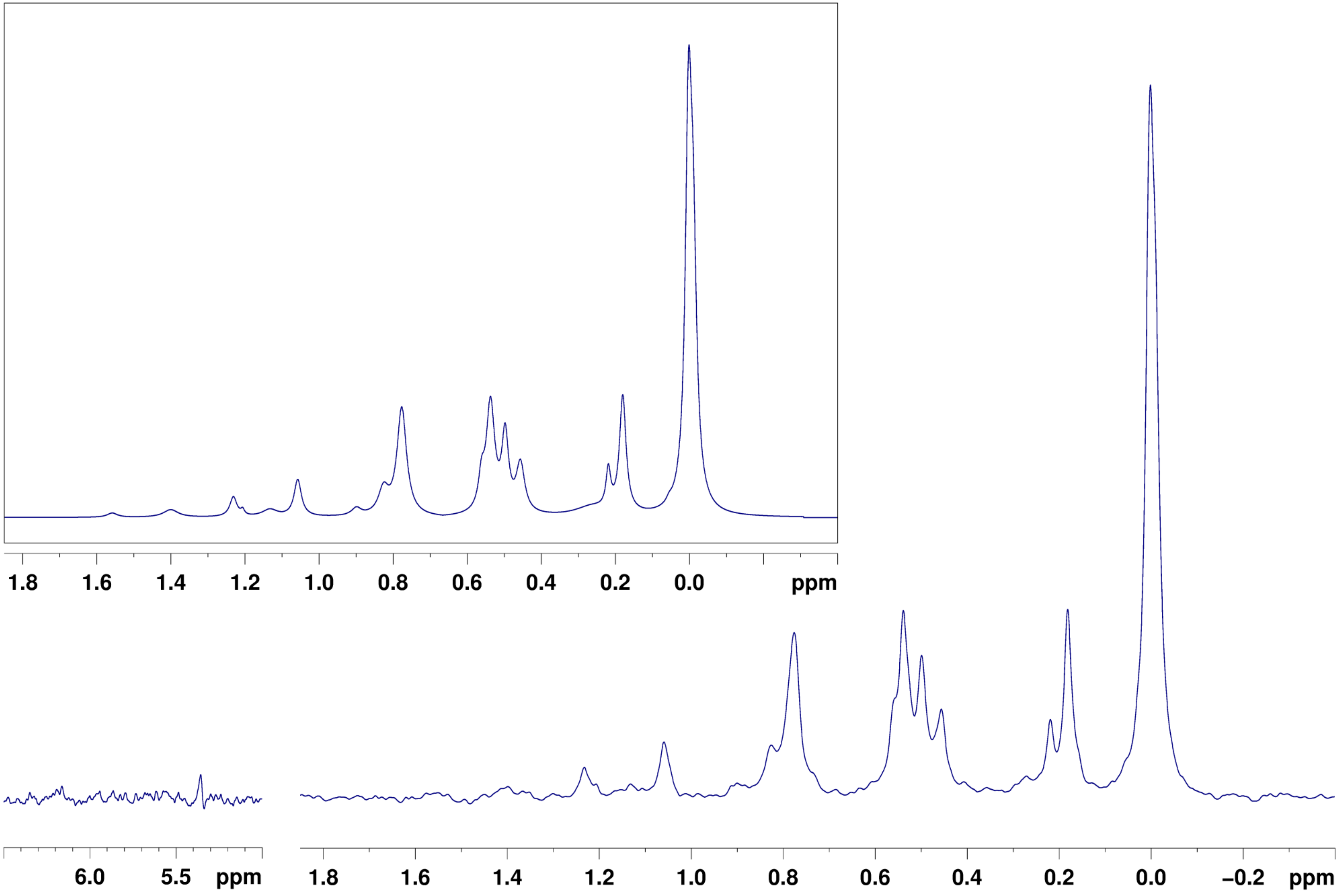

### Heart
# F1A
# NP-NC

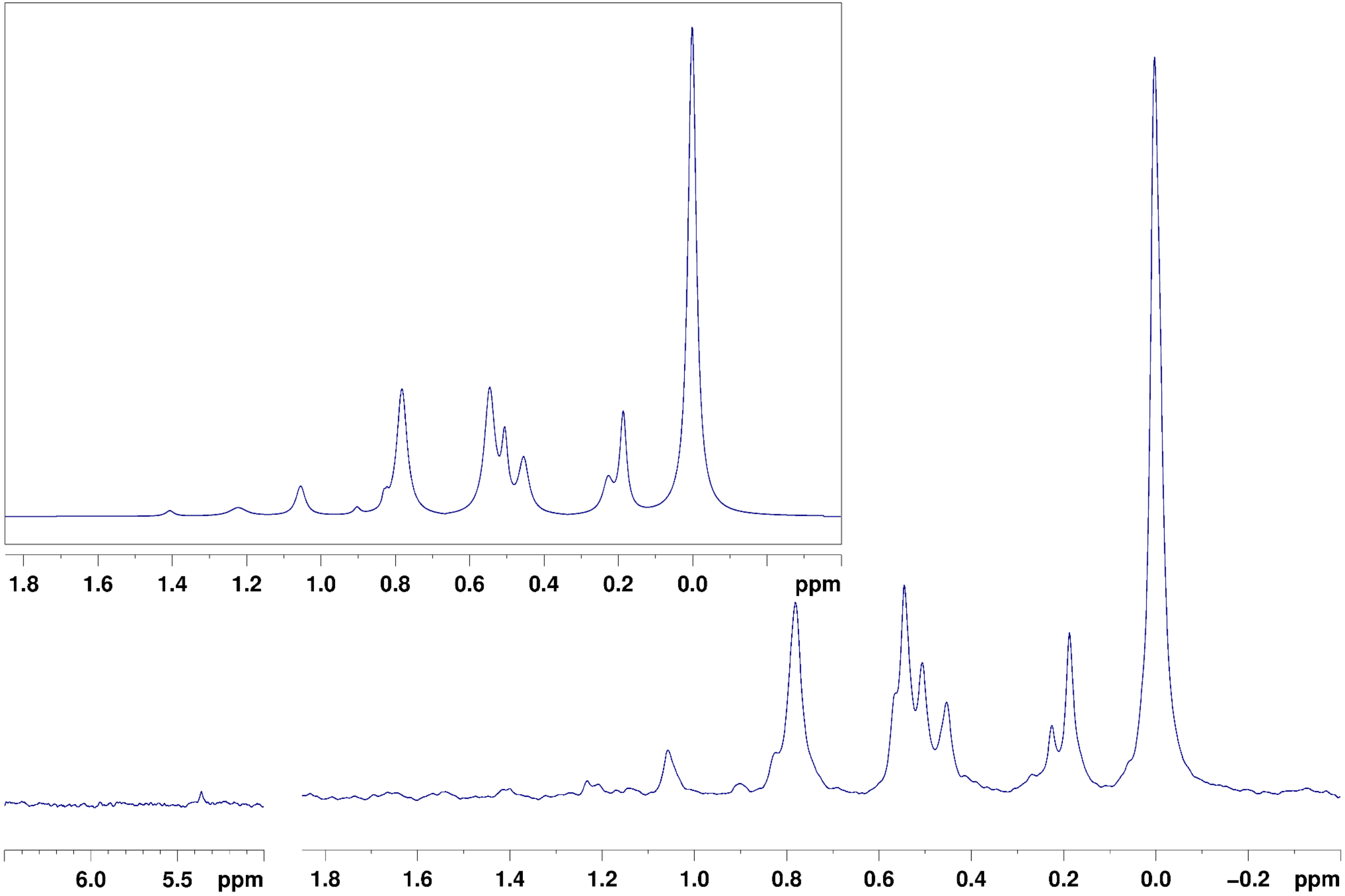

### Heart
# F1A
# LP-HC

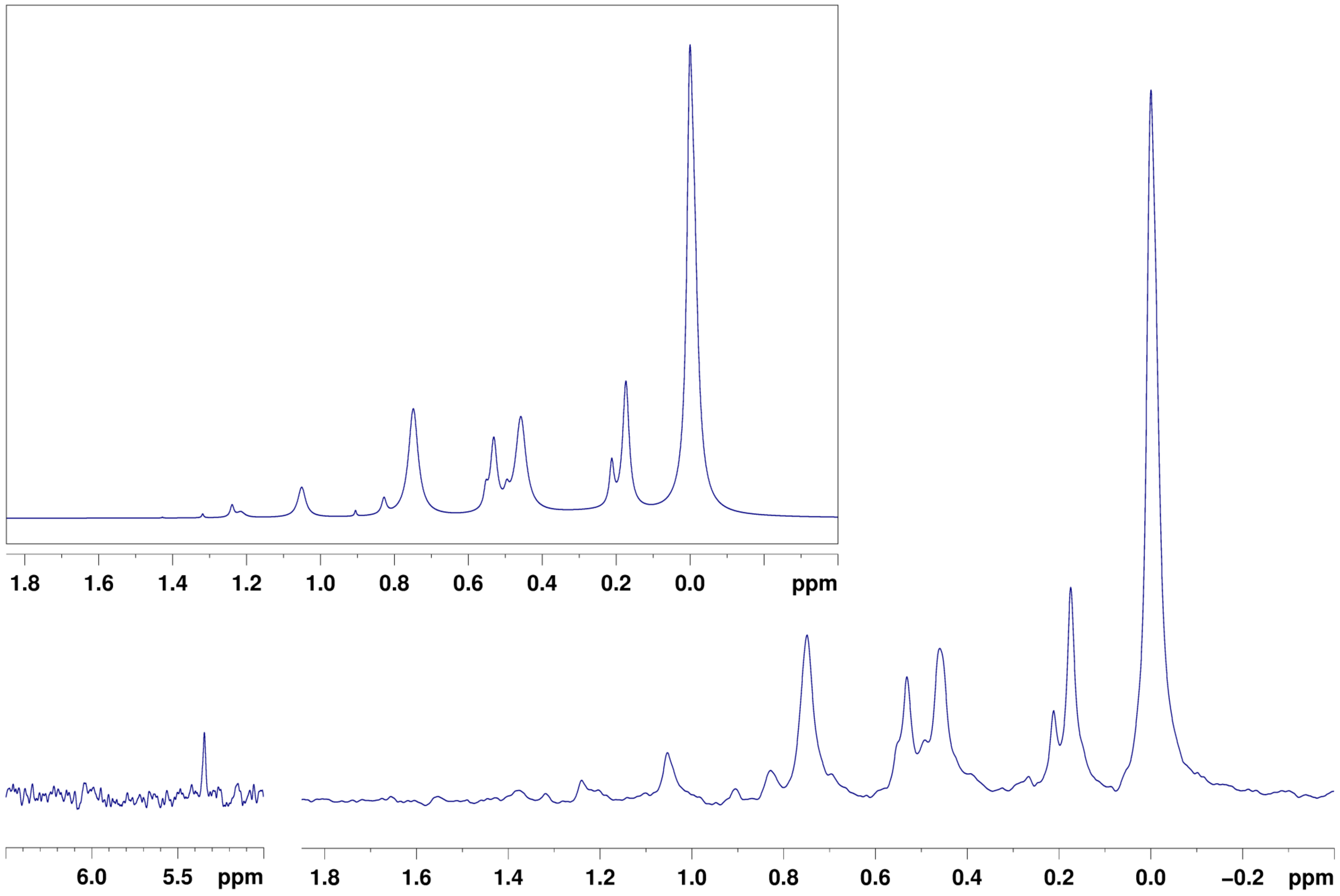

### Heart
# F2N
# NP-NC

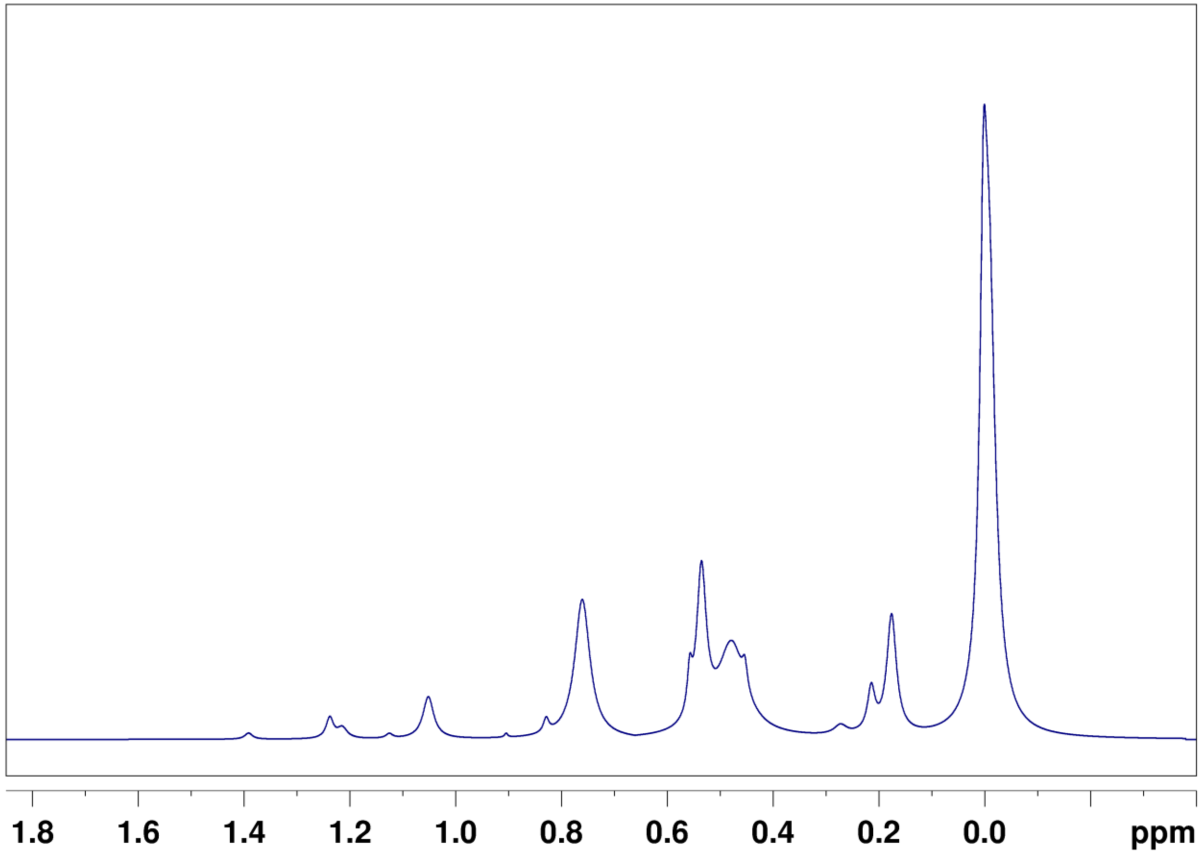


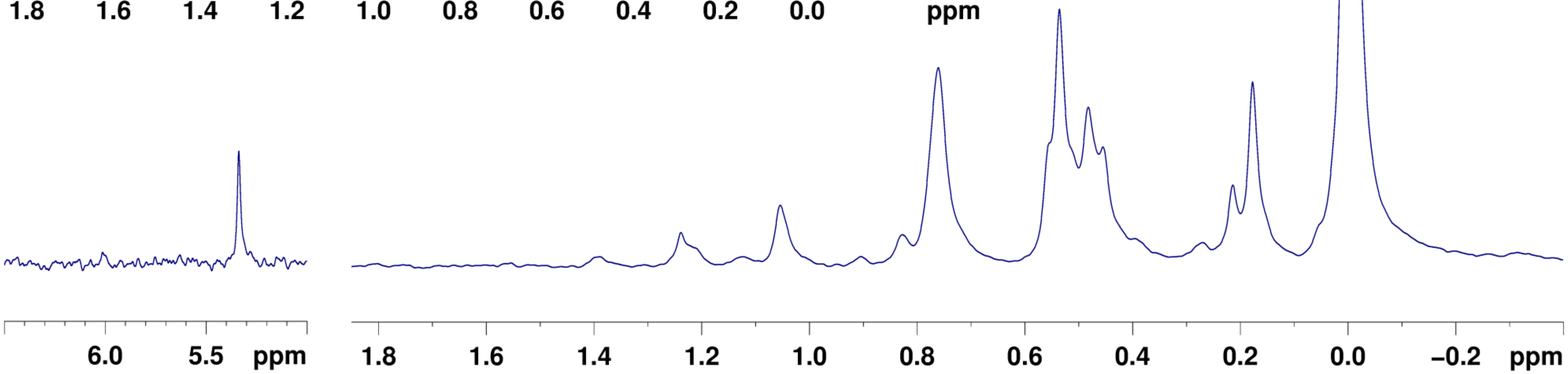

### Heart
# F2N
# LP-HC

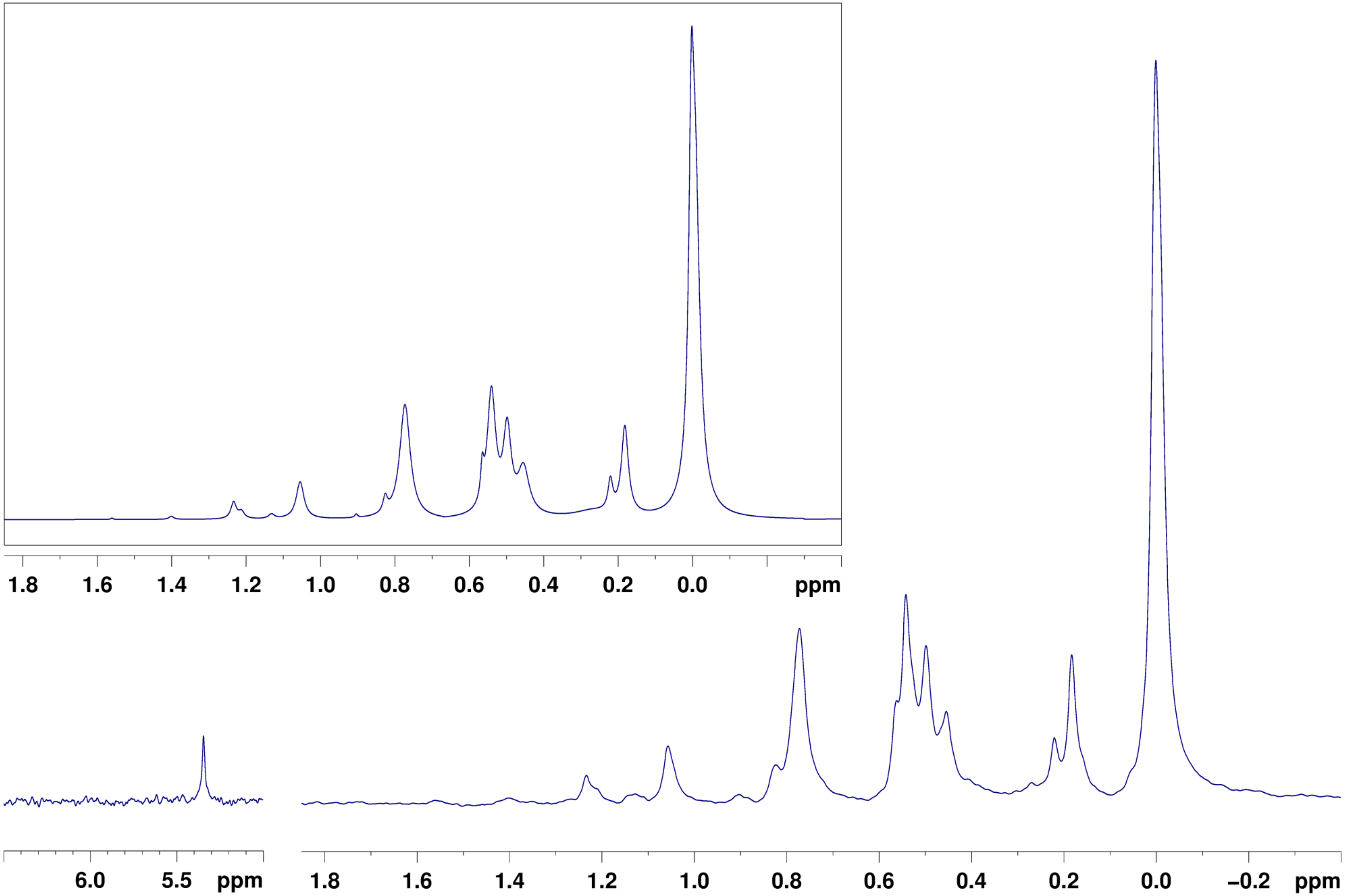

### Adipose
# F1A
# NP-NC

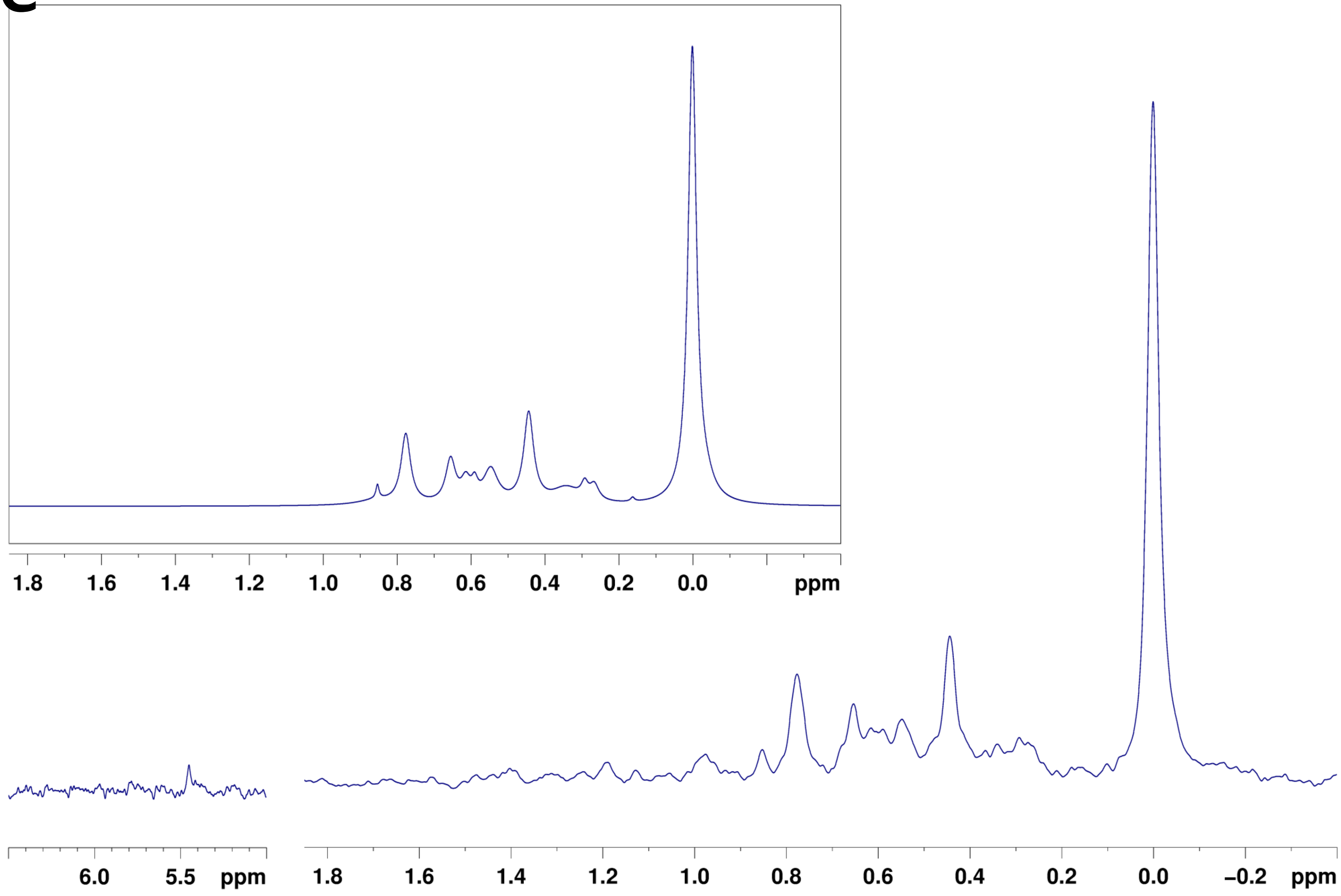

Adipose
F1A
LP-HC

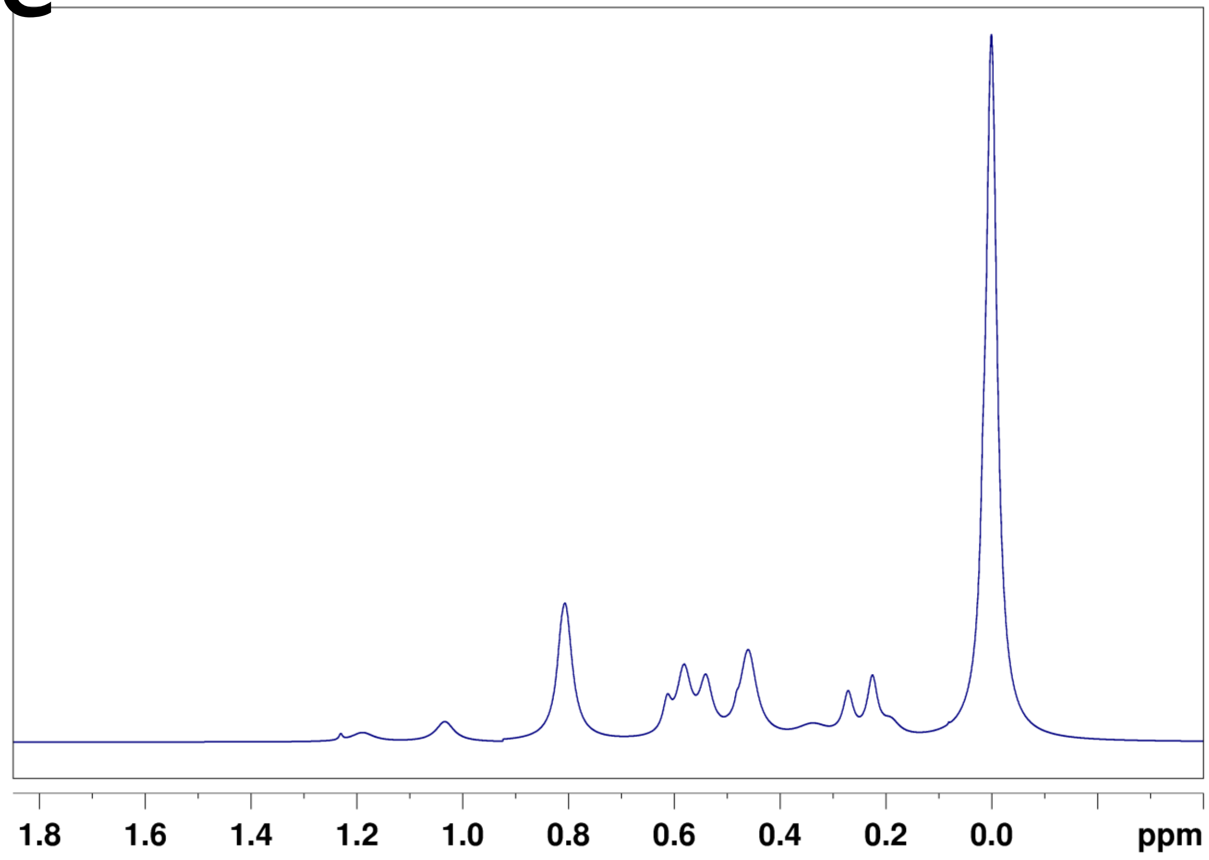


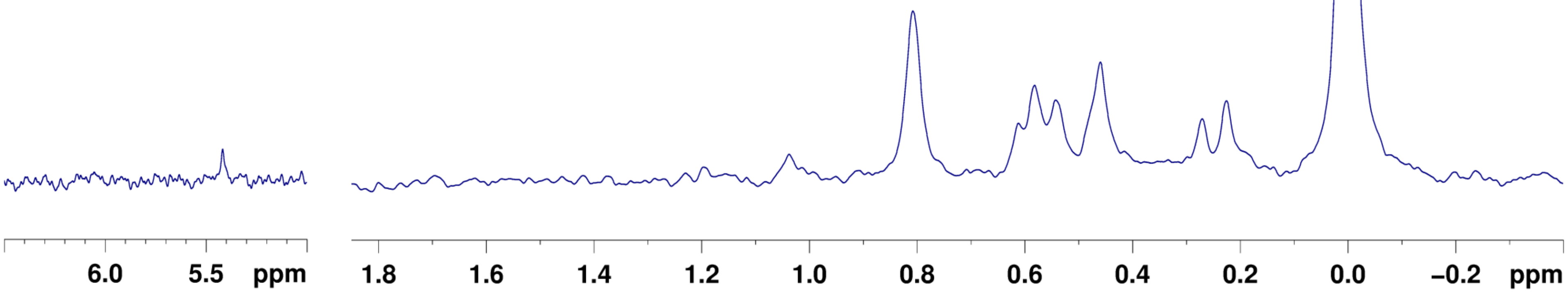

Adipose
F1A
LP-HC
PW

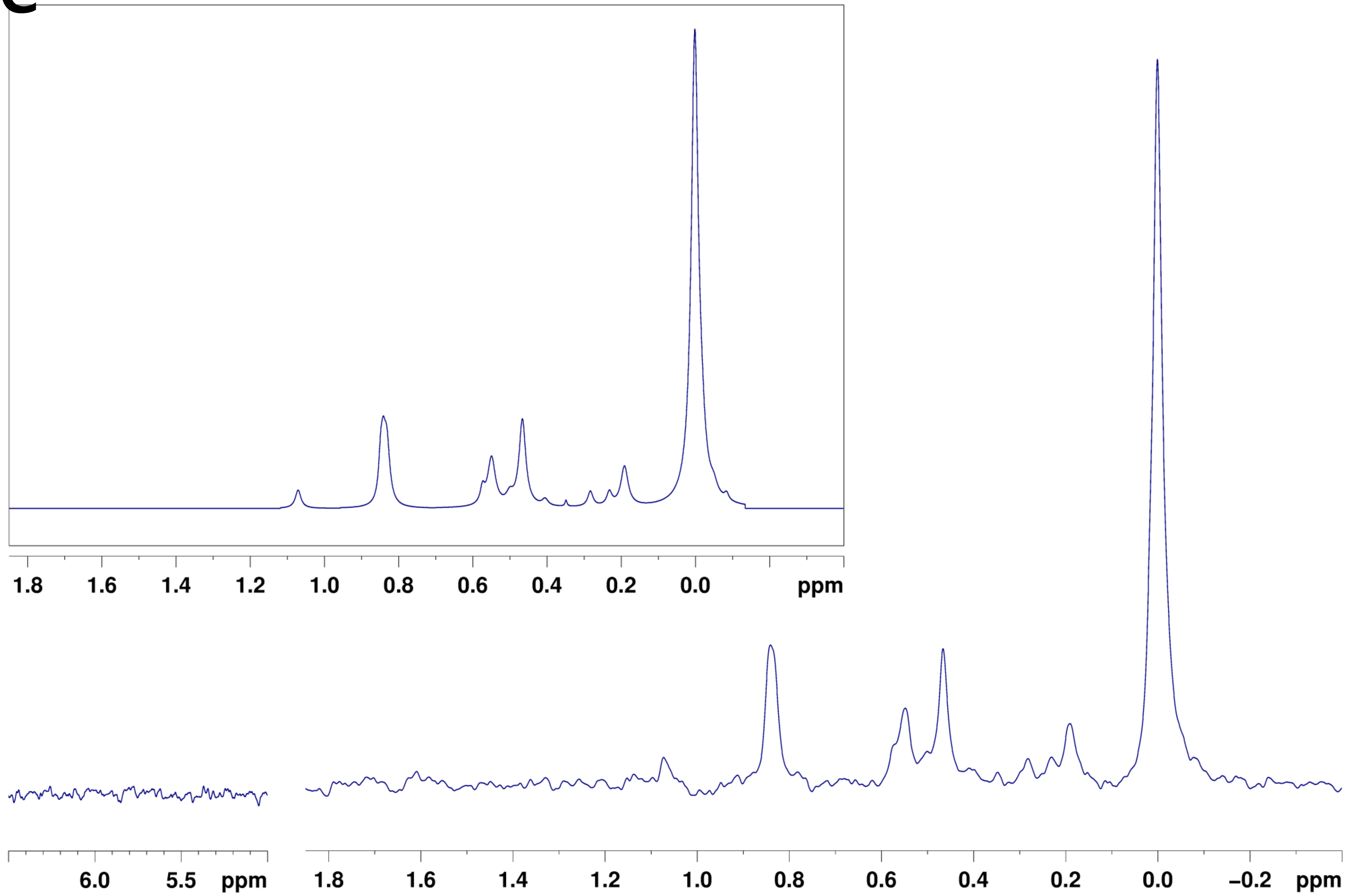

### Serum
# F2N
### Male

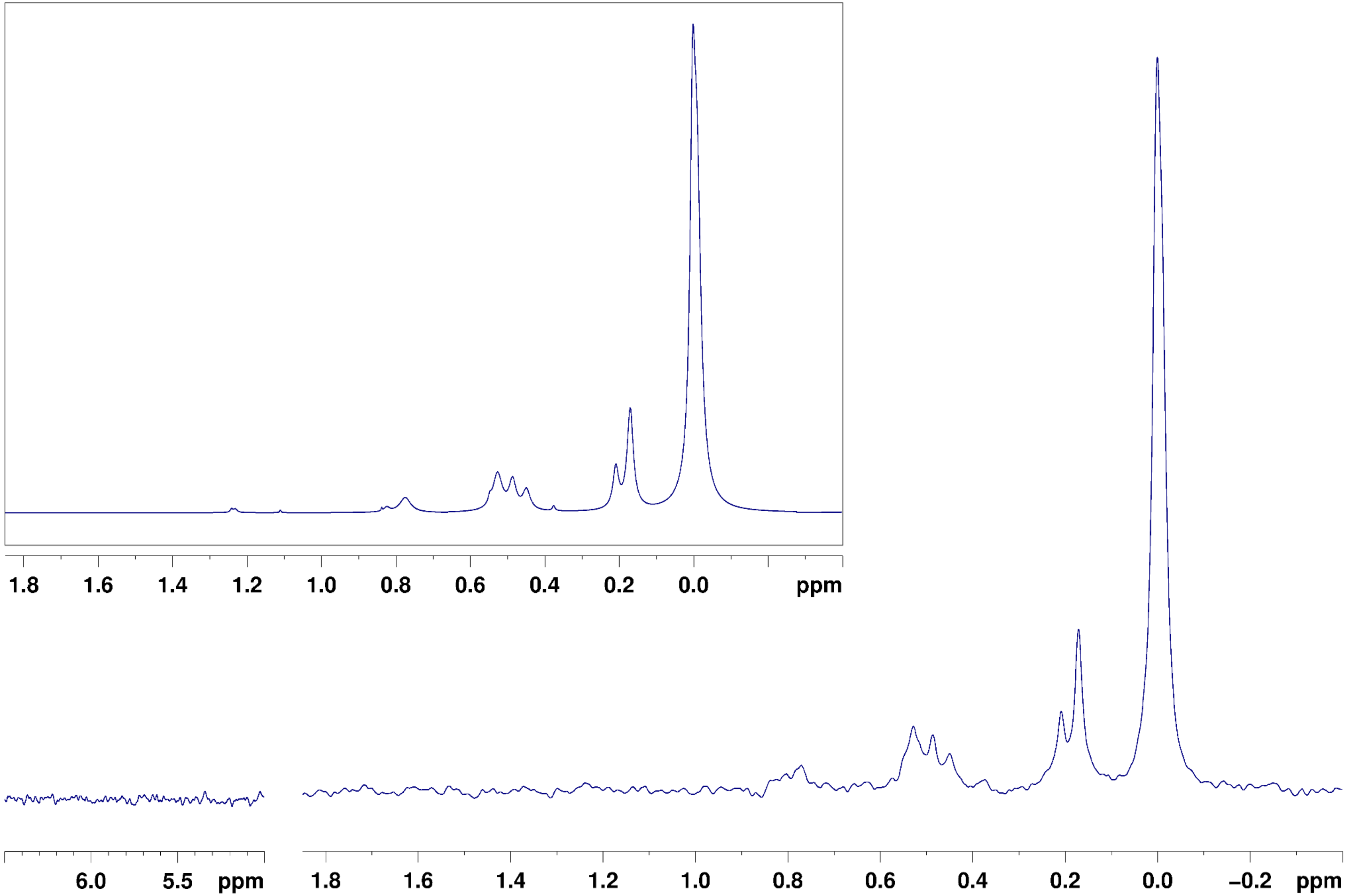

### Right Brain
# F1A
### Pooled

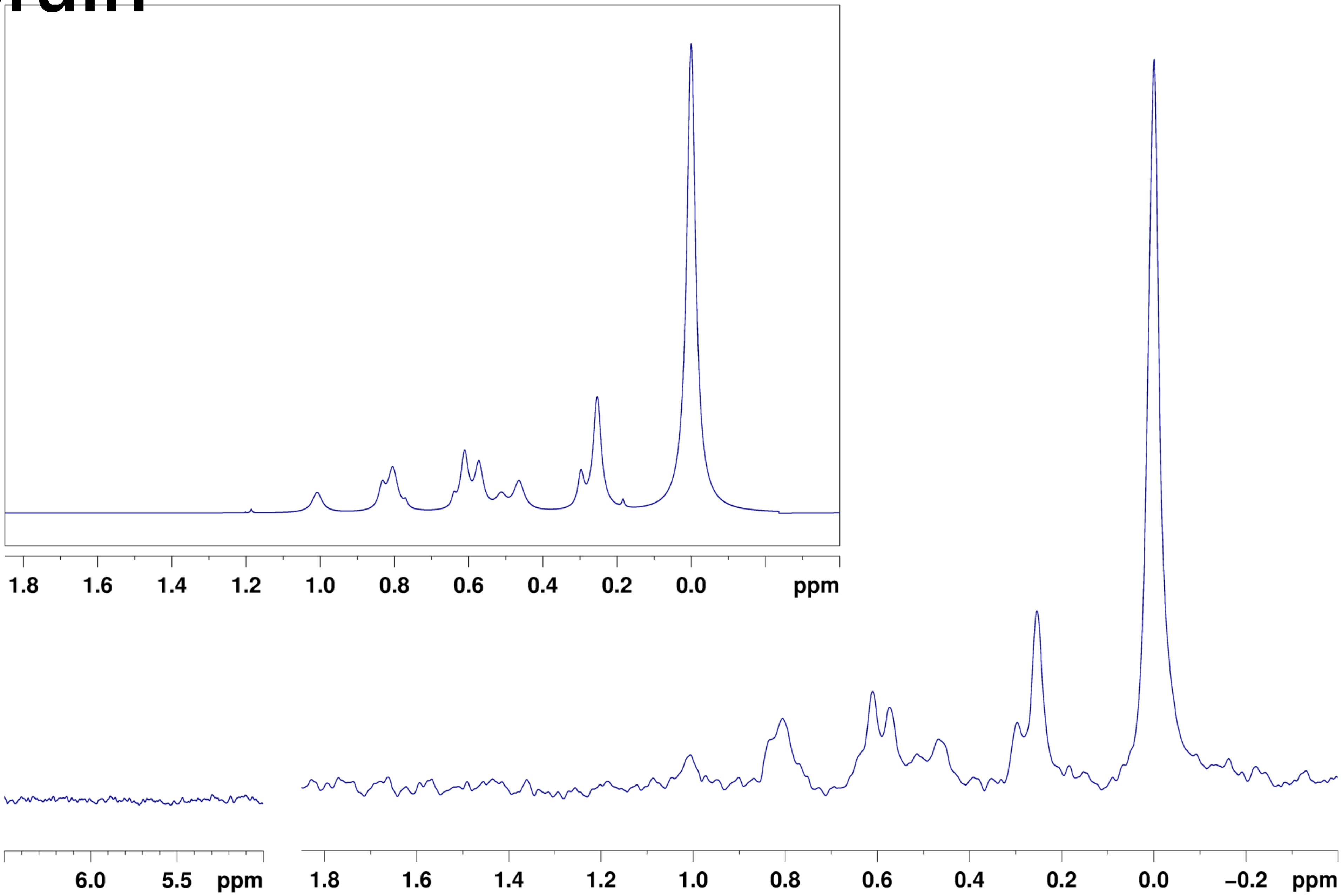

### Cerebellum
# F1A
### Male

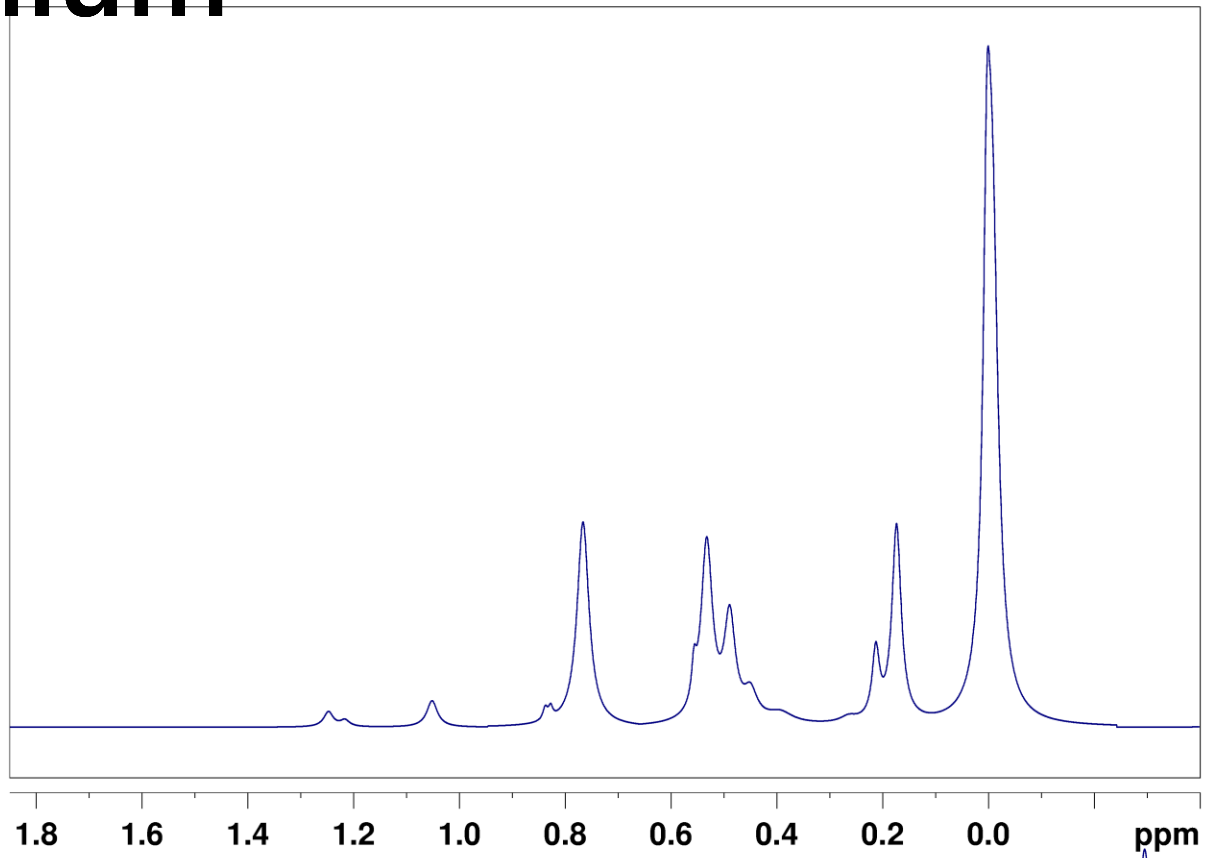


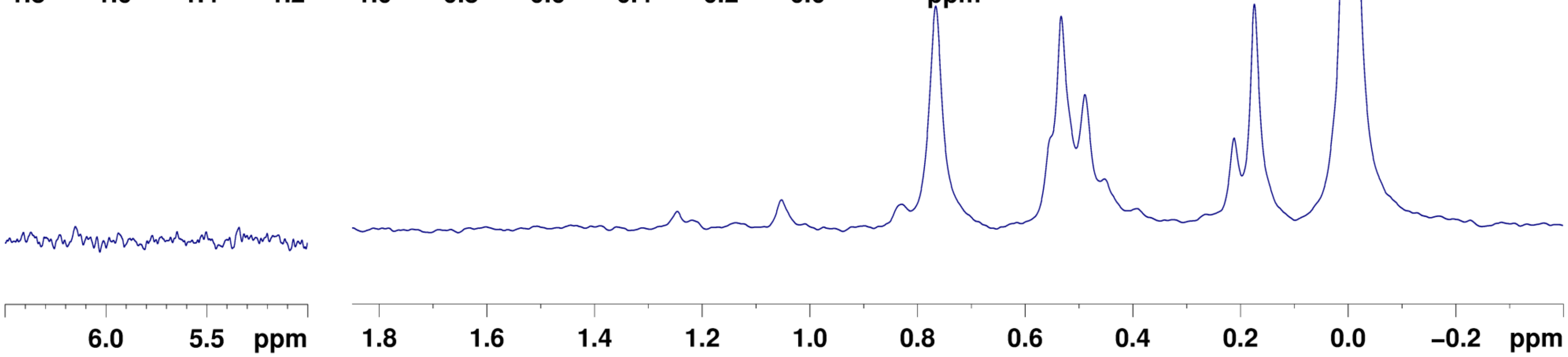

### Cerebellum
# F2N
### Female

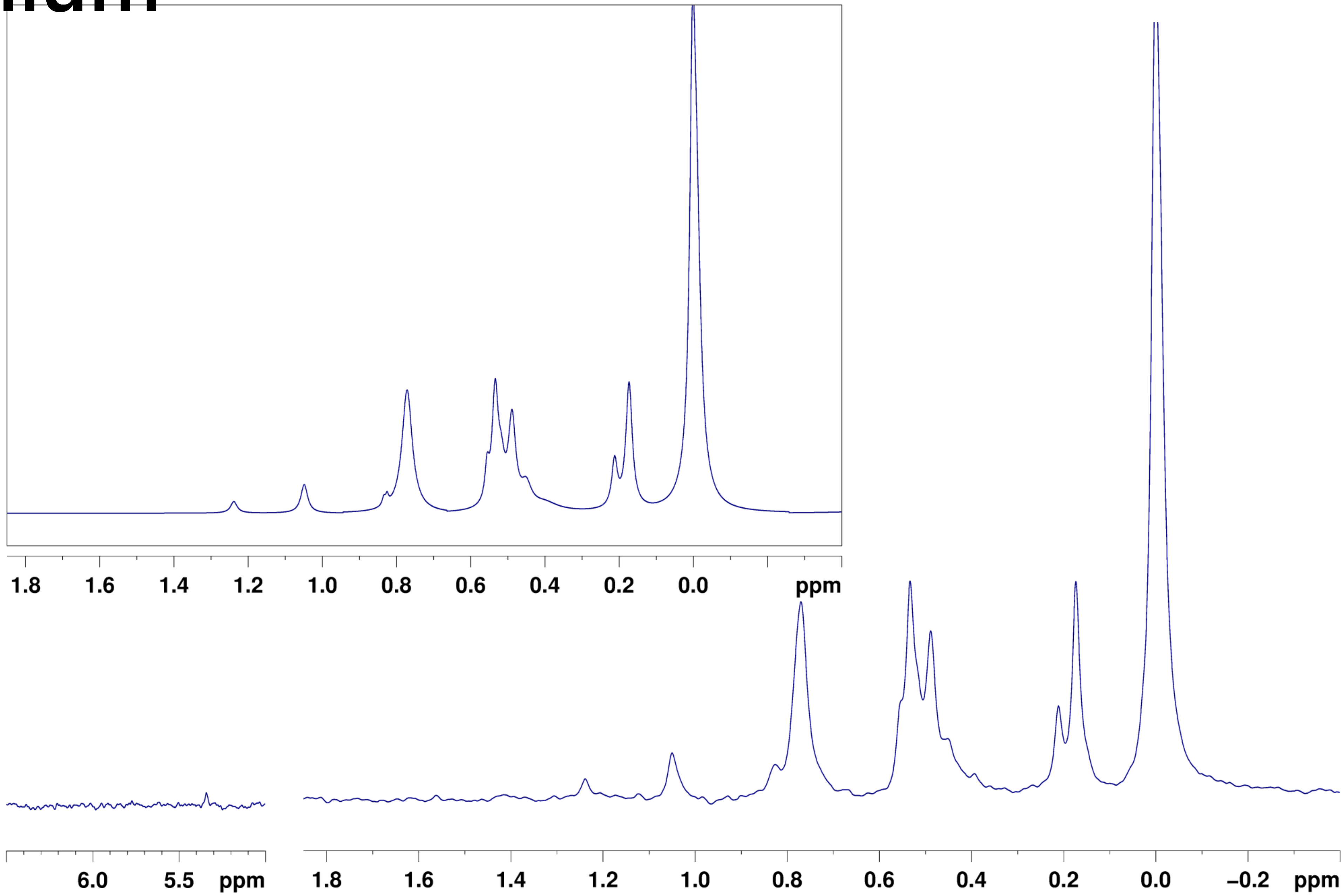

### Degradation test

- F2N, Liver, NP-NC
  - 0h (red trace)
  - +48h  (blue trace)

Liver
F2N
NP-NC
0h, +48h

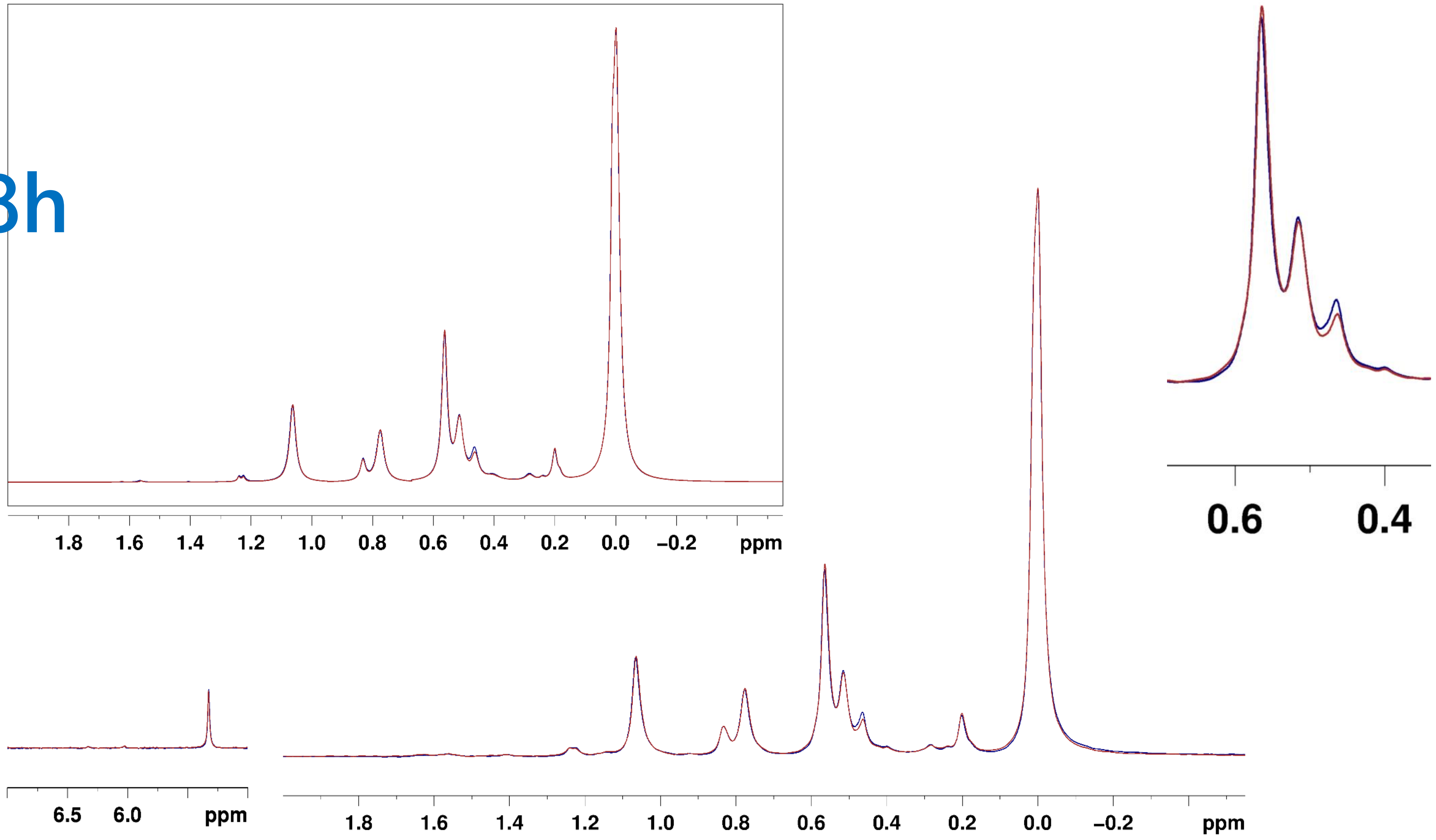
